# Dynamics of Ribosomal RNA Transcription and Abundance in Normal and Leukemic Hematopoiesis

**DOI:** 10.1101/2025.08.01.668217

**Authors:** Eleanor I. Sams, Victoria K. Feist, Erin M. Gray, Subin S. George, Charles Antony, Jill A. Henrich, Kiyan Alrobaire, Putzer J. Hung, Yulia Gonskikh, Sydney L. Moore, Dishari Ghatak, Julia Wald, Margaret C. Dunagin, Nisargbhai Shah, Jordi C.J. Hintzen, Zhengyi Wang, Noa Erlitzki, George M. Burslem, Arjun Raj, Robert L. Bowman, Kathy Fange Liu, Wei Tong, Robert A.J. Signer, Vikram R. Paralkar

## Abstract

Transcription of ribosomal RNAs (rRNAs) from rDNA repeats is a critical first step of ribosome biogenesis. Often deemed a housekeeping process, its cell-type-specific regulation is largely unexplored. We profiled nascent rRNA and mature rRNAs (ribosome subunits) across mouse hematopoiesis, and observed 10-fold, cell-type-specific variation in rRNA abundance that is largely uncoupled from proliferation and translation rates. We also tested and confirmed the longstanding assumption that acute myeloid leukemia progenitors show notably higher rRNA levels than matched normal counterparts. These trends in nascent rRNA across hematopoietic contexts parallel changes in the proportion of active rDNA repeats. Absolute quantification of rRNAs unexpectedly revealed an excess of large versus small ribosomal subunits in all cells, particularly in the myeloid lineage. Finally, using an RNA Polymerase I degron system, we found that reduced rRNA transcription promotes myeloid progenitor differentiation. Collectively, our work uncovers complex dynamics and functional significance of rRNA regulation in hematopoiesis.

## INTRODUCTION

Transcription of ribosomal RNAs (rRNAs) from rDNA repeats accounts for a large proportion of all cellular transcription, synthesizing over 80% of the total RNA in eukaryotic cells^1–4^. In mammals, rDNA repeats exist in hundreds of copies in tandem arrays across multiple chromosomes, and in any given cell only a subset are active and transcribed into 47S pre-rRNA by RNA Polymerase I (Pol I)^5–7^. Over the course of two hours^8^, 47S pre-rRNA is processed into mature 5.8S, 18S, and 28S rRNAs and assembled with 5S rRNA (transcribed by RNA Pol III) and nearly 80 ribosomal proteins to form the 40S and 60S ribosomal subunits^6,9,10^. Together, these four mature rRNAs form the catalytic core and account for the majority of the mass of the ribosome^1,3^. Despite its central role, there is limited characterization of how rRNA transcription is regulated in complex organ systems, in part due to two longstanding limitations. First, rRNA transcription and ribosome biogenesis are often considered uniform housekeeping processes with limited cell-to-cell variations or contributions to cell identity and behavior. Second, the repetitive nature of rDNA presents major technical challenges for detailed mechanistic and bioinformatic analyses^1^. As a result of these challenges, rRNA transcription is often neglected in studies of transcriptional regulation in physiological and pathological contexts.

Hematopoiesis provides an ideal model system to study cell-type-specific dynamics of rRNA transcription, given the well-defined regulatory programs that govern its hierarchical organization^11–14^. Hematopoietic stem cells (HSCs) reside at the top of this hierarchy, maintaining quiescence while balancing self-renewal and differentiation. HSCs give rise to multipotent progenitors (MPPs), which further differentiate into lineage-committed progenitors including common myeloid progenitors (CMPs), granulocyte-monocyte progenitors (GMPs), and megakaryocyte-erythrocyte progenitors (MEPs). These highly proliferative intermediates undergo terminal maturation to sustain the diverse composition of the blood system. Within this hierarchy, cell types differ in characteristics such as cell size, proliferation dynamics, and protein synthesis rates^15,16^. Notably, many of these features are closely linked to ribosome activity, raising the possibility that hematopoietic populations also differentially regulate rRNA transcription and ribosome biogenesis. In further support of a connection between cell identity and rRNA transcription, we previously found that rDNA repeats are bound by multiple hematopoietic lineage-specific transcription factors (TFs), including the myeloid TF CEBPA, which promotes rRNA transcription by facilitating occupancy of Pol I at rDNA repeat loci^17^.

It is well established that rRNA transcription is downregulated during terminal erythroid differentiation^18,19^, and there is additional evidence beyond erythropoiesis that rRNA levels are higher in progenitors compared to mature populations^19,20^. Despite these clues, rRNA levels have not been quantified across carefully dissected lineage trajectories. There is also a lack of insight into the relationship between nascent rRNA transcription and mature rRNA (ribosome subunit) abundance across populations, as well as variations of these levels during the cell cycle. Further, there is limited understanding of how rRNA dynamics are differentially regulated in non-homeostatic contexts, including in malignancy. Ribosome biogenesis is presumed to be upregulated in cancers such as acute myeloid leukemia (AML), consistent with the enlarged nucleoli of leukemic blast cells^21,22^. However, despite the potential prognostic and therapeutic implications of enhanced ribosome biogenesis in AML and other malignancies, rRNA levels in cancers have seldom been directly and precisely compared to matched rapidly-proliferating normal counterpart cells. In addition, it remains unclear whether impairing rRNA transcription levels has consequences beyond simply reducing proliferation.

In this study, we utilize an optimized rRNA FISH-Flow assay to quantify nascent and mature rRNA levels in primary mouse hematopoietic cell types. We demonstrate that rRNA abundance is a cell-type-specific property across normal hematopoietic populations, without rigid correspondence with protein synthesis rates, cell cycling, or between nascent and mature rRNAs. Beyond normal hematopoiesis, we show that rRNA levels are elevated in AML progenitors compared to matched normal cell types. Using publicly available datasets, we find that rDNA accessibility correlates with broad patterns of rRNA abundance in normal and leukemic hematopoiesis and is associated with unique genome-wide chromatin accessibility landscapes. Notably, we find that long-term rDNA repeat inactivation through rDNA methylation does not explain variations in nascent rRNA across hematopoiesis. We also uncover a marked imbalance between mature rRNAs (ribosome subunits), most evident in the myeloid lineage. Finally, we generate a Pol I degron cell line to achieve tunable control of Pol I abundance, and show that graded reductions in rRNA transcription induce and accelerate myeloid differentiation. Collectively, we demonstrate that regulation of rRNA transcription and ribosome subunit abundance is dynamic within and between contexts of hematopoiesis.

## RESULTS

### rRNA FISH-Flow accurately quantifies nascent and mature rRNA

Historical assays for quantification of rRNA transcription have been challenging to apply to complex primary tissues because they require large cell numbers (making them unfeasible for rare cells), require bulk RNA extraction (limiting their utility in heterogeneous tissues), and lack technical precision for subtle changes. We previously published an rRNA FISH-Flow protocol combining RNA FISH and flow cytometry to quantify nascent 47S pre-rRNA, mature 18S rRNA (40S ribosomal subunit), and mature 28S rRNA (60S ribosomal subunit) at single-cell resolution in cell lines^23^ (Figure 1A). To validate the ability of rRNA FISH-Flow to accurately capture dynamic rRNA changes, we engineered an RNA Polymerase I (Pol I) FKBP12^F36V^ (FKBPV) degron system in the mouse ER-HoxA9 myeloid progenitor cell line (Figure 1B). Using CRISPR-mediated homology-directed repair, we inserted an mScarlet-P2A-FLAG-FKBPV cassette at the N-terminus of endogenous *Polr1a*, which encodes the core catalytic subunit of Pol I. Treatment with the small molecule PROTAC dTAG^v^-1 (dTAG) recruits the VHL ubiquitin ligase complex to FKBPV-tagged Pol I, inducing its ubiquitination and proteasomal degradation. This system enables tunable control over Pol I abundance, allowing us to achieve graded inhibition of rRNA transcription for the purpose of validating FISH-Flow.

**Figure 1.**
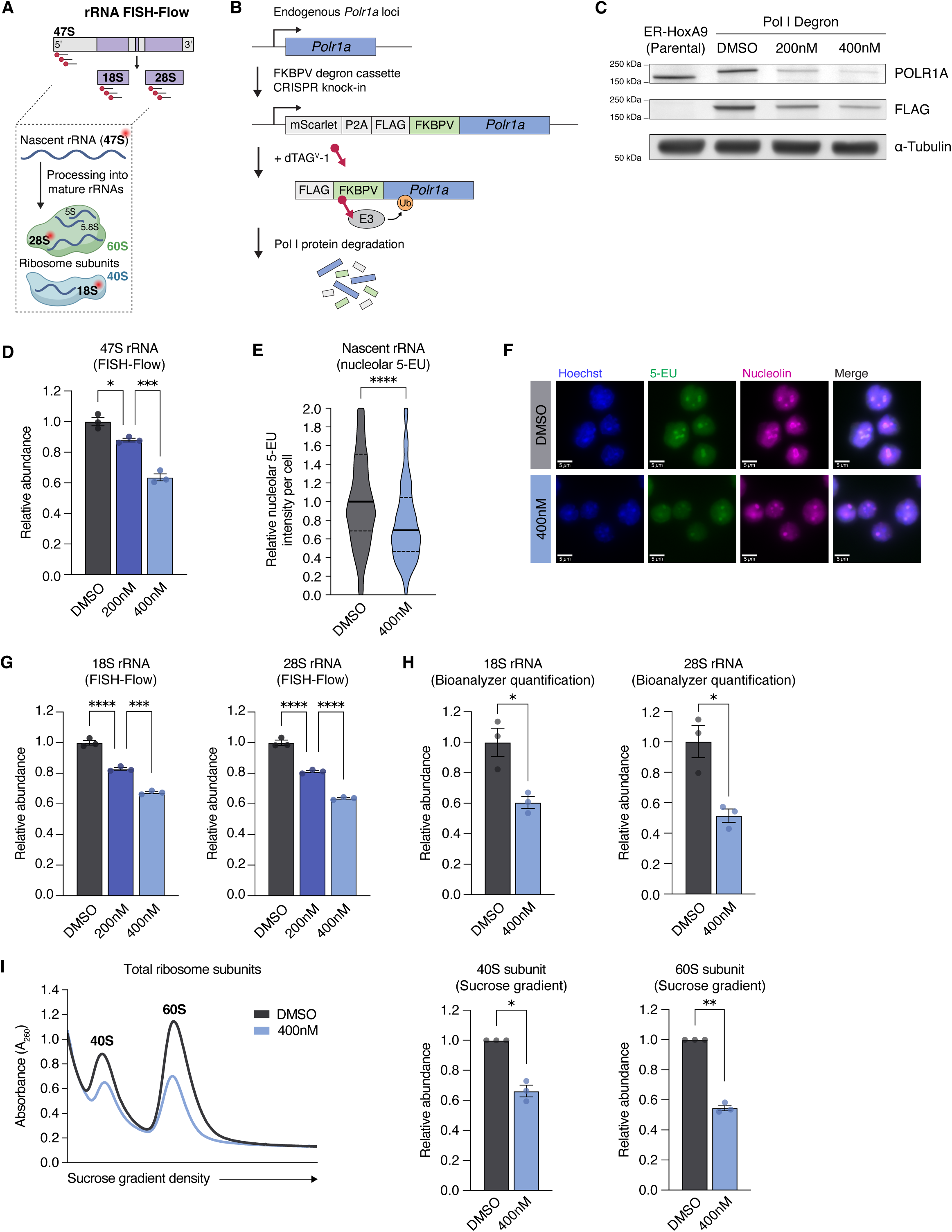
rRNA FISH-Flow accurately quantifies nascent and mature rRNA. **(A)** Schematic of rRNA FISH-Flow. Probes against 47S rRNA are used to quantify nascent rRNA levels, and probes against 18S and 28S (mature) rRNAs are used to quantify abundance of the corresponding ribosome subunits (40S and 60S, respectively). **(B)** Schematic of the design of the Pol I degron cell line. CRISPR-HDR was used to integrate an FKBPV degron cassette into biallelic endogenous *Polr1a* loci in the ER-HoxA9 cell line. The edited loci produce the FLAG-FKBPV-POLR1A fusion protein, which is ubiquitinated and degraded upon treatment with dTAG^v^-1 (dTAG). **(C)** Representative Western blots of parental ER-HoxA9 cells or Pol I degron cells treated with DMSO or dTAG for 24 hours, probed with POLR1A (RPA194) and FLAG antibodies. The parental line shows untagged Pol I (POLR1A blot) at its endogenous molecular weight, and the Pol I degron line shows tagged Pol I at higher molecular weight. The FLAG protein (FLAG blot) is only expressed in the tagged degron cells. The Pol I degron line shows dose-dependent Pol I degradation upon treatment with dTAG. α-Tubulin is shown as a loading control. **(D)** Relative median abundance of nascent 47S rRNA measured by FISH-Flow in Pol I degron cells treated with DMSO or dTAG for 24 hours, normalized to the average of DMSO replicates. n = 3 replicates per condition. Comparisons by one-way ANOVA with Sidak’s multiple comparison testing. **(E)** Relative per-cell nascent rRNA transcription measured by nucleolar 5-EU incorporation in Pol I degron cells treated with DMSO or dTAG for 24 hours, normalized to the median nucleolar 5-EU intensity of DMSO replicates. Violin plots show pooled quantifications of two replicates per condition. Horizontal solid lines depict the median and dotted lines represent quartiles. Comparisons by unpaired, two-sided *t*-test. **(F)** Representative widefield microscopy images of nucleolar 5-EU incorporation (47S rRNA) in Pol I degron cells treated with DMSO (top) or dTAG (bottom) for 24 hours, related to (E). Nuclei are labeled with Hoechst (blue), nascent RNA is labeled with 5-EU (green), and nucleoli are labeled with nucleolin (magenta). Scale bar (white line) at bottom left indicates 5 μm. **(G)** Relative median abundance of mature 18S (left) and 28S (right) rRNAs measured by FISH-Flow in Pol I degron cells treated with DMSO or dTAG for 24 hours, normalized to the average of DMSO replicates for each rRNA. n = 3 replicates per condition. Comparisons by one-way ANOVA with Sidak’s multiple comparison testing. **(H)** Relative abundance of mature 18S (left) and 28S (right) rRNAs quantified using a Bioanalyzer nucleic acid electrophoresis system in Pol I degron cells treated with DMSO or dTAG for 24 hours, normalized to the average of DMSO replicates for each rRNA. Each point depicts the average relative abundance from two technical replicates per sample. n = 3 replicates per condition. Comparisons by unpaired, two-sided *t*-test. **(I)** Sucrose gradient fractionation quantification of total 40S and 60S ribosomal subunits in Pol I degron cells treated with DMSO or dTAG for 24 hours. Left panel: representative A_260_ absorbance tracings. Middle panel: relative abundance of total 40S subunits. Right panel: relative abundance of total 60S subunits. Quantifications are normalized to paired DMSO controls for each subunit. n = 3 replicates per condition. Comparisons by one-sample *t*-test. All bar graphs show mean ± standard error of mean (SEM). ns (*p* ≥ 0.05), * (*p* < 0.05), ** (*p* < 0.01), *** (*p* < 0.001), **** (*p* < 0.0001)

Pol I degron cells treated with 200 nM or 400 nM dTAG for 24 hours showed dose-dependent reduction in Pol I abundance (Figure 1C, Supplemental Figure S1A). Using rRNA FISH-Flow, we found that these reductions in Pol I abundance led to decreases in nascent 47S rRNA of 12% (200 nM) and 36% (400 nM) (Figure 1D). Consistent with this, median nucleolar incorporation of 5-ethynyl uridine (5-EU), an orthogonal measure of nascent rRNA synthesis, was reduced by 31% in Pol I degron cells treated with 400 nM dTAG for 24 hours (Figure 1E-F). Northern blot analysis confirmed that the 47S rRNA FISH-Flow probe set, which targets the distal 5’ external transcribed spacer that is rapidly cleaved during early pre-rRNA processing, predominantly detects full-length 47S transcripts, indicating that the FISH-Flow signal primarily reflects nascent pre-rRNA and not other cleavage intermediates (Supplemental Figure S1B).

We next tested the ability of FISH-Flow to quantify downstream changes in the abundance of mature 18S and 28S rRNAs (40S and 60S ribosomal subunits). Using rRNA FISH-Flow, we found that Pol I degron cells treated with dTAG for 24 hours showed ∼18% (200 nM) or ∼36% (400 nM) reduction in both 18S and 28S rRNAs (Figure 1G). The magnitudes of these changes were validated by quantifying mature rRNA abundance using a Bioanalyzer automated nucleic acid electrophoresis system (Figure 1H) and by quantifying total cellular 40S and 60S ribosomal subunit abundance using sucrose gradient fractionation (Figure 1I).

The inclusion of a DAPI staining step in FISH-Flow also enabled us to stratify cell cycle populations for separate quantification. At baseline as well as at all dTAG doses, cells in S/G2/M showed higher rRNA levels than those in G1, as expected (Supplemental Figure S1C-E). Graded reductions in nascent and mature rRNA levels were proportionately observed across cell cycle phases (Supplemental Figure S1C-E).

Together, these findings establish that rRNA FISH-Flow enables precise quantification of small dynamic changes in the abundance of nascent and mature rRNAs, with the ability to uncouple quantification across cell cycle phases.

### Optimization of rRNA FISH-Flow for quantification of nascent and mature rRNA levels in primary mouse bone marrow

We next adapted rRNA FISH-Flow for primary mouse bone marrow (BM) to profile levels of nascent rRNA and mature ribosome subunits across hematopoietic cell types (Figure 2A). In this optimized protocol, BM samples were harvested and stained with surface marker antibodies, subjected to gentle fixation and permeabilization, and then hybridized with pools of probes against mouse 47S, 18S, or 28S rRNA. The duration of time between mouse euthanasia and sample fixation was kept under an hour to minimize *ex vivo* changes to ribosome biogenesis and accurately capture *in vivo* rRNA levels. After staining with DAPI, we used flow cytometry to acquire fluorescence intensity of surface markers, DNA content, and rRNA abundance. This enabled quantification of rRNA levels in populations stratified by cell identity and cell cycle phase. rRNA levels were quantified by extracting the median fluorescence intensity (MFI) of rRNA probes in each cell type of interest. To compare rRNA levels in a standardized manner between cell types and across biological replicates, all quantifications were internally normalized to the median signal of the total BM CD45^+^ population for each mouse (i.e., median of CD45^+^ cells = 1).

**Figure 2.**
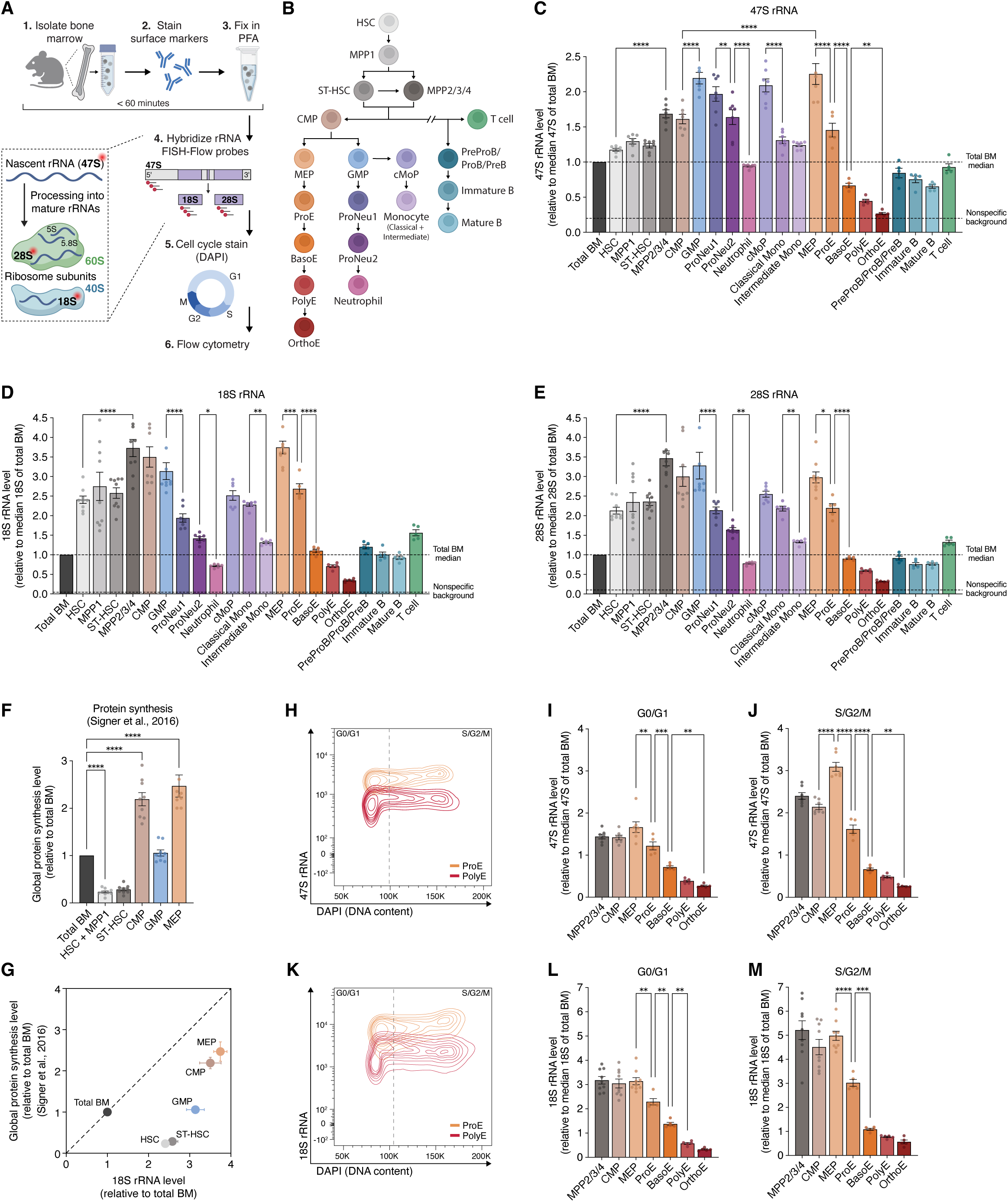
Nascent and mature rRNA levels are tightly associated with hematopoietic cell identity. **(A)** Schematic of the rRNA FISH-Flow assay used to quantify rRNA levels in primary mouse bone marrow (BM). **(B)** Schematic of the hematopoietic hierarchy, depicting cell types analyzed in this study. HSC, hematopoietic stem cell; MPP1, multipotent progenitor 1; ST-HSC, short-term HSC; MPP2/3/4, multipotent progenitors 2, 3, and 4; CMP, common myeloid progenitor; GMP, granulocyte-monocyte progenitor; ProNeu1/ProNeu2, pro-neutrophil 1 and 2; cMoP, common monocyte progenitor; Mono, monocyte; MEP, megakaryocyte-erythroid progenitor; ProE, proerythroblast; BasoE, basophilic erythroblast; PolyE, polychromatic erythroblast; OrthoE, orthochromatic erythroblast; PreProB/ProB/PreB, pre-pro B cell, pro B cell, and pre B cell; Immature/Mature B, immature/mature B cell. **(C-E)** Relative abundance of 47S rRNA (C), 18S rRNA (D), and 28S rRNA (E) in each cell type, normalized to the median level of each respective rRNA in the total BM CD45^+^ population (upper dashed line). The nonspecific background signal of each rRNA probe set is also shown (lower dashed line). **(F)** Relative mouse *in vivo* global protein synthesis level in different hematopoietic cell types, normalized to the mean of total BM. Data obtained from Signer et al., 2016^15^. **(G)** Comparison of relative 18S rRNA abundance (as shown in D) and relative global protein synthesis level (as shown in F) across hematopoietic cell types. **(H)** Representative flow cytometry plot comparing 47S rRNA abundance across cell cycle phases (DAPI) in ProE (top, orange) and PolyE (bottom, red) populations. **(I-J)** Relative abundance of 47S rRNA (normalized to the median 47S level of total BM) across the erythroid lineage in cells in G0/G1 (I) and S/G2/M (J) phases of the cell cycle. **(K-M)** Same as H-J, showing 18S rRNA. In some cases, replicates outside of the axis limits are not shown ( ≤ 2 replicates per cell type). All bar graphs and scatter plots show mean ± SEM. ns (*p* ≥ 0.05), * (*p* < 0.05), ** (*p* < 0.01), *** (*p* < 0.001), **** (*p* < 0.0001) by one-way ANOVA with Sidak’s multiple comparison testing (n = 5-9 biological replicates per cell type).

During protocol optimization, we observed that while bone marrow cell types collectively fell within a broad distribution of rRNA signal, eosinophils (marked by Siglec F surface expression) showed aberrantly high probe intensity (Supplemental Figure S2A). Given reports that eosinophil granules are susceptible to nonspecific binding by both probes and fluorophores^24^, we excluded eosinophils (accounting for <5% of total BM cells) from all analyses. We also used anti-GFP-mRNA probes to measure nonspecific background fluorescence intensity relative to each rRNA probe set, and used it to set the lower limit of reliable quantification for each rRNA (Supplemental Figure S2B-E). Apart from eosinophils, all other cell types showed low nonspecific signal. In summary, we optimized rRNA FISH-Flow to render it suitable for precise quantification of rRNA levels across a wide dynamic range in heterogeneous primary cell populations.

### Nascent and mature rRNA levels are tightly associated with hematopoietic cell identity

Using rRNA FISH-Flow, we identified 9-fold variation in nascent 47S rRNA abundance and 11-fold variation in mature 18S and 28S rRNA levels across normal mouse hematopoietic cell types (Figure 2B-E; Supplemental Figure S3; Supplemental Table S1). rRNA levels were intermediate in stem cells, highest in progenitors, and progressively lower in maturing populations across different lineages. Specifically, HSCs exhibited nascent rRNA levels 1.2x and mature rRNA levels 2.4x the median of total BM. Nascent and mature rRNA levels increased from HSCs to multipotent progenitors (MPPs) (MPP2/3/4 47S levels 1.7x and 18S levels 3.7x total BM median), and the highest rRNA levels were found in committed progenitor populations including MEPs (47S levels 2.3x and 18S levels 3.7x total BM median) and GMPs (47S levels 2.2x and 18S levels 3.1x total BM median).

Different lineages displayed varied dynamics of rRNA decrease or plateau during terminal maturation. During erythroid differentiation, rRNA levels progressively decreased and were lowest in orthochromatic erythroblasts, the stage of erythropoiesis prior to enucleation and production of reticulocytes (47S and 18S levels both 0.3x total BM median). This recapitulated the previously reported shutdown of rRNA transcription in terminal erythropoiesis^18,19^. Distinct trends were observed during myeloid differentiation, where nascent and mature rRNA levels progressively decreased from GMPs to mature neutrophils and monocytes.

We compared mature rRNA abundance with previously-published data of *in vivo* mouse protein synthesis rates in hematopoietic stem and progenitor populations (HSPCs)^15^ (Figure 2F-G). Global translation levels were very low in HSCs - only 0.2x the mean across total BM. This contrasted with the relatively high level of mature rRNAs in HSCs, and surprisingly indicated that HSCs actively transcribe rRNA and maintain robust ribosome subunit content despite low translation rates. Further committed progenitor populations such as CMPs, GMPs, and MEPs had higher global translation rates, though notably these rates also did not perfectly correlate with mature rRNA levels. Thus, protein synthesis rates cannot be used to reliably infer ribosome subunit abundance or rRNA synthesis rates.

Different hematopoietic populations have different cycling parameters, leading us to consider whether differences in cell cycle phase composition could act as a potential confounder in our quantification. To account for this, we used DAPI profiling to compare rRNA levels between G0/G1 and S/G2/M populations of each primary cell type. Within many cell types, cells in S/G2/M consistently exhibited higher levels of each rRNA species (Supplemental Figure S4A-C). However, dynamic rRNA trends were preserved between cell types and within differentiation trajectories (Figure 2H-M; Supplemental Figure S4A-C and S5A-F), and we observed multiple cases where cells with near-identical cell cycle distributions showed clear differences in nascent and mature rRNA levels (e.g., pro-erythroblasts vs. polychromatic erythroblasts). Overall, we did not observe any strong correlation between cell cycle distribution and nascent or mature rRNA levels (Figure 3A-B; Supplemental Figure S5G-H), indicating that trends in rRNA levels are not artifacts of cell cycle distribution differences and instead reflect true biological variation determined by cell identity. Collectively, our results indicate that nascent and mature rRNA levels are a cell-type-specific property within the normal homeostatic hematopoietic system.

**Figure 3.**
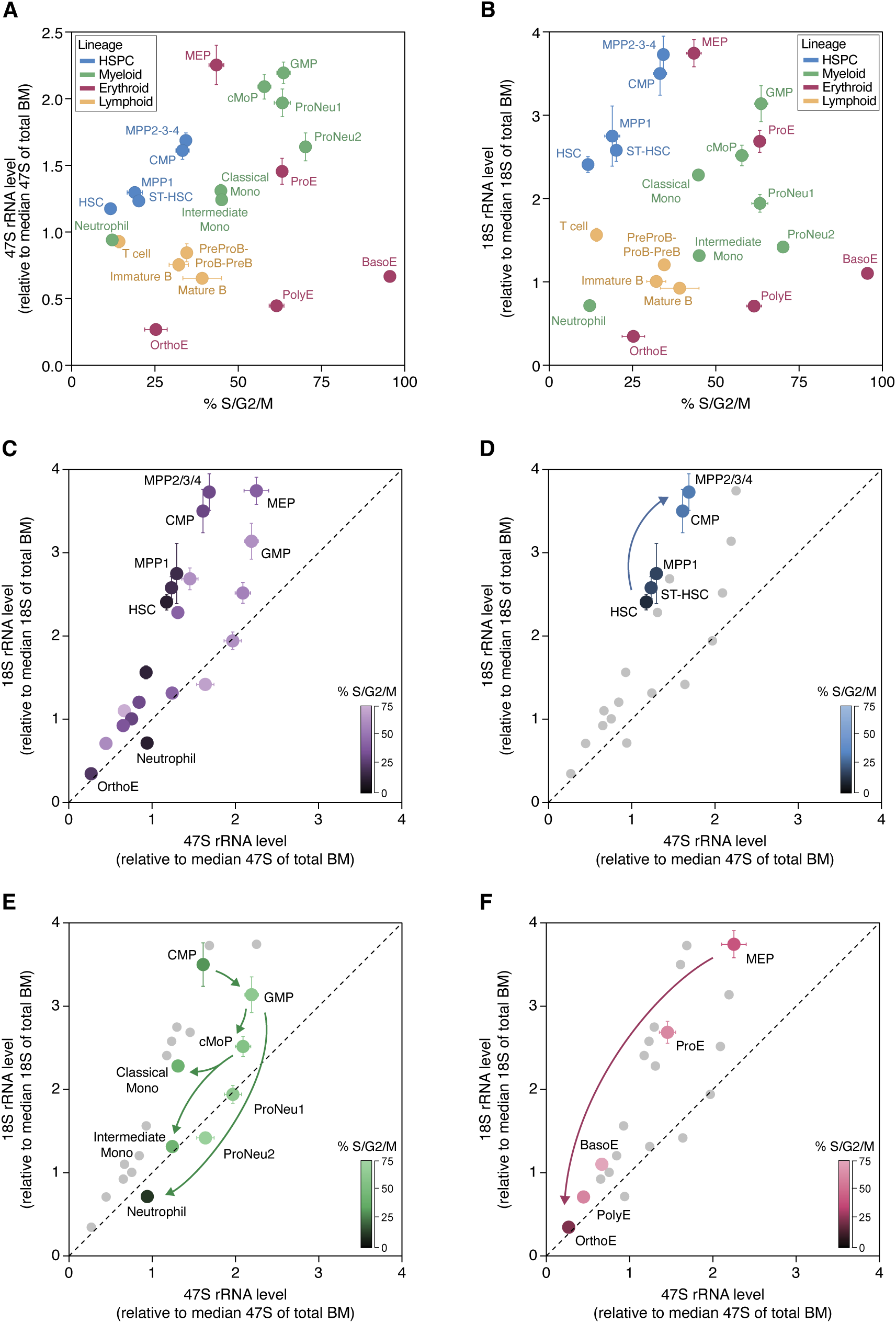
Correspondence between nascent and mature rRNA levels is non-linear and independent of cell cycle distribution. **(A-B)** Comparison of cell cycle phase distribution (% S/G2/M) and 47S rRNA level (A) or 18S rRNA level (B) across normal hematopoietic lineages. **(C-F)** Comparison of relative nascent rRNA (47S rRNA) and ribosome subunit abundance (18S rRNA) across all normal hematopoietic cell types (C), HSPCs (D), myeloid lineage (E), and erythroid lineage (F). The color of each cell type represents the percentage of cells in S/G2/M cell cycle phases. Color scales are capped at 75%. All points represent mean ± SEM.

### Correspondence between nascent and mature rRNA levels is non-linear

We next investigated the relationship between nascent and mature rRNA levels across the hematopoietic system. By comparing 47S and 18S rRNA levels across cell types, we found that the overall distribution was skewed towards the 18S axis, without a definite linear relationship (Figure 3C-F; Supplemental Figure S5I-L). Our data instead indicated a greater spread in 18S rRNA compared to 47S, and a lack of strict correspondence, with mismatches along both dimensions. Intermediate monocytes showed proportional nascent and mature rRNAs relative to total BM (47S 1.2x and 18S 1.3x total BM median) (Figure 3E). Classical monocytes, on the other hand, had nearly the same level of nascent rRNA (47S 1.3x total BM median), but almost double the mature rRNA of intermediate monocytes (18S 2.3x total BM median) (Figure 3E). This was despite both populations having 45% of cells in S/G2/M (Supplemental Figure S5G). Conversely, despite equivalent mature rRNA levels, nascent rRNA in MEPs was 1.3-fold that of MPP2/3/4 (Figure 3C). Cells with greater S/G2/M percentages did not consistently fall in any particular zone of the plot, indicating that cell cycle distribution cannot completely explain correspondence (or lack thereof) between nascent and mature rRNA levels. Rather, our findings suggest additional mechanisms beyond rRNA synthesis that regulate mature rRNA abundance across cell types, such as ribosome subunit lifespan. In summary, rRNA is likely tightly regulated at the levels of both transcriptional output as well as mature rRNA (ribosome subunit) stability in different homeostatic cell types.

### Nascent and mature rRNA levels are higher in AML compared to normal hematopoietic counterparts

To explore trends in rRNA levels beyond normal hematopoiesis, we quantified nascent and mature rRNA abundance in mice with acute myeloid leukemia (AML). While it is widely presumed that ribosome biogenesis is upregulated in a variety of malignancies, detailed profiling of nascent rRNA transcription and ribosome subunit abundance in matched normal and cancer cell type populations is largely lacking. We used two transgenic AML mouse models driven by combinations of two of the three most commonly mutated genes in human AML (DNMT3A, FLT3, NPM1)^25^. The first model combines hematopoietic *Dnmt3a* knockout and homozygous germline *Flt3* internal tandem duplication (*Flt3*^ITD^)^26^ (model referred to as ‘DF’), and the second model combines inducible heterozygous *Npm1*^c^ mutation and *Flt3*^ITD^ ^27^ (model referred to as ‘NF’). To compare rRNA levels in AML cells to normal hematopoietic populations, we spiked in wild type BM cells into AML samples prior to fixation, and normalized all AML rRNA MFIs to the median of the wild type spike-in (Figure 4A; Supplemental Figure S6, S7A). As previously reported^26,27^, AML BM showed expansion of the lineage^-^, c-Kit^+^ populations, and we applied identical gating strategies as wild type BM to define matched progenitor cell subpopulations in AML (Supplemental Figure S6; Supplemental Table S1).

**Figure 4.**
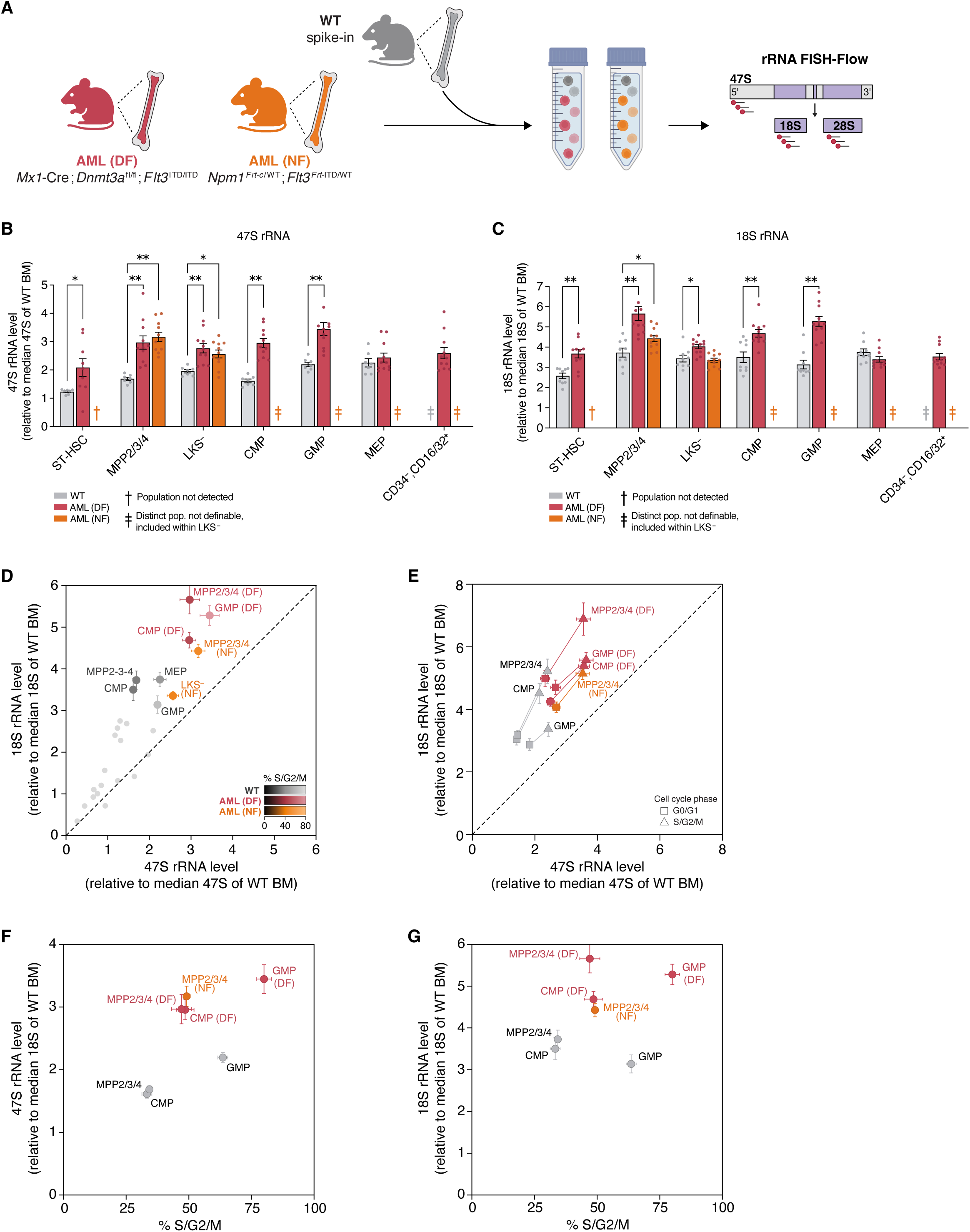
Nascent and mature rRNA levels are higher in AML compared to normal hematopoietic counterparts. **(A)** Schematic of wild-type (WT) BM spike-in used to normalize AML (DF) and AML (NF) rRNA MFIs to total WT BM. For all panels, WT is shown in grey, AML (DF) is shown in red, and AML (NF) is shown in orange. **(B-C)** Relative levels of 47S rRNA (B) and 18S rRNA (C) in AML stem and progenitor-like cells compared to matched normal counterparts. All levels normalized to the median rRNA levels of total WT BM. **†** indicates populations that were not detected at measurable frequencies. **‡** indicates populations that were not separately definable (did not form distinct populations on flow cytometry) and are instead included within the LKS^-^ population. **(D)** Comparison of relative nascent rRNA (47S rRNA) and ribosome subunit abundance (18S rRNA) across AML populations and normal counterparts (normalized to the median rRNA levels of total WT BM). The color of each cell type represents the percentage of cells in S/G2/M cell cycle phases. **(E)** Comparison of 47S and 18S rRNA levels in cell cycle phase subpopulations of AML and WT cell types. For each cell type, the G0/G1 (square) and S/G2/M (triangle) populations are shown independently and connected by a line. **(F-G)** Comparison of cell cycle phase distribution (% S/G2/M) and relative 47S rRNA level (F) or 18S rRNA level (G) across AML and WT cell types. In some cases, replicates outside of the axis limits are not shown ( ≤ 2 replicates per cell type). All bar graphs and scatter plots show mean ± SEM. ns (*p* ≥ 0.05), * (*p* < 0.05), ** (*p* < 0.01), *** (*p* < 0.001), **** (*p* < 0.0001) by two-sided Mann-Whitney tests with FDR correction (Benjamini–Hochberg) (n = 7-11 biological replicates per cell type).

Across both AML models, we found that nascent and mature rRNA levels were higher in specific AML HSPCs compared to their normal counterparts (Figure 4B-D; Supplemental Figure S7B-C). AML (DF) GMP-like cells had the highest nascent rRNA levels (3.4x the median of total wild type BM and 1.6x normal GMPs), exceeding the levels of all normal hematopoietic populations. Nascent rRNA abundance was similarly elevated 3x-3.2x total normal BM and 1.8x-1.9x normal counterparts in AML (DF) CMP-like and MPP2/3/4-like cells and AML (NF) MPP2/3/4-like cells. Mature rRNA levels were highest in AML (DF) MPP2/3/4-like cells (18S rRNA 5.7x median of total normal BM and 1.5x normal MPP2/3/4) and were also elevated in AML (DF) CMP-like and GMP-like cells and AML (NF) MPP2/3/4-like cells compared to their normal counterparts and all normal populations. While many of these AML populations had a greater percentage of cells in S/G2/M compared to their normal counterparts, elevated rRNA levels in AML persisted in cell cycle phase-stratified populations (Figure 4E; Supplemental Figure S7D-H). Strikingly, less-proliferative populations within both AMLs (AML CMP-like and MPP2/3/4-like cells) showed higher rRNA levels compared to more-proliferative normal populations (GMPs) (Figure 4F-G). Through these analyses, we identified that different AML models exhibit distinct patterns of rRNA abundance, with elevated nascent and mature rRNA in a subset of AML stem and progenitor-like populations compared to normal counterpart cells. In the AML (DF) model, both nascent and mature rRNA levels are elevated across most HSPC populations, whereas in the AML (NF) model, marked increases are largely restricted to 47S rRNA in MPP2/3/4-like cells. Notably, HSPC-like populations in both AML models have been reported to exhibit robust transplantability^26,28^, indicating a potential link between elevated rRNA abundance and leukemia stemness.

### rDNA transcriptional machinery components are downregulated during myeloid maturation

We next examined potential mechanisms that might explain trends in hematopoietic rRNA transcription. We first assessed whether gene expression of components of the core rDNA transcriptional machinery correlated with trends in nascent rRNA abundance. This machinery consists of factors and complexes that coordinate rRNA transcription, including UBTF, which occupies and activates rDNA repeats, its interaction partner TCOF1, the TATA-binding protein complex SL-1 (5 subunits), RNA Pol I (13 subunits), the RNA Pol I initiation factor RRN3, and the transcription termination factor TTF1. In addition, MYC is known to promote transcription of the genes encoding several of these components^29,30^. We quantified mRNA levels of each of these factors across the myeloid lineage by analyzing two published mouse single-cell RNA-sequencing (scRNA-seq) datasets^31,32^. In both datasets, expression of the core rDNA machinery was downregulated during neutrophil differentiation (Figure 5A-B), consistent with the progressive decrease in nascent rRNA. Interestingly, core machinery expression remained high across HSCs and progenitor populations despite increasing nascent rRNA levels. Some components of this machinery are known to also be regulated by post-translational modifications; these gene expression measurements provide only a broad readout of machinery abundance. Therefore, while gene expression of the core rDNA machinery correlates with and may regulate patterns of nascent rRNA in certain populations such as maturing neutrophils, additional mechanisms likely control the activity of rDNA machinery during HSPC maturation.

**Figure 5.**
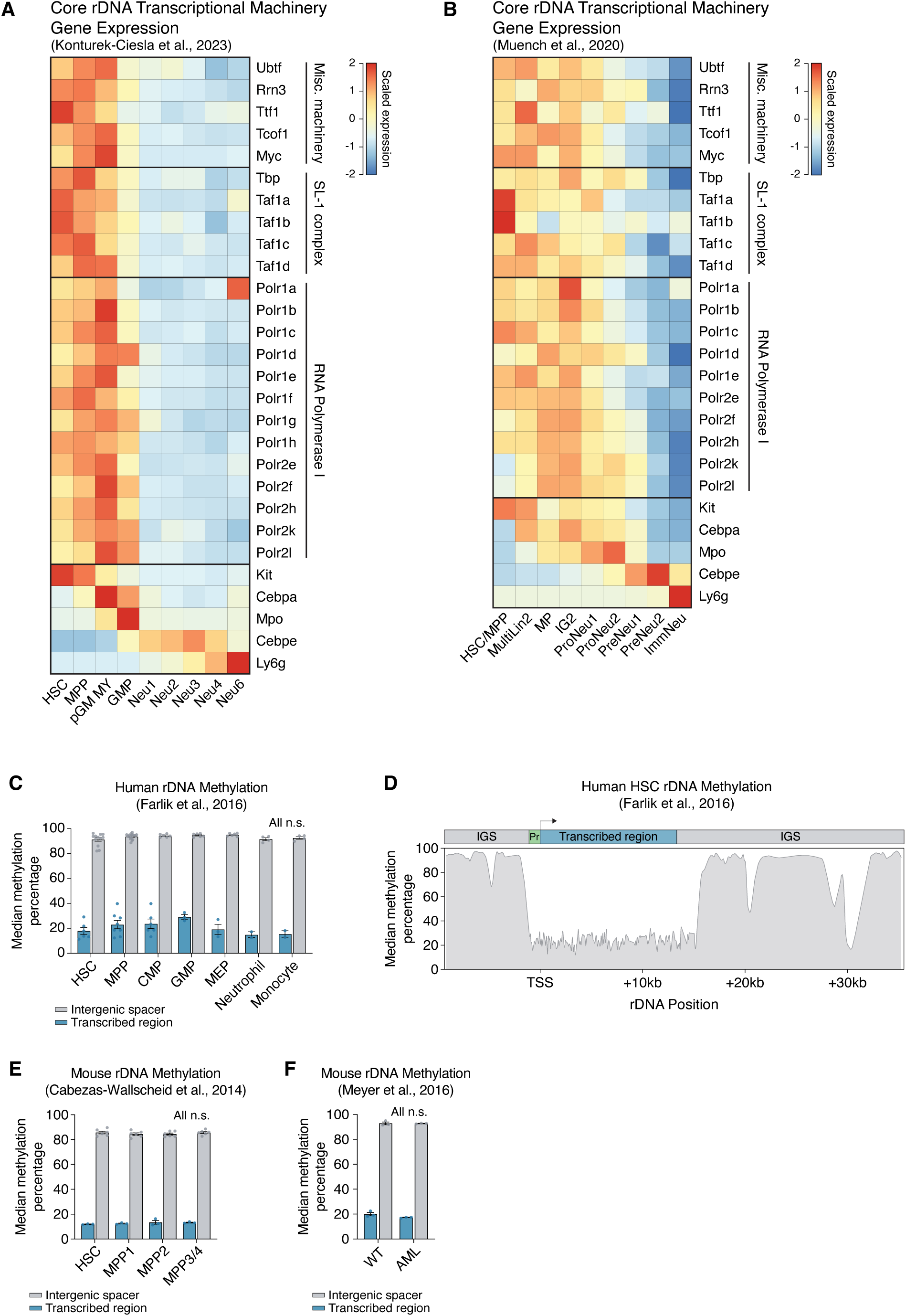
Distinct regulation of rDNA transcriptional machinery and rDNA methylation across hematopoiesis. **(A-B)** Heatmaps showing gene expression patterns of core rDNA transcriptional machinery components across mouse stem and progenitor cells and the myeloid lineage, quantified using scRNA-seq. Genes are grouped by protein complex, and positive control hematopoietic markers are shown at the bottom (*Kit* marks HSPCs, *Cebpa* and *Mpo* mark myeloid progenitors, *Cebpe* and *Ly6g* mark neutrophils). Gene expression values are scaled to a fixed range from –2 to 2 using z-score normalization across all cell types. Cell identity labels (columns) are listed as assigned in the original scRNA-seq publications, and broadly recapitulate the HSPC-to-neutrophil trajectory assayed by our study. pGM MY, pro-granulocyte-monocyte myeloid progenitor; MultiLin2, multi-lineage progenitor 2; MP, monocyte-committed progenitor; IG2, bipotential monocytic-granulocytic intermediates; ImmNeu, immature neutrophil. **(C)** Median CpG methylation percentages across the rDNA transcribed region and intergenic spacer (IGS) in normal human hematopoietic cell types. **(D)** Representative track depicting CpG methylation percentages across the rDNA transcribed region and IGS in normal human HSCs. Signal at each position represents composite methylation percentage across all rDNA repeats from all cells in the assayed population. The top schematic indicates the location of the rDNA promoter, transcribed region, and IGS. **(E-F)** Same as in C, depicting normal mouse cell types (E) and mouse WT LSK cells and AML (*Mx-Cre/+;Flt3^ITD/ITD^;Dnmt3a^fl/-^*) cells (F). All bar graphs show mean ± SEM. All comparisons are ns (*p* ≥ 0.05) by two-sided Mann-Whitney tests with FDR correction (Benjamini–Hochberg).

### rDNA repeat accessibility but not methylation correlates with patterns of nascent rRNA abundance in normal and leukemic hematopoiesis

To better resolve mechanisms potentially influencing trends in nascent rRNA abundance, we investigated whether rDNA chromatin state, which distinguishes active and inactive rDNA genes, might underlie trends in rRNA transcription. Core features of rDNA chromatin regulation, including CpG methylation and repeat accessibility, are broadly conserved across mammalian systems. We first profiled CpG methylation along the rDNA locus, which lacks CpG islands and instead contains CpGs distributed throughout the transcribed region and intergenic spacer (IGS)^7,33,34^. The IGS is densely methylated in all rDNA repeats (both active and inactive), while the transcribed region is only methylated in long-term inactive copies^7,33,34^. To quantify methylation levels of these rDNA regions, we mapped published human^35^ and mouse^36^ whole genome bisulfite sequencing (WGBS) datasets to custom genome assemblies previously optimized by us for high-throughput rDNA mapping and visualization^37^. As expected, CpGs in the IGS were highly methylated in all cell types (∼85% in mouse, ∼90-95% in human) (Figure 5C-E). Methylation levels were lower across the rDNA transcribed region for all cell types (∼13% in mouse, ∼15-30% in human) and showed no meaningful variation between cell types. We also profiled a published reduced representation bisulfite sequencing (RRBS) dataset^26^ from the AML (DF) mouse model that we used for rRNA FISH-Flow, and found no difference in rDNA methylation levels between normal and leukemic cells (Figure 5F). We therefore found that the proportion of long-term inactivated (methylated) rDNA copies does not change significantly in normal or malignant hematopoiesis, and cannot account for differences in nascent rRNA.

To further explore rDNA chromatin state mechanisms, we next investigated rDNA accessibility. In any given cell, a subset of rDNA repeats are accessible (depleted of histones across the entire 13-kb transcribed locus) and available for Pol I to transcribe, while the rest remain chromatinized^5,6^. We quantified rDNA accessibility across normal hematopoietic cell types by mapping published human^38^ and mouse^39^ scATAC-seq datasets to rDNA, using standardized reference maps to assign cell identities (Figure 6A-D). Accessibility was highest in GMPs and progressively lower in maturing monocytes and neutrophils; early erythroid progenitors also demonstrated increased accessibility relative to terminal erythroblasts. Importantly, accessibility was lower in HSCs and early progenitors compared to GMPs and erythroid progenitors, providing a potential explanation for varying nascent rRNA levels across HSPCs (Figure 6A,D; Supplemental Figure S8A-B).

**Figure 6.**
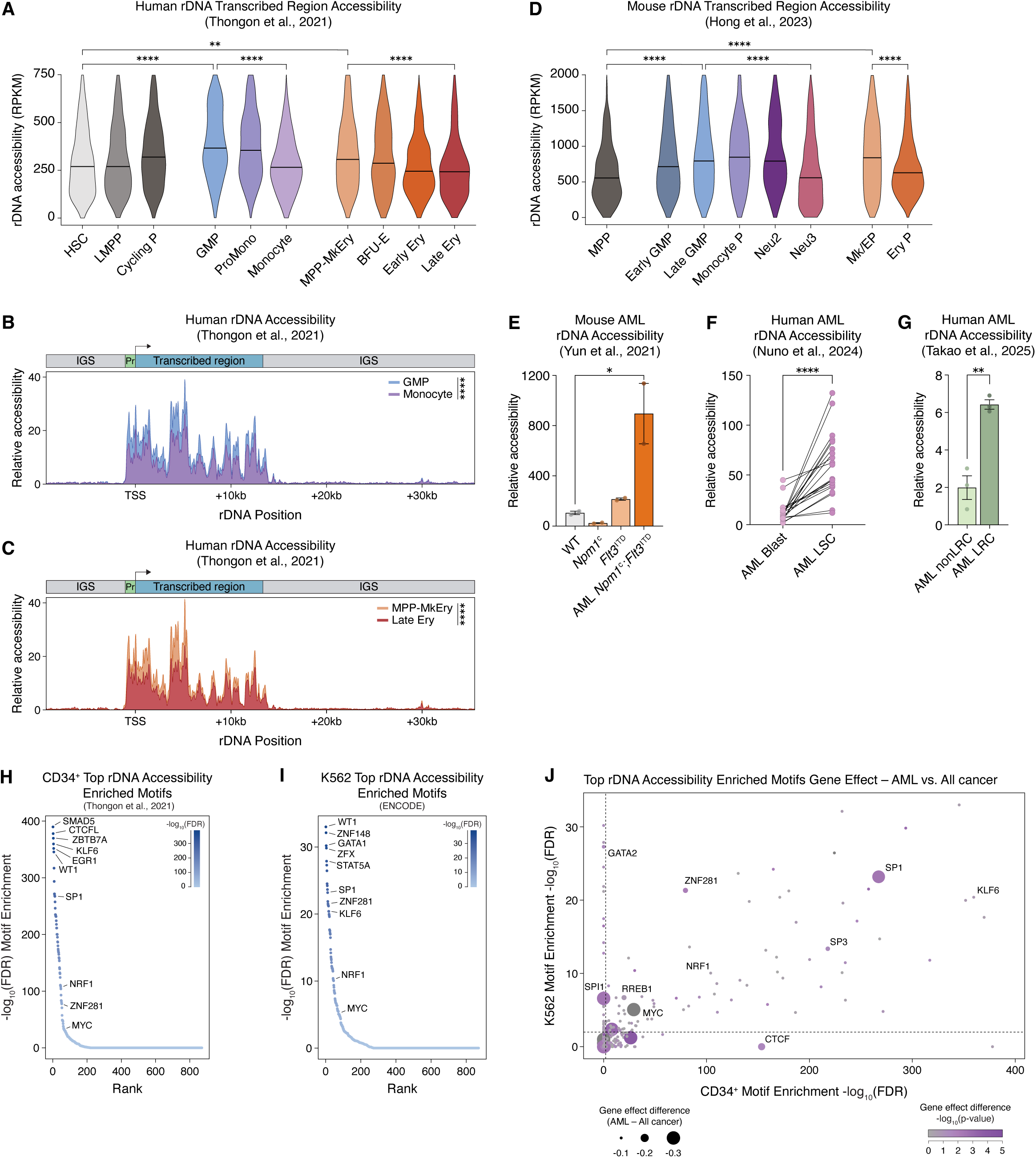
Accessibility of rDNA repeats correlates with patterns of rRNA transcription and with distinct genome-wide regulatory landscapes. **(A)** Violin plots depicting accessibility of the rDNA transcribed region across normal human cell types quantified using scATAC-seq. Values represent normalized accessibility expressed as RPKM. Horizontal lines within each distribution indicate the median accessibility of each cell type. Comparisons by two-sided Mann-Whitney tests with FDR correction (Benjamini–Hochberg). LMPP, lymphoid-myeloid-primed progenitor; Cycling P, cycling progenitor; MPP-MkEry, multipotent megakaryocyte-erythroid progenitor; BFU-E, burst-forming unit-erythroid; Early/Late Ery, early/late erythroblast. **(B-C)** Representative tracks depicting accessibility along the rDNA transcribed region and intergenic spacer (IGS) in human myeloid (B) and erythroid (C) lineages. The top schematic indicates the location of the rDNA promoter, transcribed region, and IGS. Signal at each position represents composite accessibility across all rDNA repeats from all cells in the assayed population. Comparisons by two-sided Mann-Whitney tests with FDR correction (Benjamini–Hochberg). **(D)** Same as in A, showing mouse cell types. Monocyte P, monocyte progenitor; Neu2/Neu3, neutrophil 2 and 3; Mk/EP, megakaryocyte-erythroid progenitor; EryP, erythroid progenitor. **(E)** Relative accessibility of the rDNA transcribed region in WT HSPCs, pre-AML *Npm1^c^* HSPCs, pre-AML *Flt3*^ITD^ HSPCs, and AML *Npm1^c^*;*Flt3*^ITD^ HSPCs. Bars show mean ± SEM with comparisons by one-way ANOVA with Sidak’s multiple comparison testing. **(F)** Relative accessibility of the rDNA transcribed region in paired AML blast cells and leukemia stem cells (LSCs) from the same patients. Pairs are connected by solid lines. Comparisons by two-sided paired *t*-test. **(G)** Relative accessibility of the rDNA transcribed region in proliferative AML non-label-retaining cells (non LRCs) and quiescent AML label-retaining cells (LRCs). AML cells are from a human patient-derived xenograft driven by MOZ-CBP translocation. Bars show mean ± SEM with comparisons by two-sided unpaired *t*-test. **(H-I)** Transcription factor motifs enriched in genome-wide chromatin accessibility peaks showing increased accessibility in primary human CD34⁺ cells (H) or K562 cells (I) with higher rDNA accessibility. Higher rDNA accessibility corresponds to the top 10% (H) and top 20% (I) rDNA accessibility cell populations. -log_10_(FDR) values indicate motif enrichment significance from two-sided Mann-Whitney tests with FDR correction (Benjamini–Hochberg). **(J)** Comparison of CRISPR dependency gene effect scores in AML cell lines versus all cancer cell lines for TFs enriched in H and I. More negative scores indicate greater dependency (gRNA dropout after CRISPR knockout). Point size represents the magnitude of the difference in gene effect scores (AML minus all cancer lines), with larger points indicating stronger selective dependency in AML cell lines. Dot color denotes the statistical significance of the gene effect difference. Axes show -log_10_(FDR) from two-sided Mann-Whitney tests with FDR correction (Benjamini–Hochberg), and dot color represents -log_10_(*p*-value) from unpaired two-sided *t*-tests comparing gene effect scores. Dashed lines indicate a TF motif enrichment significance threshold of *p* < 0.01. For all comparisons, ns (*p* ≥ 0.05), * (*p* < 0.05), ** (*p* < 0.01), *** (*p* < 0.001), **** (*p* < 0.0001).

To investigate whether differential rDNA accessibility could explain elevated nascent rRNA levels in AML, we also mapped published bulk ATAC-seq datasets from human and mouse AML. HSPCs from a mouse AML model driven by co-mutated *Npm1*^c^ and *Flt3*^ITD^ (comparable to the AML NF model used in our FISH-Flow quantifications) displayed higher rDNA accessibility compared to normal HSPCs and pre-AML *Npm1*^c^ or *Flt3*^ITD^ alone^40^ (Figure 6E). Across a cohort of human AML patient samples representing multiple different AML driver mutations, the leukemia stem cell (LSC) population (defined as CD34^+^, CD38^-^, CD99^+^, TIM3^+^) exhibited higher rDNA accessibility compared to the rest of the leukemic blast population, indicating both heterogeneity within the AML hierarchy, as well as the possibility that LSCs may differ from normal HSCs in having an increased number of active rDNA copies^41^ (Figure 6F). In a study that tracked post-mouse-transplant cell division in an AML patient-derived xenograft, rDNA accessibility was higher in the quiescent (non-dividing) label-retaining cell (LRC) population compared to non-LRC (dividing) cells^42^ (Figure 6G); notably, only the former was reported as being able to engraft leukemia in a serial mouse transplant. Overall, we found that rDNA accessibility, marking the number of active rDNA loci, correlates with and may partly explain differences in nascent rRNA abundance within and between normal and leukemic hematopoiesis.

We next used scATAC-seq datasets from human normal hematopoiesis^38^ and the K562 myeloid cell line (ENCODE^43^) to identify transcription factors (TFs) correlating with and potentially driving differential rDNA accessibility and activity. To investigate whether cells with higher numbers of open rDNA repeats are enriched for specific TF programs, we first stratified primary HSPCs based on rDNA accessibility into top 10% and bottom 10% cell populations, and K562 cells into top 20% and bottom 20%. We compared genome-wide chromatin accessibility landscapes between these subpopulations, and performed TF motif enrichment analysis on global differential peaks in cells showing higher rDNA accessibility (Figure 6H-I). Motifs of KLF6, ZNF281, MYC, SP1, and NRF1 were enriched in both HSPCs and K562 cells with high rDNA accessibility. Notably, many TFs that are established regulators of HSPC identity (e.g., RUNX, GATA, and HOX family TFs) were not enriched in these analyses. Overlap of TF motifs between primary HSPCs and K562 cells suggests that activated chromatin accessibility landscapes driven by these shared factors may represent a common feature of cells with higher numbers of active rDNA repeats.

Aiming to identify novel cell-type-specific regulators of rDNA accessibility, we integrated these TF results with cell line dependencies from the Cancer Dependency Map (DepMap) Project^44^, which contains whole genome CRISPR-based gene dependency scores from human cell lines spanning over 25 cell types. Given that rRNA transcription is vital for proliferation and survival, we reasoned that TFs that regulate any step upstream of hematopoietic rDNA accessibility would likely show up in DepMap either as global or myeloid-specific dependencies. We obtained CRISPR dependency gene effect scores of all factors in 20 AML cell lines and across all 1178 cancer cell lines, and found that SP1 has selectively higher dependency in myeloid lines, while ZNF281 and NRF1 are universally essential across all lines (Figure 6J; Supplemental Figure S8C-D). This nominates SP1 on the one hand, and ZNF281 and NRF1 on the other, as candidate myeloid-specific and universal regulators (respectively) of the number of open rDNA repeats. Collectively, these findings demonstrate that the global chromatin landscapes of cells with different numbers of open rDNA repeats are characterized by differential enrichment of TF motifs, including both myeloid-specific and pan-essential TFs.

### Absolute quantification reveals an imbalance between mature 18S and 28S rRNAs

To expand beyond our relative quantifications of rRNA abundance in primary hematopoietic cell types, we next identified the absolute abundance of mature rRNA molecules per cell in each population. To do so, we first compared *in vivo* rRNA abundance to levels in myeloid cell lines. We used ER-HoxA9 as well as a related cell line, ER-HoxB8, immortalized by transgenic myeloid oncogenes HoxA9 and HoxB8 respectively^45^. We mixed cell lines with wild type primary BM spike-in and measured relative rRNA levels (Figure 7A; Supplemental Figure S9A). Nascent and mature rRNA were significantly higher in both cell lines relative to all normal and leukemic populations, with ER-HoxA9 cells exhibiting nascent rRNA levels 3.8x and mature rRNA levels 6.9x total BM median, while ER-HoxB8 cells displayed nascent rRNA levels 4.8x and mature rRNA levels 8.8x total BM median (Figure 7B; Supplemental Figure S9B-D). Notably, ER-HoxB8 cells proliferate slightly faster than ER-HoxA9 cells (Supplemental Figure S9E).

**Figure 7.**
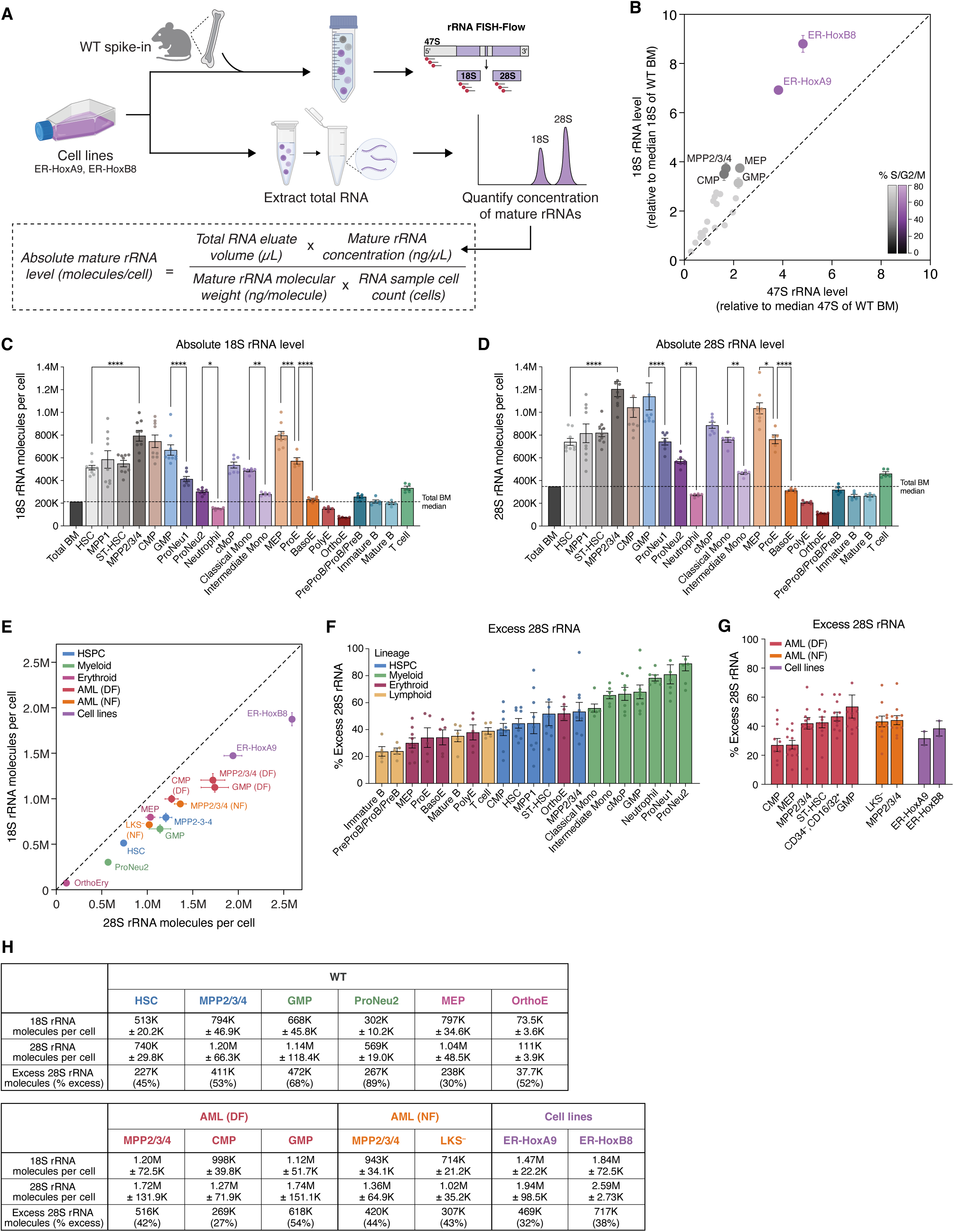
Absolute quantification reveals an imbalance between mature 18S and 28S rRNAs. **(A)** Experimental workflow using mouse cell lines to quantify absolute numbers of mature (18S and 28S) rRNA molecules per cell. **(B)** Comparison of relative nascent rRNA (47S rRNA) and ribosome subunit abundance (18S rRNA) in ER-HoxA9 and ER-HoxB8 cell lines and WT HSPCs (normalized to the median rRNA levels of total WT BM). The color of each cell type represents the percentage of cells in S/G2/M cell cycle phases. **(C-D)** Bar plots showing the absolute number of 18S (C) and 28S (D) rRNA molecules per cell across normal hematopoietic cell types. **(E)** Comparison of the average number of 18S and 28S rRNA molecules per cell across hematopoietic cell types. Values represent the average number of rRNA molecules per cell across all replicates per cell type. Cell types are colored by lineage and context. **(F)** Excess 28S rRNA molecules per cell compared to 18S. Each point represents the comparison of 28S and 18S rRNA abundance in paired samples from the same mice. Cell types are arranged in ascending order of 28S excess, and colored by lineage. **(G)** Same as in F, showing AML and cell line populations. **(H)** Table listing the average mature rRNA molecules per cell (range indicates ± SEM) and excess 28S rRNA molecules (number of molecules and percent excess) for select cell types (see Supplemental Fig S9H for all cell types). In some cases, replicates outside of the axis limits are not shown ( ≤ 2 replicates per cell type). All bar graphs and scatter plots show mean ± SEM. ns (*p* ≥ 0.05), * (*p* < 0.05), ** (*p* < 0.01), *** (*p* < 0.001), **** (*p* < 0.0001) by one-way ANOVA with Sidak’s multiple comparison testing.

Using these cell lines and their relative rRNA levels, we quantified the absolute number of mature rRNA molecules per cell (Figure 7A). We extracted total RNA from ER-HoxA9 and ER-HoxB8 cells, taking care to minimize sample loss at every step, and measured the concentration of 18S and 28S rRNA using a Bioanalyzer automated nucleic acid electrophoresis system. We used the known masses of mature rRNAs^46^ to calculate absolute numbers of 18S and 28S rRNA molecules per ER-HoxA9 and ER-HoxB8 cell. We then used relative MFI quantifications to calculate absolute mature rRNA levels in total bone marrow (Supplemental Figure S9F) and therefore in all primary normal and leukemic populations. 18S rRNA levels ranged from 73,500-797,000 molecules per cell across the normal hematopoietic system, and 714,000-1,200,000 across the AML hierarchy (Figure 7C,H; Supplemental Figure S9G-H). 28S rRNA levels similarly ranged from 111,000-1,200,000 molecules per cell in normal hematopoiesis and 920,000-1,740,000 in AML (Figure 7D,H; Supplemental Figure S9G-H). Importantly, these quantifications enabled direct comparison of 18S and 28S rRNA numbers. We observed a surprising imbalance in mature rRNAs, with 28S rRNA (large ribosome subunit) being more abundant than 18S rRNA (small ribosome subunit) in every cell type (Figure 7E; Supplemental Figure S9G-H). The ProNeu2 population, for example, had around 302,000 molecules of 18S rRNA per cell, yet over 569,000 molecules of 28S rRNA per cell. This imbalance remained apparent when comparing mature rRNA ratios of paired samples from the same mice (Figure 7F-H; Supplemental Figure S9H). 28S rRNA ranged from being in ∼20% excess (Immature B cell) to ∼90% excess (ProNeu2), indicating that post-transcriptional mature rRNA dynamics are both subunit-specific and cell-type-specific. Notably, the excess of 28S molecules was most strongly pronounced in the normal myeloid lineage (Figure 7F). We further confirmed the excess of 28S rRNA in primary cells by directly measuring mature rRNAs using Bioanalyzer automated electrophoresis quantification of RNA samples from WT and AML (DF) total BM (Supplemental Figure S9I). These strong, lineage-specific trends further support that regulation of rRNA dynamics is a cell-type-specific property.

### Reduced rRNA transcription promotes myeloid progenitor differentiation

To determine the effect of altered rRNA transcription on cell behavior and identity, we used the ER-HoxA9 Pol I degron cell line described in Figure 1 to achieve a range of reduced Pol I levels, and thus nascent and mature rRNA levels. Pol I degradation led to dose-dependent decreases in proliferation evident within 48 hours following dTAG treatment (Figure 8A-B), accompanied by progressive accumulation of cells in G1 (Supplemental Figure S10A). These effects occurred in the absence of substantial cell death, as viability remained high across treatment conditions and declined only modestly after prolonged treatment with 400 nM dTAG (Figure 8C). This indicates that, in this system, reduced rRNA synthesis primarily slows proliferative kinetics rather than triggering overt cell death. This could reflect the fact that the ER-HoxA9 cell line is p53-null (Supplemental Figure S10B), and therefore lacks the ability to stabilize p53 in response to nucleolar stress. Pol I degradation also caused an expected reduction in cell size by 48 hours as measured by forward scatter area (FSC-A) (Figure 8D).

**Figure 8.**
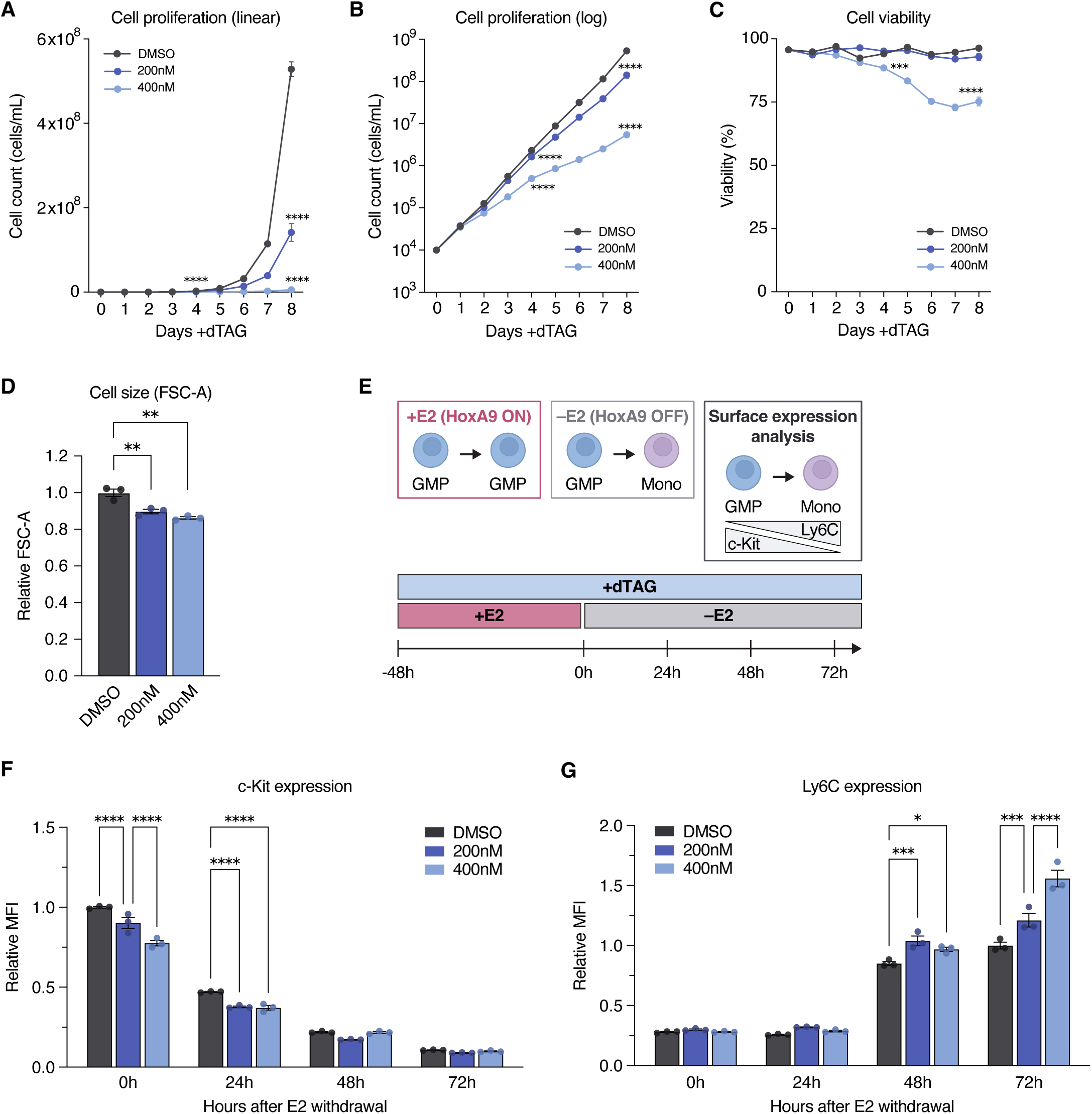
Reduced rRNA transcription promotes myeloid progenitor differentiation. **(A-B)** Proliferation of Pol I degron cells treated with DMSO or the indicated doses of dTAG. Cell counts shown on a linear (A) or log (B) scale. Comparisons by one-way ANOVA with Sidak’s multiple comparison testing. **(C)** Viability of Pol I degron cells treated with DMSO or the indicated doses of dTAG. Comparisons by one-way ANOVA with Sidak’s multiple comparison testing. **(D)** Relative cell size (measured using FSC-A) of Pol I degron cells treated with DMSO or the indicated doses of dTAG for 48 hours. **(E)** Schematic of the conditional ER-HoxA9 differentiation mechanism and estradiol (E2) withdrawal time course. Supplementation with E2 (+E2) maintains the cells in a myeloid progenitor state, and E2 withdrawal (–E2) inactivates HoxA9 and triggers myeloid differentiation. Pol I degron cells were pretreated with dTAG for 48 hours under +E2 growth conditions (0h), followed by E2 withdrawal and continued dTAG treatment during a 72 hour differentiation time course. Cell identity was assessed every 24 hours using flow cytometry for surface expression of the progenitor marker c-Kit and the differentiation marker Ly6C. **(F)** Relative c-Kit expression (MFI) throughout the E2 withdrawal time course in Pol I degron cells treated with DMSO or the indicated doses of dTAG, normalized to the average expression in DMSO replicates at the 0h timepoint. Comparisons by two-way ANOVA with Sidak’s multiple comparison testing. **(G)** Relative Ly6C expression (MFI) throughout the E2 withdrawal time course in Pol I degron cells treated with DMSO or the indicated doses of dTAG, normalized to the average expression in DMSO replicates at the 72h timepoint. Comparisons by two-way ANOVA with Sidak’s multiple comparison testing. All bar graphs, growth curves, and viability measurements show mean ± SEM. ns (*p* ≥ 0.05), * (*p* < 0.05), ** (*p* < 0.01), *** (*p* < 0.001), **** (*p* < 0.0001). n = 3 replicates per condition.

The ER-HoxA9 line was generated by immortalizing mouse bone marrow with the myeloid oncogenic transcription factor HoxA9 fused to an estrogen receptor (ER) domain. When cultured with beta-estradiol (E2), a HoxA9 transcriptional program maintains the cells in a myeloid progenitor state, while E2 withdrawal inactivates HoxA9 and triggers myeloid differentiation. To assess whether graded reductions in rRNA transcription influence differentiation dynamics, we pretreated Pol I degron cells with 200 nM or 400 nM dTAG for 48 hours under +E2 growth conditions, followed by E2 withdrawal and continued dTAG treatment during a 72 hour differentiation time course (Figure 8E). We assessed cell identity using flow cytometry for surface expression of the progenitor marker c-Kit and the differentiation marker Ly6C. Cells not treated with dTAG rapidly downregulated c-Kit and progressively upregulated Ly6C following E2 withdrawal, reflecting their normal progression through the differentiation program (Figure 8F-G). In contrast, dTAG-treated cells showed dose-dependent reduction in c-Kit expression before E2 withdrawal (Figure 8F). After E2 withdrawal, dTAG-treated cells upregulated Ly6C expression more rapidly and to a greater extent than untreated cells (Figure 8G). Notably, we validated that the per-cell abundance of the ER-HoxA9 protein remained unchanged, demonstrating that these shifts in cell identity cannot be attributed to reduction in the transgenic fusion protein (Supplemental Figure S10C). Together, these findings indicate that reduced rRNA transcription induces and accelerates myeloid progenitor differentiation, supporting an active role for cell-type-specific levels of rRNA transcription and ribosome subunits in maintenance of cell identity.

## DISCUSSION

Despite the fact that rRNAs are the most abundant RNAs in the cell, their dynamics within normal and disordered organ systems are surprisingly understudied. Here, we used an optimized rRNA FISH-Flow assay to precisely quantify nascent rRNA (surrogate for transcription) and mature rRNA (ribosome subunit) abundance in distinct cell types within primary mouse bone marrow, enabling analysis across contexts of hematopoiesis. Our detailed profile of normal hematopoiesis reveals that within the nearly 10-fold variation in nascent and mature rRNA abundance across cell types, rRNA levels are a cell-type-specific property.

HSCs have been previously shown to actively engage in ribosome biogenesis^19^ despite low cycling and global protein synthesis rates^15,16^. In agreement with this result, we found that nascent rRNA levels in HSCs are above the median of total BM. This is surprising because mature ribosome subunits are reported to have half-lives ranging in days^47,48^, and HSCs, which are in G0 and cycle rarely (no more than once every few months)^49,50^, would thus be expected to require only minimal new rRNA transcription to maintain a stable ribosome pool in the absence of constant dilution to daughter cells. One possible explanation for their non-low levels of nascent rRNA is that mature subunits may have particularly short half-lives in HSCs. Bulk ribosome subunit breakdown can occur in autophagosomes in a process known as ribophagy^51–53^. HSCs are known to have high levels of generalized and selective autophagy in order to maintain quiescence and self-renewal capacity^54–56^. It is conceivable that rapid ribosome subunit degradation and constant synthesis of new ribosomes may also be essential for stemness; we hypothesize that accumulation of “aged” and error-prone ribosomes^52,57–59^ may be detrimental to HSCs for reasons of translation fidelity and protein quality control.

In addition, HSCs maintain a fairly high number of mature ribosome subunits despite having very low rates of translation. A plausible explanation is that HSCs must be poised to rapidly enter the cell cycle and increase translation upon cytokine signaling. Maintaining a high number of inactive ribosome subunits, ready to engage in translation when needed, likely helps HSCs achieve this state of poised quiescence.

In HSCs and across the hematopoietic tree, we identified a surprising imbalance in mature rRNA (28S versus 18S) abundance. This imbalance, as well as the mismatch between relative mature and nascent rRNA levels, indicates that steady-state ribosome subunit numbers are determined by more than just nascent rRNA synthesis alone. Instead, they are likely shaped either by downstream steps of ribosome assembly and subunit maturation, or more likely by mature ribosome subunit half-lives. The variable degree of imbalance across cell types further suggests that rRNA half-lives may be cell-type-specific. In addition to bulk ribophagy pathways, ribosome subunits can be targeted for degradation through selective ribosome surveillance and degradation pathways. In yeast, 40S and 60S subunit turnover are well established to occur through subunit-specific degradation pathways^52,60^, and growing evidence suggests similar pathways exist in mammalian cells^61–63^. Differential regulation of these turnover pathways could therefore contribute to the mature rRNA mismatches observed across cell types and help explain dynamic cell-type-specific patterns of nascent versus mature rRNA abundance across hematopoiesis. The pronounced excess of 28S rRNA (60S subunit) in the normal myeloid lineage hints at a possible functional relevance of this imbalance in myeloid development or biology, potentially reflecting greater asymmetry in 40S versus 60S subunit degradation activity. Future experiments will be required to define the mechanisms and consequences of these lineage-specific rRNA imbalances.

We tested the functional significance of cell-type-specific rRNA abundance by generating a myeloid progenitor Pol I degron cell line to achieve tunable control of rRNA abundance, where we identified that graded reduction in rRNA transcription compromises progenitor identity and accelerates differentiation. This suggests that rRNA transcription and ribosome subunit levels actively contribute to maintenance of cell identity, and we speculate that this may occur in part through regulation of cell-type-specific translation dynamics. In addition to supporting global translational capacity, ribosome numbers likely also shape gene-specific translation landscapes, in line with the Ribosome Concentration Hypothesis^64^. Such a model, extended more broadly to the hematopoietic system, would be an attractive explanation for why different cell types maintain different numbers of ribosomes. Further testing of this model will help quantify the degree to which specific cellular proteomes are shaped by ribosome abundance in both physiological and pathological contexts.

We identified that broad patterns of nascent rRNA might be partly explained by trends in core rDNA transcription machinery gene expression and rDNA accessibility. How these regulatory processes are differentially fine-tuned between cell types remains incompletely understood. Transcription of some components of the core rDNA machinery is known to be promoted by MYC^29,30^, but that cannot exclusively explain rRNA transcription in HSCs and MPPs; these populations are preserved or even increased in number in hematopoietic MYC knockout mouse models even as downstream hematopoiesis is eliminated^65,66^. Additional TFs beyond MYC likely control abundance of rDNA machinery, warranting future cell-type-specific dissection. Furthermore, differences in abundance of core rDNA machinery do not fully explain variations in nascent rRNA within stem and progenitor populations. Our previous work established that numerous hematopoietic lineage-specific TFs localize to rDNA^17^. It is possible that these factors locally control rRNA transcription independent of the abundance of core machinery, potentially by fine-tuning rDNA accessibility, assembly of the Pol I Preinitiation Complex at the rDNA promoter, or Pol I processivity. Along these lines, we had also shown that loss of the myeloid TF CEBPA rapidly reduces Pol I occupancy on rDNA without affecting the overall abundance of any components of rDNA machinery^17^; other cell-type-specific TFs may assume similar regulatory roles.

In this current work, through our analyses of rDNA accessibility, we found that genome-wide chromatin accessibility landscapes differ between cells with greater versus lesser number of active rDNA copies. Cells with higher copy activation showed increased accessibility of regions enriched with SP1 and NRF1 motifs, indicating increased levels or activity of these TFs. SP1 is a broadly expressed TF with critical roles in embryonic hematopoietic specification and maintenance across multiple lineages^67,68^. Given this established role in normal hematopoietic development as well as its strong and selective dependency in AML cell lines, SP1 emerges as a compelling candidate for further study. NRF1, on the other hand, is a ubiquitously expressed TF involved in diverse cellular pathways, and its mouse knockout leads to mid-embryonic lethality due to impaired fetal liver hematopoiesis^69,70^. It is a dependency across all cell lines, nominating it as a possible universal regulator of rDNA copy activation. For SP1, NRF1, as well as other TFs we identified as being associated with high rDNA repeat accessibility, additional investigation will be required to disentangle correlation from causation.

In our analyses of rRNA levels in AML, we focused on mouse models driven by combined *Dnmt3a*;*Flt3*^ITD^ or *Npm1*^c^;*Flt3*^ITD^ mutations, which recapitulate common genetic subtypes of human AML. AML cells exhibited elevated nascent and mature rRNA levels compared to normal HSPC counterparts, which bolsters the longstanding idea that rRNA transcription, and ribosome biogenesis more broadly, may represent therapeutically targetable vulnerabilities in AML^71,72^. While additional investigation is needed to more comprehensively resolve the mechanisms that upregulate rRNA levels in AML, we identified that the number of active rDNA copies is increased in AML and may contribute to elevated nascent rRNA. Strikingly, leukemia stem cell (LSC) populations appeared to have higher rDNA activation than the rest of the leukemic blast population, which may indicate a selective therapeutic vulnerability relative to normal HSCs. Together, these findings raise the possibility that dysregulated rRNA pathways may represent clinically relevant vulnerabilities in human AML, motivating future studies to define their relevance across additional genetic subtypes and their potential association with therapy response, relapse risk, or clinical risk stratification.

We acknowledge certain limitations to our analyses. First, our 47S rRNA FISH probes hybridize to the 5’ tip of the nascent 47S pre-rRNA transcript (which is rapidly cleaved during the first steps of pre-rRNA processing), and it is possible that the probes also capture truncated transcripts or transcripts undergoing delayed processing. Thus, variation in early rRNA processing dynamics may contribute to our 47S rRNA FISH-Flow quantifications. Future studies focused on rRNA processing, including emerging Nanopore RNA sequencing approaches^73^, will help further resolve cell-type-specific rRNA dynamics. In addition, we accounted for differences in proliferation dynamics between populations by staining with DAPI to determine the percentage of cells in S/G2/M phases. However, this is not a direct measure of proliferation rate or doubling time, and therefore we cannot directly associate rRNA levels with proliferation kinetics. For our AML analyses, additional profiling of rRNA levels in non-malignant models of rapidly proliferative hematopoiesis will address the extent to which higher nascent and mature rRNA levels in AML are consequences of malignant identity versus rapid proliferation. Finally, further investigation using additional mouse models and human patient samples will assess the generalizability of our finding of elevated rRNA abundance in AML.

In summary, our study reveals that rRNA synthesis and ribosome biogenesis exhibit distinct cell-type-specific patterns across normal hematopoiesis. Moreover, our finding that rRNA transcription level actively contributes to cell identity positions rRNA pathways as dynamic regulators of cell fate and function.

## METHODS

### Cell lines and cell culture

#### Cell culture conditions

ER-HoxA9 and ER-HoxB8 mouse cell lines were a generous gift from Andres Blanco (University of Pennsylvania) and David Sykes (Massachusetts General Hospital). Cell lines were cultured at 37° C in a 5% CO_2_ incubator using RPMI-1640 media (Invitrogen, 22400071) containing 10% fetal bovine serum (GeminiBio, 100-106), 2% stem cell factor (SCF) conditioned media (prepared from a Chinese hamster ovary cell line that stably secretes SCF), 1% Penicillin/Streptomycin (Invitrogen, 15140122), and 50 µM β-estradiol (Fisher Scientific, MP021016562). For estradiol withdrawal (–E2) experiments, cells were cultured in the same media, without β-estradiol. Aside from E2 withdrawal experiments, all experiments were performed in media containing E2. Cells were cultured at concentrations between 10K-800K cells/mL for all experiments. For Pol I degron proliferation measurements, cells were plated at 10K cells/mL and treated with the indicated doses of DMSO (Sigma Aldrich, D2650) or dTAG (Tocris, 6914), counted every 24 hours, and passaged with fresh media with replenished dTAG every 72 hours. For ER-HoxA9 and ER-HoxB8 proliferation measurements, cells were plated at 20K cells/mL, counted every 24 hours, and passaged with fresh media every 48 hours. All cell counts for harvests, proliferation, and viability measurements were performed using a BD Accuri C6 Flow Cytometer (BD Biosciences).

#### Generation of Polr1a degron cell line

The *Polr1a* (Pol I) degron cell line was generated by using CRISPR HDR as previously described^17^ to integrate an mScarlet-P2A-FLAG-FKBPV cassette into the N-termini of endogenous *Polr1a* alleles in the ER-HoxA9 cell line. Briefly, a guide RNA targeting the region near the *Polr1a* start codon was complexed with Cas9 and electroporated into ER-HoxA9 cells together with a donor plasmid containing the mScarlet-P2A-FLAG-FKBPV degron cassette flanked by 450bp homology arms. Single-cell clones were isolated, expanded, and screened for biallelic integration by PCR. The selected clone was subsequently validated by immunoblotting to confirm successful tagging and degradation upon treatment with dTAG.

#### Immunoblotting

For blots detecting POLR1A protein expression (anti-RPA194 and anti-FLAG), Pol I degron cells were plated at 30K cells/mL 24 hours prior to dTAG treatment to allow for recovery, and cells were then treated with DMSO or the indicated doses of dTAG for 24 hours. 2 x 10^6^ cells were centrifuged at 300 x *g* for 5 minutes at 4° C and washed in cold PBS. Pelleted cells were lysed in 45 μL lysis buffer [200 μL 5X RIPA buffer, 20 μL PIC (Sigma Aldrich, P8340), 20 μL PMSF (Cell Signaling Technology, 8553S), 20 μL Chymostatin (Sigma Aldrich, EI6), 10 μL PI (Sigma Aldrich, P0044), 1 μL Benzonase (Millipore, E1014), 729 μL nuclease free water] for 1 hour on ice. Samples were then prepared in 4x Laemmli (BioRad, 1610747). For blots detecting the ER-HoxA9 fusion, Pol I degron cells were plated at 10K cells/mL and treated with DMSO or the indicated doses of dTAG for 72 hours. 5 x 10^6^ cells were centrifuged at 300 x *g* for 5 minutes at 4° C and washed twice in cold PBS. Pelleted cells were lysed in 100 μL lysis buffer [400 μL SDS dye, 50 μL BME, 20 μL PIC, 20 μL PMSF, 20 μL Chymostatin, 490 μL nuclease free water] and immediately boiled at 95° C for 5 minutes. Samples were then sonicated for 2 sets of 5 cycles (30 seconds on, 30 seconds off) using a Bioruptor machine (Diagenode).

Prepared samples were electrophoresed in NuPAGE 4%-12% Bis-Tris polyacrylamide gels (Invitrogen), and wet transferred overnight at 24V to PVDF membranes (Millipore). Membranes were blocked for 1 hour in 5% nonfat milk in TBST and incubated at 4° C overnight in primary antibody, followed by 1 hour incubation in fluorescent secondary antibody. Images were captured using an Odyssey CLx imaging system (LI-COR Biosciences), and protein quantification was carried out using ImageJ software (Fiji). For all blots, samples were prepared and loaded using equal cell numbers to enable bulk per-cell quantification. For measurements of Pol I abundance, blots for anti-FLAG were used for quantification.

Primary antibodies and concentrations used were: anti-RPA194 1:500 (Santa Cruz Biotechnology, sc-48385), anti-FLAG-M2 1:1000 (Sigma, F1804), anti-ERα 1:500 (Santa Cruz Biotechnology, sc-8002), anti-αTubulin 1:2000 (Cell Signaling Technology, 3873S), anti-GAPDH 1:2000 (Cell Signaling Technology, 2118S). Secondary antibodies and concentrations used were: anti-rabbit 680RD 1:1000 (LI-COR Biosciences, 925-68073), anti-mouse 800CW 1:1000 (LI-COR Biosciences, 925-32212).

#### Northern blotting

For Northern blot analyses, ER-HoxA9 and ER-HoxB8 cells were plated at 80K cells/mL and cultured for 24 hours, and primary wild type and AML (DF) total bone marrow cells were freshly isolated following mouse euthanasia. Samples of 10 million cells were harvested for total RNA extraction using the TRIzol-Chloroform method. In brief, cells were spun down at 300 x *g* for 5 minutes at RT, washed once in cold PBS, and pellets were then resuspended in 100 µL PBS and 1 mL of TRIzol (Invitrogen, 15596018) and mixed by pipetting 25 times to ensure complete lysis and release of all RNA. 200 µL of chloroform (FisherScientific, BP1145-1) was added to each sample, and samples were mixed vigorously by hand. Samples were then incubated at RT for 3 minutes and spun down at 12,000 x *g* at 4° C for 15 minutes. The top aqueous layer containing the RNA was transferred to a new tube, ensuring that the entirety of the aqueous layer was collected. Samples were mixed with 500 µL of isopropanol (Fisher Scientific, A461-500), incubated at RT for 10 minutes, and spun down at 12,000 x *g* at 4° C for 10 minutes. RNA pellets were then resuspended in 1 mL of 75% ethanol, mixed by brief vortexing, and spun down at 7,500 x *g* at 4° C for 5 minutes. Pelleted RNA was then allowed to air dry until clear, and was resuspended in 50 µL of nuclease free water. RNA samples were further purified through column purification (Zymo Research R1013-2, R-1003).

Northern blot analysis was performed as previously described^74^ with minor changes. In short, 10 ug of RNA was denatured for 10 min at 95° C in 1x RNA loading dye [1x MESA, 13.6% formaldehyde, 6.4% formamide, bromophenol blue, and xylene cyanol] and separated on a 1.25% denaturing agarose gel containing 1x MESA and 6.7% formaldehyde. The gel was run in 1x MESA at 100 V at 4° C for 17 hours. Capillary transfer was performed onto a Hybond N+ membrane in 10x SSC. After UV cross-linking, the membrane was hybridized with the 5’-[^32^P]-labeled-DNA oligonucleotide probe (5’ to 3’: AGAGAAAAGAGCGGAGGTTCGGGACTCCAA) at 65° C overnight, washed (2 x 10 minutes) at 37° C, and exposed to a phosphorimager screen. The signal was detected using a Typhoon FLA 9500 (GE Healthcare) phosphoimager and quantified using ImageJ software (Fiji).

#### 5-EU incorporation assays

5-EU incorporation assays were performed using the Invitrogen Click-iT™ RNA Alexa Fluor™ 488 Imaging Kit (Thermo Scientific C10329) with minor protocol modifications. Poly-L-lysine coated glass slides (Sigma Aldrich P0425) were prepared by first drawing a 2 cm diameter circular staining area on the upper slide surface using a hydrophobic pen (Fisher Scientific, NC9827128), and then adding 300 µL retronectin and incubating for 4 hours at 37° C. Retronectin was removed, slides were air dried for 10 minutes, and 500K Pol I degron cells in media containing either DMSO or 400 nM dTAG were added to each slide and incubated for 24 hours at 37° C to adhere to the slides during dTAG treatment. The following day, cells were incubated with dTAG and 1mM 5-EU for 30 minutes at 37° C, washed with 300 µL PBS, and fixed in 300 µL 4% paraformaldehyde in PBS for 20 minutes at room temperature. Following fixation, cells were washed twice in 300 µL PBST [50 mL of DPBS + 250 µL of 10% Tween 20] and permeabilized with 300 μl of 0.5% (v/v) Triton X-100 (Sigma-Aldrich, 9036-19-5) in PBS for 15 minutes. After two additional washes with PBST, cells were incubated with 300 μl of blocking buffer [9 mL of DPBS + 1 mL goat serum (Millipore Sigma F0926) + 200 µL 10 % Tween 20], washed twice in 300 µL PBST, and incubated overnight at 4° C with primary anti-nucleolin antibody (1:100, Abcam ab22758) prepared in blocking buffer. Cells were then washed twice in PBST and incubated for 1 hour with 10 µg/mL Hoechst 33342 and AF647 anti-rabbit secondary antibody (1:250, Invitrogen A-21244) prepared in blocking buffer. Cells were washed once and incubated for 30 minutes in 300 µL AF488 Click-iT reaction cocktail freshly prepared according to manufacturer instructions. Following the Click-iT reaction, cells were washed once in Click-iT rinse buffer, washed twice in PBST, and coverslips were mounted using ProLong Gold Antifade reagent (Invitrogen, P36930) and sealed using clear nail polish.

Images were acquired using a Leica DM6000 widefield microscope with a 63x objective, and analyzed using Cell Profiler (v4.2.8). For image analysis, per-cell nucleolar 5-EU intensity was calculated by subtracting background nucleoplasmic 5-EU signal from the total nucleolar 5-EU intensity, as defined below. Nucleoli were identified based on nucleolin signal. Two replicate slides were imaged for each condition, and violin plots display pooled quantifications from approximately 250 cells per replicate (∼500 total cells per condition).

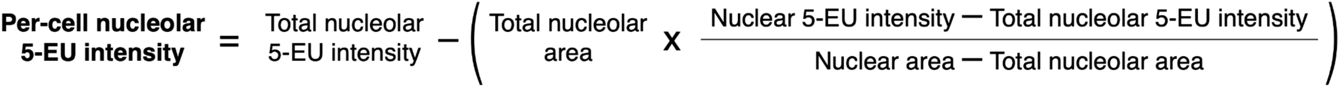

#### Sucrose gradient fractionation

Pol I degron cells were plated at 30K cells/mL 24 hours prior to dTAG treatment to allow for recovery, and cells were then treated with DMSO or 400 nM dTAG for 24 hours. Cells were then counted using a BD Accuri C6 Flow Cytometer (BD Biosciences), and 2.5-4 x 10^6^ cells were harvested and washed with ice cold PBS. Pelleted cells were lysed in 250 μL of ribosome lysis buffer [20 mM Tris-Cl pH 7.5, 140 mM KCl, 1% Triton-X-100, 0.5 mM DTT] and incubated on ice for 10 minutes. Lysate was clarified by centrifugation at 17,000 x *g* for 10 minutes. To dissociate all ribosomes into free ribosomal subunits, the supernatant was treated with EDTA (final concentration 60 mM) for 30 minutes on ice. 190 μL of EDTA-treated lysate was layered on top of 7-30% sucrose gradient [20 mM Tris-Cl pH 7.5, 140 mM KCl, 5 mM EDTA, 0.5 mM DTT] prepared using a gradient station (Biocomp instruments) and ultracentrifuged at 40,000 RPM for 2 hours at 4° C in a SW60Ti rotor (Beckman). After centrifugation, gradients were analyzed using a fractionator equipped with a Triax™ UV flow cell (Biocomp instruments), and the A_260_ absorbance profile was recorded. Total 40S and 60S subunit abundances were quantified by calculating the area under the curve of corresponding peaks using ImageJ software (Fiji). For each replicate, quantifications from dTAG-treated cells were normalized to the paired DMSO control that was processed in parallel, with the corresponding DMSO value set to 1.

#### Pol I degron estradiol withdrawal and flow cytometry

For the estradiol withdrawal time course, Pol I degron cells were plated at 30K cells/mL in media containing estradiol (+E2) and treated with the indicated doses of DMSO or dTAG. After 48 hours of culture in +E2 media with dTAG, E2 was withdrawn by washing 1.5 million cells per sample 3 times in PBS. Cells were then resuspended in –E2 media containing the indicated doses of DMSO or dTAG and split across plates for subsequent time points. At each time point, 250K cells per sample were harvested and washed with cold PBS. Cells were stained for 30 minutes on ice in the dark with an antibody cocktail containing anti-c-Kit BV605 (1:100) and anti-Ly6C BV785 (1:100) diluted in cell staining buffer (BioLegend, 420201). Cells were then washed twice in 100 μL cell staining buffer, resuspended in 300 μL cell staining buffer, and analyzed on a FACSymphony A3 cytometer (BD Biosciences). Flow cytometry data were analyzed using FlowJo version 10.10.0 (BD Biosciences). Marker expression was quantified as the median fluorescence intensity (MFI) of the indicated marker within total singlets.

#### Mice

Wild-type C57BL/6J mice (JAX# 000664) were aged 8-20 weeks prior to experimental use. C57BL/6J CD45.1 mice (JAX# 002014) aged 8-20 weeks were utilized for normalization spike-in for AML (DF) and AML (NF) bone marrow. For the AML (DF) transgenic model, *Mx-Cre;Flt3^ITD^*;*Dnmt3a^fl^*mice were generously gifted by Sara Meyer (Thomas Jefferson University), Leighton Grimes (Cincinnati Children’s Hospital Medical Center), and Martin Carroll (University of Pennsylvania). Mice with the genotype *Mx-Cre/+;Flt3^ITD/ITD^;Dnmt3a^fl/fl^*became clinically moribund from AML at around 5 weeks of age, at which point they were euthanized for experimental use. For the AML (NF) transgenic model, *Rosa26*^Flpo-ERT2/wt^;*Npm1*^Frt-c/wt^;*Flt3*^Frt-ITD/wt^ mice had previously been induced to develop overt AML as described^27^, and bone marrow had been harvested and frozen. We reconstituted NF AML *in vivo* by thawing and transplanting these pre-frozen samples. Eight-week-old female C57BL/6J homozygous CD45.1 mice were lethally irradiated (1100 cGy total body irradiation administered in two fractions of 550 cGy), and then transplanted via retro-orbital injection with 2 x 10^5^ freshly-thawed NF AML cells and 4 x 10^5^ heterozygous CD45.1/CD45.2 support cells. Engraftment was assessed by flow cytometry analysis of CD45.2^+^ CD45.1^-^ leukemic cells in the peripheral blood. Mice were euthanized when engraftment exceeded 50% or when they became clinically moribund. All housing and husbandry was carried out under the guidance of the University of Pennsylvania’s Animal Resources Center and followed approved IACUC protocol guidelines. Equal numbers of male and female mice were used for all analyses.

#### Generation of rRNA FISH-Flow probe sets

rRNA FISH-Flow probe oligo sequences were previously published^23^ and were purchased from Integrated DNA Technologies (IDT). Probe conjugation with Quasar 670 dye (Biosearch Technologies, FC-1065S-5) and HPLC purification was carried out as previously described^75^. Probe pellets were resuspended in TE buffer at a concentration of 12.5 µM.

#### rRNA FISH-Flow on the Pol I degron cell line

For rRNA FISH-Flow experiments involving the Pol I degron cell line, cells were plated at 30K cells/mL 24 hours prior to dTAG treatment to allow for recovery, and cells were then treated with the indicated doses of dTAG or DMSO (equivalent to the highest volume of dTAG) for 24 hours. For each sample, 500K cells were harvested, fixed in 4% paraformaldehyde in PBS (Thermo Scientific, J19943-K2), and rRNA FISH-Flow was carried out as previously described^23^. To normalize across samples, a 40% spike-in of fixed parental ER-HoxA9 cells was added to degron samples during the first step of setting up FISH-Flow reaction plates. For rRNA MFI quantification of each sample, the Pol I degron (mScarlet^+^) rRNA MFI was internally normalized to the spike-in ER-HoxA9 (mScarlet^-^) rRNA MFI to obtain the final normalized MFI. Pol I degron cell cycle quantifications were obtained from DAPI profiles derived from rRNA FISH-Flow samples.

#### rRNA FISH-Flow on primary bone marrow

Mouse femurs, tibias, fibulas, and hips were isolated, soaked in PBS-EDTA [DPBS (Gibco, 14190-136), 0.5% EDTA (Invitrogen, 15575038)], and crushed with a mortar and pestle. The resultant slurry was passed through a 40 µm cell strainer (Fisher Scientific, 22-363-547) and placed immediately on ice without ACK lysis to minimize cell stress. Samples of 20 million cells were stained with 200 µL of a surface marker antibody cocktail (all antibodies at 1:100 dilution). All remaining steps were carried out protected from light. Samples were then spun down and fixed in 5 mL of 4% paraformaldehyde in PBS under gentle rocking at room temperature (RT) for 10 minutes. No additional permeabilization step was required, as we found that paraformaldehyde alone achieved adequate permeabilization for FISH probes. Following fixation, samples were washed once with 500 µL PBS-EDTA and once with 500 µL wash buffer [40 mL nuclease-free water (Fisher, BP561-1), 5 mL 20X SSC (Invitrogen, AM9770), 5 mL formamide (Invitrogen, AM9342)]. Samples were then resuspended in 100 µL of hybridization buffer [8 mL nuclease-free water, 1 mL 20X SSC, 1 mL formamide, 1 g dextran sulfate (Sigma-Aldrich, D5906-50G)] containing either 20 µL 47S rRNA probe pool, 10 µL 18S rRNA probe pool, or 10 µL 28S rRNA probe pool, and incubated at 30° C for 3 hours. Following hybridization, 1 mL of wash buffer was added directly to the hybridization buffer suspension, and samples were spun down at 1000 x *g* for 5 minutes. Pellets were resuspended in 500 µL of wash buffer and incubated at 30° C for 30 minutes to remove unbound probe. Samples were then spun down at 1000 x *g* for 5 minutes, resuspended in 500 µL of 4’,6-diamidino-2-phenylindole (DAPI) (Fisher Scientific, D1306; 4 ng/µL concentration prepared in wash buffer), and incubated at 30° C for 30 minutes. Cells were washed once in 500 µL of 2X SSC buffer [45 mL nuclease free water, 5 mL 20X SSC], resuspended in 300 µL of cell staining buffer, and data were acquired using an LSR Fortessa cytometer (BD Biosciences) or FACSymphony A3 cytometer (BD Biosciences). Nascent (47S) and mature (18S, 28S) rRNA intensities were recorded in separate experiments. For samples with fewer than 20 million cells available, all volumes for staining and washes were scaled down accordingly.

For all AML (DF) and AML (NF) samples, a 10% spike-in of wild type (WT) CD45.1 BM cells was added to enable normalization of AML rRNA levels to the median of total WT BM. For AML (DF) samples, the spike-in was added prior to surface staining because all AML mouse-derived cells were CD45.1^-^. The combined cell mixture was then stained with anti-CD45.1 (PE) to distinguish WT spike-in cells from AML cells. For AML (NF) samples, the spike-in was added after surface staining and washing, immediately prior to fixation. This was necessary because any residual WT recipient or support cells would also be CD45.1^+^. Accordingly, AML mouse-derived cells were stained with anti-CD45.1 (BV785) to identify and exclude any endogenous CD45.1^+^ WT recipient or support cells, and the WT CD45.1 spike-in was separately stained with anti-CD45.1 (PE).

To quantify relative rRNA MFIs in ER-HoxA9 and ER-HoxB8 cell lines, clonally derived lines were plated at 180K cells/mL 24 hours prior to harvest to allow optimal health and exponential growth. 5 million ER-HoxA9 or ER-HoxB8 cells and 5 million freshly harvested WT mouse bone marrow cells were separately stained with anti-CD45.2 antibody in two different colors (PE and AF700). After washing away unbound antibody, stained cells were mixed and immediately fixed together and subjected to the full FISH-Flow protocol to enable normalization of rRNA levels in cell lines to the median of total WT BM.

### Flow cytometry analysis and normalization

#### Normal hematopoiesis

Samples consistently exhibited > 95% cells with positive rRNA FISH-Flow probe staining for all three rRNA species. Eosinophils (<5% of bone marrow cells) were excluded due to their aberrantly high signal with even nonspecific (anti-GFP-mRNA) probes (Supplemental Figure S2). Initial gating steps were consistently applied to all samples as follows: DAPI profile was first used to exclude all sub-G1 (dead) cells, followed by live cell gating using FSC-A vs SSC-A. All doublets and eosinophils (high rRNA probe, high SSC-A) were then excluded. The CD45^+^ population was next defined for use as internal normalization of rRNA probe fluorescence within each sample. Gating strategies for each cell type were then applied as listed in Supplemental Figure S3 and Supplemental Table S1. For each sample, the gate defining cell cycle phase subpopulations (G0/G1 and S/G2/M) was manually applied based on the DAPI profile of the CD45^+^ population. All average % S/G2/M distributions for each cell type were calculated based on replicates from 47S rRNA FISH-Flow experiments. rRNA level in each cell type was quantified by extracting the median fluorescence intensity (MFI) of the rRNA probe, and normalized by dividing by the rRNA MFI of the CD45^+^ population (total BM) of that sample. We defined a minimum cutoff of at least 50 cells for the rarest cell populations to obtain a reliable rRNA probe MFI, though most MFIs were obtained from tens of thousands of events within each population. All flow data were analyzed using FlowJo version 10.10.0 (BD Biosciences).

#### AML

Initial gating steps were applied to all AML (DF) and AML (NF) samples as described above, except that spiked-in CD45.1^+^ BM cells were used for normalization. Gating of AML stem and progenitor-like populations was guided by normal BM profiles and unstained fluorescence controls. Gating strategies are listed in Supplemental Figure S6 and Supplemental Table S1.

rRNA MFIs were extracted from each AML population and internally normalized to the rRNA MFI of the CD45.1^+^ (total WT BM) spike-in. Given the slight known variation in BM composition between CD45.1 and CD45.2 mice, we also directly compared rRNA levels in BM from both mouse strains. To do so, BM cells from CD45.2 WT mice and CD45.1 WT mice were combined in a 1:1 ratio, stained with CD45.2 (AF700) and CD45.1 (PE), and processed through all remaining steps of the rRNA FISH-Flow assay. Ratio of total BM rRNA MFIs from CD45.2^+^ mice compared to CD45.1^+^ mice were found to be 1.17 (47S), 1.09 (18S), and 1.13 (28S) (n = 4-8 mouse pairs per probe) (Supplemental Figure S7A). These (small) correction factors were applied to the spike-in-normalized AML rRNA MFIs by dividing each normalized MFI by the corresponding correction factor.

#### Cell lines

Cell line (PE CD45.2^+^) and primary BM spike-in (AF700 CD45.2^+^) gating was used to measure rRNA MFIs separately from each population, and cell line MFIs were internally normalized to the CD45.2^+^ primary BM (Supplemental Figure S9A).

#### Nonspecific rRNA FISH-Flow probe background signal

Negative control anti-GFP-mRNA probes were used to determine the threshold of nonspecific FISH probe binding in non-eosinophil cells. WT bone marrow cells were hybridized with an equal concentration of AF594 anti-GFP-mRNA probe and rRNA probe, and the fluorescence of both probe sets in the non-eosinophil population was measured. Nonspecific background signal was calculated as follows: (non-eos anti-GFP MFI/unstained anti-GFP MFI) / (non-eos rRNA MFI/unstained rRNA MFI). The nonspecific rRNA FISH-Flow probe background signal was determined to be 0.2 fold of the total BM signal for 47S rRNA, 0.04 fold of 18S rRNA, and 0.1 fold of 28S rRNA (n = 4-6 mice per rRNA probe) (Supplemental Figure S2B-E). We deemed these thresholds to be the lower limits of reliable quantification for each probe set.

### Absolute quantification of mature rRNAs

#### Quantification of ER-HoxA9 and ER-HoxB8 cell lines

Absolute quantification of mature rRNAs for calculating mature rRNA molecules per cell in primary cell types was performed using the ER-HoxA9 and ER-HoxB8 cell lines. To do so, cells were plated at 180K cells/mL 24 hours prior to harvest and were counted immediately before harvest using a BD Accuri C6 Flow Cytometer (BD Biosciences). From each flask, 15 million cells were harvested to be used for primary rRNA FISH-Flow as described above, and an additional two replicates of exactly 5 million cells were harvested for absolute quantification. Total RNA was extracted using the TRIzol-Chloroform method as described above (see Northern blotting methods). Care was taken at every step to avoid sample or pellet loss, and to ensure full reconstitution of all RNA in the final solution. For this reason, RNA samples for absolute quantification were not column purified.

Two dilutions (1:3 and 1:9) for each RNA sample were prepared in RNA ScreenTape Sample Buffer (Agilent, 5067-5577) as described in the Agilent RNA ScreenTape Assay for TapeStation Systems protocol, and run on an RNA ScreenTape (Agilent, 5067-5576) using the Agilent 4150 TapeStation System. Peaks corresponding to 18S and 28S rRNAs in the resulting electropherogram were manually adjusted to achieve consistent peak boundaries across samples, and the resulting concentrations of 18S rRNA and 28S rRNA (in ng/µL) were obtained from each corresponding peak. In total, two flasks were set up for each cell line (ER-HoxA9 and ER-HoxB8), with two RNA samples harvested from each flask, and two dilutions of each RNA sample quantified using the TapeStation system. All samples were analyzed on two separate TapeStation runs, for a total of 32 final absolute quantification measurements for each mature rRNA. Molecular weights^46^ of 18S rRNA (1.06 x 10^-9^ ng per molecule) and 28S rRNA (2.66 x 10^-9^ ng per molecule) were used for calculation of absolute number of mature rRNA molecules as listed in Figure 7A. Contribution of rRNA modifications to the molecular weight of each mature rRNA was deemed to be negligible. For all replicates from each flask, the rRNA absolute quantification value was divided by the relative rRNA MFI for the respective flask (obtained from FISH-Flow) to obtain the absolute quantification value corresponding to the total BM CD45^+^ (rRNA MFI = 1) population. Once replicates with RNA degradation (RIN < 8) were excluded, there were a total of 29 independent measurements of the absolute number of mature rRNA molecules corresponding to the total BM CD45^+^ population, which were then averaged and used to calculate the absolute number of mature rRNA molecules for each primary cell type (Supplemental Figure S9F-H).

#### Quantification of the Pol I degron cell line

Quantification of mature rRNAs in Pol I degron cells was carried out as described above. In brief, Pol I degron cells were plated at 30K cells/mL 24 hours prior to dTAG treatment to allow for recovery, and cells were then treated with DMSO or 400 nM dTAG for 24 hours. 4 x 10^6^ cells were harvested and total RNA was extracted using the TRIzol-chloroform method without column purification. Two dilutions (1:1 and 1:4) for each RNA sample were prepared and analyzed using the Agilent 4150 TapeStation System. Mature rRNA abundance was averaged across the two dilutions for each sample, and quantifications were normalized to the average of the DMSO replicates to obtain the relative abundance of each mature rRNA species.

#### Quantification of primary bone marrow

Quantification of mature rRNAs in total bone marrow samples from wild type and AML (DF) mice was carried out as described above. In brief, total bone marrow cells were freshly isolated following mouse euthanasia, and samples of 10 million cells were harvested for total RNA extraction using the TRIzol-Chloroform method without column purification. One dilution (1:2) for each sample was prepared and analyzed using the Agilent 4150 TapeStation System.

#### Identification of p53 mutation in ER-HoxA9 Pol I degron cells

PCR amplification of the *Trp53* coding region was performed using genomic DNA from ER-HoxA9 Pol I degron cells, and products were subjected to Sanger sequencing. Sequence analysis identified a C-to-A substitution that introduces a premature stop codon in *Trp53* (Cys → Stop).

#### Analysis of core rDNA transcriptional machinery gene expression

Single-cell RNA-sequencing (scRNA-seq) datasets (H5 or RDS files) were downloaded from GEO (accessions GSE142341^31^ and GSE175702^32^). Seurat^76^ was used to read the datasets and extract average gene expression of components of the core rDNA transcriptional machinery and associated factors for each cell type. For visualization, gene expression values were scaled to a fixed range from –2 to 2 using z-score normalization across all cell types.

#### Analysis of rDNA methylation

Mouse whole Genome Bisulfite-Sequencing (WGBS) and Reduced Representation Bisulfite Sequencing (RRBS) datasets were downloaded from GEO (WGBS accession GSE52709 and RRBS accessions GSE77026; GSE57114). Human WGBS datasets were procured from the European Genome-Phenome Archive (accession EGAD00001002732). Raw reads were trimmed of adapter sequences using Trimmomatic v0.38^77^, and mapped using Bowtie2 v2.5.0^78^ to customized genome assemblies (Mouse_mm39-rDNA_genome_v1.0, Human_hg38-rDNA_genome_v1.0) that were previously generated by our lab by masking endogenous rDNA-like sequences and adding a single reference rDNA sequence of the appropriate species as a ‘Chromosome R’^37^. Non-bisulfite converted read filtering and methylation calling were performed using Bismark v0.24.2^79^. We exploited the dense read coverage over chrR to apply stringent mapping cutoffs (minimum of 300 reads for WGBS and 1500 reads for RRBS at each CpG), enabling high confidence quantification of methylation percentage at sites that passed the threshold. rDNA methylation shows binary distributions, with near-complete methylation across the IGS, and low methylation across the promoter and transcribed region. Methylation percentage of each rDNA region was calculated by taking the median of the methylation ratio ( methylated reads / [methylated+unmethylated reads]) across all CpGs within the region. Plotted methylation tracks (Figure 5D) were smoothed for visualization using LOESS regression (locally estimated scatterplot smoothing) = 0.015.

#### Analysis of rDNA accessibility

Human and mouse scATAC-seq datasets were downloaded from GEO (accessions GSE171220; GSE183675) and K562 scATAC-seq datasets were obtained from ENCODE (ENCSR217VXJ, ENCSR332SXG, ENCSR308ZGJ). Raw scATACseq reads were mapped to custom genome assemblies (Mouse_mm39-rDNA_genome_v1.0, Human_hg38-rDNA_genome_v1.0) previously generated by our lab^37^ using cellranger-atac^80^ (10X Genomics). ArchR v1.0.2^81^ was used to assign cell type identities to scATAC-seq barcodes from scRNA-seq datasets of the matching species in order to harmonize our custom scATAC-seq analysis with the above-described scRNA-seq analysis and with cell populations included in our study^82^. ArchR was further used to call peaks and perform transcription factor motif analysis. Plotted accessibility tracks (Figure 6B-C; Supplemental Figure S8A-B) were smoothed for visualization using LOESS regression (locally estimated scatterplot smoothing) = 0.006.

To identify genome-wide accessibility programs associated with differential rDNA accessibility, cells were stratified into populations with the top and bottom 10% rDNA accessibility (Thongon et al. human CD34^+^ dataset^38^, comprising a combined population of HSC, LMPP, Cycling progenitor, and MPP-MkEry) or top and bottom 20% rDNA accessibility (ENCODE K562 cell line dataset). Percent cutoffs were defined empirically for each dataset by testing a range of cutoffs from 5% to 30%, and picking the cutoff showing the greatest number of differential peaks in top vs bottom analysis. ArchR v1.0.2^81^ was used to identify TF motifs enriched in high rDNA accessibility cells compared to low in each dataset. The requirement of each enriched TF for global vs AML-specific cell growth was assessed by integrating CRISPR-based gene dependency scores from the Cancer Dependency Map (DepMap 24Q4^44^) (https://depmap.org/portal/).

Bulk ATAC-seq datasets were downloaded from GEO (accessions GSE146616; GSE256495; GSE205989), and raw reads were trimmed of adapter sequences using Trimmomatic v0.38^77^ and mapped to customized genome assemblies^37^ (Mouse_mm39-rDNA_genome_v1.0, Human_hg38-rDNA_genome_v1.0) using Bowtie2 v2.5.0^78^). BigWig tracks were generated using bamCoverage from deepTools v3.5.1^83^. Signal coverage across each rDNA region was normalized by the region length and reads per million. Coverage was further normalized to the median signal in the rDNA intergenic spacer region (IGS) across all samples to generate fold-change values relative to background, which were used to define relative accessibility.

#### Data visualization and statistical analysis

All bar graphs were created in GraphPad Prism v10.4.2. Scatterplots, heatmaps, and rDNA tracks were created using ggplot2 v3.5.2^84^ in RStudio. Representative flow cytometry plots were generated from FlowJo version 10.10.0 (BD Biosciences). Figures and schematics were generated using Adobe Illustrator and BioRender.com. Statistical analysis was completed using GraphPad Prism v10.4.2 and R v4.3.2^85^. Statistical significance is indicated as follows: ns (*p* ≥ 0.05), * (*p* < 0.05), ** (*p* < 0.01), *** (*p* < 0.001), **** (*p* < 0.0001).

## Data availability

Processed scATAC-seq datasets (ArchR objects and cell-type-specific bigwig tracks) will be deposited to GEO. Custom genomes used for rDNA mapping are available from GitHub (https://github.com/vikramparalkar/rDNA-Mapping-Genomes) as previously published^37^.

| **Antibody** | **Species** | **Clone** | **Fluorophore** | **Vendor/Catalog Number** |
| --- | --- | --- | --- | --- |
| CD106 | Rat anti-mouse | 429 (MVCAM.A) | BV650 | BD Biosciences, 740471 |
| CD115 | Rat anti-mouse | AFS98 | APC-Cy7 | BioLegend, 135531 |
| CD117 (c-Kit) | Rat anti-mouse | 2B8 | BV605 | BioLegend, 105847 |
| CD11b | Rat anti-mouse | M1/70 | FITC | BioLegend, 101205 |
| CD11c | Hamster anti-mouse | N418 | Biotin | BioLegend, 117303 |
| CD150 (SLAM) | Rat anti-mouse | TC15-12F12.2 | PE-Cy7 | BioLegend, 115913 |
| CD16/32 | Rat anti-mouse | S17011E | AF700 | BioLegend, 156619 |
| CD16/32 | Rat anti-mouse | 93 | PE-Cy7 | BioLegend, 101317 |
| CD19 | Rat anti-mouse | 6D5 | PerCP-Cy5.5 | BioLegend, 115533 |
| CD19 | Rat anti-mouse | 6D5 | Biotin | BioLegend, 115503 |
| CD3 | Rat anti-mouse | 17A2 | BV711 | BioLegend, 100241 |
| CD3 | Rat anti-mouse | 17A2 | Biotin | BioLegend, 100243 |
| CD34 | Rat anti-mouse | RAM34 | FITC | Thermo Fisher, 11-0341-82 |
| CD44 | Rat anti-mouse | IM7 | BV785 | BioLegend, 103041 |
| CD45.1 | Mouse anti-mouse | A20 | BV785 | BioLegend, 110743 |
| CD45.1 | Mouse anti-mouse | A20 | PE | BioLegend, 110707 |
| CD45.2 | Mouse anti-mouse | 104 | AF700 | Southern Biotechnology, 1800-27 |
| CD45.2 | Mouse anti-mouse | 104 | PE | BioLegend, 109807 |
| CD45R (B220) | Rat anti-mouse | RA3-6B2 | PE-Cy7 | BioLegend, 103221 |
| CD45R (B220) | Rat anti-mouse | RA3-6B2 | Biotin | BioLegend, 103203 |
| CD48 | Hamster anti-mouse | HM48-1 | BV711 | BioLegend, 103439 |
| CD49b | Rat anti-mouse | DX5 | Biotin | BioLegend, 108903 |
| CD81 | Hamster anti-mouse | 2F7 | PE | Southern Biotechnology, 1825-09 |
| CD90.2 | Rat anti-mouse | 53-2.1 | Biotin | BioLegend, 140313 |
| IgM | Mouse anti-mouse | MHM-88 | Pacific Blue | BioLegend, 314513 |
| Ly6A/E (Sca-1) | Rat anti-mouse | D7 | BV650 | BioLegend, 108143 |
| Ly6A/E (Sca-1) | Rat anti-mouse | D7 | Biotin | BioLegend, 108103 |
| Ly6C | Rat anti-mouse | HK1.4 | BV785 | BioLegend, 128041 |
| Ly6G | Rat anti-mouse | 1A8 | BV605 | BioLegend, 127633 |
| Ly6G | Rat anti-mouse | 1A8 | Biotin | BioLegend, 127603 |
| Ly6G/Ly6C (Gr-1) | Rat anti-mouse | RB6-8C5 | Biotin | BioLegend, 108403 |
| NK1.1 | Mouse anti-mouse | PK136 | Biotin | BioLegend, 108703 |
| Siglec F | Rat anti-mouse | S17007L | PE-Cy7 | BioLegend, 155527 |
| Streptavidin | Anti-all species | N/A | PerCP-Cy5.5 | BioLegend, 405214 |
| Ter119 | Rat anti-mouse | TER-119 | BV650 | BioLegend, 116235 |
| Ter119 | Rat anti-mouse | TER-119 | Biotin | BioLegend, 116203 |

## Acknowledgements

We would like to thank Nancy Speck, Gerd Blobel, Peter Klein, Martin Carroll, Ivan Maillard, Rahul Kohli, Crystal Conn, John Murray, and members of the Paralkar Lab for valuable feedback and scientific discussion. V.R.P. is supported by National Institutes of Health (NIH) grants R35 GM138035 (National Institute of General Medical Sciences [NIGMS]) and R21 HL175623 (National Heart, Lung, and Blood Institute [NHLBI]), the Greg Wolf Fund, the University of Pennsylvania University Research Foundation [URF] Grants Program, and philanthropic funding from Meghneel Gore and Anjali Dave. E.I.S, J.A.H., Z.W., and S.L.M. are Cell and Molecular Biology students through the Biomedical Graduate Studies program at the University of Pennsylvania. E.I.S is supported by the PERFORM-KUH U2C/TL1 grant DK136784 (National Institute of Diabetes and Digestive and Kidney Diseases [NIDDK]). C.A. is supported by a Fellow Scholar Award from the American Society of Hematology (ASH) and a Co-Operative Center for Excellence in Hematology (CCEH) Type B grant by the National Institute of Diabetes and Digestive and Kidney Diseases (NIDDK). K.F.L. is supported by NIH grants R35GM156342 (NIGMS) and R01HL160726 (NHLBI). J.A.H. is supported by NIH grant T32 HL007439 (National Heart, Lung, and Blood Institute [NHLBI]). P.J.H. is supported by NIH grant T32 HL007439-45 (National Heart, Lung, and Blood Institute [NHLBI]), the Marlene Shlomchik Fellowship in Cancer Research (Abramson Cancer Center, University of Pennsylvania), and the AACR-John and Elizabeth Leonard Family Foundation Basic Cancer Research Fellowship (American Association for Cancer Research [AACR]). N.E. is supported by NIH grant F30 CA281040, NIH training grants T32 GM008216 and T32 GM007170, and a Mike S. Brown Dissertation Fellowship. Research on this project was supported by the University of Pennsylvania Abramson Cancer Center Support Grant, supported by National Cancer Institute (NCI) grant P30 CA016520. Flow cytometry data were generated in the University of Pennsylvania Cytomics and Cell Sorting Shared Resource Laboratory at the University of Pennsylvania (RRID: SCR_022376). Microscopy data were generated in the Cell and Developmental Biology Microscopy Core at the University of Pennsylvania (RRID: SCR_022373).

## Author contributions

V.R.P. conceived the project and supervised the study. E.I.S. and V.K.F. designed and performed all primary bone marrow FISH-Flow experiments, with rRNA FISH-Flow protocol optimization by E.M.G. and additional studies by C.A. The Pol I degron cell line was designed and generated by C.A. All Pol I degron experiments were designed and performed by E.I.S., with contributions from S.L.M. for 5-EU analyses, contributions from Y.G. and K.F.L. for Northern blot analyses, and contributions from D.G. and W.T. for sucrose gradient fractionation analyses. Bioinformatic analyses for bisulfite sequencing and ATAC-seq were performed by S.S.G. and V.R.P., with additional input from N.E. rRNA FISH-Flow probes were generated by C.A., E.I.S, and K.A., with contributions from J.W., M.C.D., J.C.J.H., G.M.B., and A.R. V.K.F., J.A.H. K.A., P.J.H., and Z.W. contributed to breeding and maintaining the mouse colony and harvesting mouse bone marrow. Primary cells for *Npm1^c^*;*Flt3*^ITD^ AML transplant experiments were provided by N.S. and R.L.B. R.A.J.S. provided protein synthesis data. E.I.S. and V.K.F. made and edited the figures, including writing scripts for graphical representation. E.I.S. and V.R.P. wrote the manuscript with input from all authors.

## REFERENCES

1. Paralkar, V. R. Transcription factor regulation of ribosomal RNA in hematopoiesis. Current Opinion in Hematology 31, 199 (2024).

2. Daiß, J. L., Griesenbeck, J., Tschochner, H. & Engel, C. Synthesis of the ribosomal RNA precursor in human cells: mechanisms, factors and regulation. Biological chemistry 404, (2023).

3. Moss, T., Langlois, F., Gagnon-Kugler, T. & Stefanovsky, V. A housekeeper with power of attorney: the rRNA genes in ribosome biogenesis. Cellular and Molecular Life Sciences 64, 29–49 (2006).

4. Islam, R. A. & Rallis, C. Ribosomal Biogenesis and Heterogeneity in Development, Disease, and Aging. Epigenomes 7, (2023).

5. Pederson, T. The nucleolus. Cold Spring Harb. Perspect. Biol. 3, a000638–a000638 (2011).

6. Sharifi, S. & Bierhoff, H. Regulation of RNA Polymerase I Transcription in Development, Disease, and Aging. Annual review of biochemistry 87, (2018).

7. Moss, T., Mars, J. C., Tremblay, M. G. & Sabourin-Felix, M. The chromatin landscape of the ribosomal RNA genes in mouse and human. Chromosome research: an international journal on the molecular, supramolecular and evolutionary aspects of chromosome biology 27, (2019).

8. Quinodoz, S. A. et al. Mapping and engineering RNA-driven architecture of the multiphase nucleolus. Nature (2025) doi:10.1038/s41586-025-09207-4.

9. Dörner, K., Ruggeri, C., Zemp, I. & Kutay, U. Ribosome biogenesis factors—from names to functions. The EMBO Journal (2023) doi:10.15252/embj.2022112699.

10. Baßler, J. & Hurt, E. Eukaryotic Ribosome Assembly. Annual review of biochemistry 88, (2019).

11. Olson, O. C., Kang, Y. A. & Passegué, E. Normal Hematopoiesis Is a Balancing Act of Self-Renewal and Regeneration. Cold Spring Harbor perspectives in medicine 10, (2020).

12. Rieger, M. A. & Schroeder, T. Hematopoiesis. Cold Spring Harb Perspect Biol 4, a008250 (2012).

13. Cedar, H. & Bergman, Y. Epigenetics of haematopoietic cell development. Nature Reviews Immunology 11, 478–488 (2011).

14. L. A. Liggett, V. G. S. Unraveling Hematopoiesis through the Lens of Genomics. Cell 182, 1384–1400 (2020).

15. Signer, R. A. J. et al. The rate of protein synthesis in hematopoietic stem cells is limited partly by 4E-BPs. Genes Dev 30, 1698–1703 (2016).

16. Signer, R. A. J., Magee, J. A., Salic, A. & Morrison, S. J. Haematopoietic stem cells require a highly regulated protein synthesis rate. Nature 509, 49–54 (2014).

17. Antony, C. et al. Control of ribosomal RNA synthesis by hematopoietic transcription factors. Mol Cell 82, 3826–3839.e9 (2022).

18. Le Goff, S. et al. p53 activation during ribosome biogenesis regulates normal erythroid differentiation. Blood 137, (2021).

19. Jarzebowski, L. et al. Mouse adult hematopoietic stem cells actively synthesize ribosomal RNA. RNA 24, 1803–1812 (2018).

20. Hayashi, Y. et al. Downregulation of rRNA transcription triggers cell differentiation. PLoS One 9, e98586 (2014).

21. Smetana, K., Šubrtová, H., Jirásková, I. & Rosa, L. A Further Study on the Incidence of Nucleoli in Myeloblasts of Patients Suffering from Acute Myeloid Leukemia. Hematology (Amsterdam, Netherlands) 2, (1997).

22. Hein, N., Hannan, K. M., George, A. J., Sanij, E. & Hannan, R. D. The nucleolus: an emerging target for cancer therapy. Trends in molecular medicine 19, (2013).

23. Antony, C., Somers, P., Gray, E. M., Pimkin, M. & Paralkar, V. R. FISH-Flow to quantify nascent and mature ribosomal RNA in mouse and human cells. STAR Protoc 4, 102463 (2023).

24. Valnes, K. & Brandtzaeg, P. Selective inhibition of nonspecific eosinophil staining or identification of eosinophilic granulocytes by paired counterstaining in immunofluorescence studies. J Histochem Cytochem 29, 595–400 (1981).

25. Papaemmanuil, E. et al. Genomic Classification and Prognosis in Acute Myeloid Leukemia. N Engl J Med 374, 2209–2221 (2016).

26. Meyer, S. E. et al. DNMT3A Haploinsufficiency Transforms FLT3ITD Myeloproliferative Disease into a Rapid, Spontaneous, and Fully Penetrant Acute Myeloid Leukemia. Cancer Discov 6, 501–515 (2016).

27. Bowman, R. L. et al. In vivo models of subclonal oncogenesis and dependency in hematopoietic malignancy. Cancer Cell 42, 1955–1969.e7 (2024).

28. Singh, I. et al. Leukemia stem cell expansion cultures reveal clonal drivers of leukemogenesis and therapy response. bioRxiv (2026) doi:10.64898/2026.02.24.707683.

29. Muhar, M. et al. SLAM-seq defines direct gene-regulatory functions of the BRD4-MYC axis. Science (New York, N.Y.) 360, (2018).

30. Poortinga, G. et al. c-MYC coordinately regulates ribosomal gene chromatin remodeling and Pol I availability during granulocyte differentiation. Nucleic acids research 39, (2011).

31. Muench, D. E. et al. Mouse models of neutropenia reveal progenitor-stage-specific defects. Nature 582, 109–114 (2020).

32. Konturek-Ciesla, A. et al. Temporal multimodal single-cell profiling of native hematopoiesis illuminates altered differentiation trajectories with age. Cell Rep 42, 112304 (2023).

33. Hori, Y., Engel, C. & Kobayashi, T. Regulation of ribosomal RNA gene copy number, transcription and nucleolus organization in eukaryotes. Nature Reviews Molecular Cell Biology 24, 414–429 (2023).

34. Hori, Y., Shimamoto, A. & Kobayashi, T. The human ribosomal DNA array is composed of highly homogenized tandem clusters. Genome research 31, (2021).

35. Farlik, M. et al. DNA Methylation Dynamics of Human Hematopoietic Stem Cell Differentiation. Cell Stem Cell 19, 808–822 (2016).

36. Cabezas-Wallscheid, N. et al. Identification of regulatory networks in HSCs and their immediate progeny via integrated proteome, transcriptome, and DNA methylome analysis. Cell Stem Cell 15, 507–522 (2014).

37. George, S. S., Pimkin, M. & Paralkar, V. R. Construction and validation of customized genomes for human and mouse ribosomal DNA mapping. J Biol Chem 299, 104766 (2023).

38. Thongon, N. et al. Hematopoiesis under telomere attrition at the single-cell resolution. Nat Commun 12, 6850 (2021).

39. Hong, T. et al. TET2 modulates spatial relocalization of heterochromatin in aged hematopoietic stem and progenitor cells. Nat Aging 3, 1387–1400 (2023).

40. Yun, H. et al. Mutational synergy during leukemia induction remodels chromatin accessibility, histone modifications and three-dimensional DNA topology to alter gene expression. Nat Genet 53, 1443–1455 (2021).

41. Nuno, K. et al. Convergent epigenetic evolution drives relapse in acute myeloid leukemia. Elife 13, (2024).

42. Takao, S. et al. Epigenetic mechanisms controlling human leukemia stem cells and therapy resistance. Nat Commun 16, 3196 (2025).

43. ENCODE Project Consortium. An integrated encyclopedia of DNA elements in the human genome. Nature 489, 57–74 (2012).

44. Website. DepMap, Broad (2024). DepMap 24Q4 Public. Figshare+. Dataset. 10.25452/figshare.plus.27993248.v1.

45. Sykes, D. B. et al. Inhibition of Dihydroorotate Dehydrogenase Overcomes Differentiation Blockade in Acute Myeloid Leukemia. Cell 167, 171–186.e15 (2016).

46. Agilent Technologies. Nucleic Acids Sizes and Molecular Weights. https://www.agilent.com/files/Mobio/Nucleic%20Acids_Sizes_and_Molecular_Weights_2pgs.pdf.

47. Loeb, J. N., Howell, R. R. & Tomkins, G. M. Turnover of ribosomal RNA in rat liver. Science 149, 1093–1095 (1965).

48. Stoykova, A. S., Dudov, K. P., Dabeva, M. D. & Hadjiolov, A. A. Different rates of synthesis and turnover of ribosomal RNA in rat brain and liver. J Neurochem 41, 942–949 (1983).

49. Passegué, E., Wagers, A. J., Giuriato, S., Anderson, W. C. & Weissman, I. L. Global analysis of proliferation and cell cycle gene expression in the regulation of hematopoietic stem and progenitor cell fates. The Journal of Experimental Medicine 202, 1599 (2005).

50. Pietras, E. M., Warr, M. R. & Passegué, E. Cell cycle regulation in hematopoietic stem cells. The Journal of Cell Biology 195, 709 (2011).

51. An, H. & Harper, J. W. Ribosome Abundance Control Via the Ubiquitin-Proteasome System and Autophagy. J Mol Biol 432, 170–184 (2020).

52. Lafontaine, D. L. J. A ‘garbage can’ for ribosomes: how eukaryotes degrade their ribosomes. Trends Biochem Sci 35, 267–277 (2010).

53. Kraft, C., Deplazes, A., Sohrmann, M. & Peter, M. Mature ribosomes are selectively degraded upon starvation by an autophagy pathway requiring the Ubp3p/Bre5p ubiquitin protease. Nat Cell Biol 10, 602–610 (2008).

54. Mortensen, M. et al. The autophagy protein Atg7 is essential for hematopoietic stem cell maintenance. J Exp Med 208, 455–467 (2011).

55. Ho, T. T. et al. Autophagy maintains the metabolism and function of young and old stem cells. Nature 543, 205–210 (2017).

56. Kasbekar, M., Mitchell, C. A., Proven, M. A. & Passegué, E. Hematopoietic stem cells through the ages: A lifetime of adaptation to organismal demands. Cell Stem Cell 30, 1403–1420 (2023).

57. Willi, J. et al. Oxidative stress damages rRNA inside the ribosome and differentially affects the catalytic center. Nucleic Acids Research 46, 1945 (2018).

58. Shcherbik, N. & Pestov, D. G. The Impact of Oxidative Stress on Ribosomes: From Injury to Regulation. Cells 8, 1379 (2019).

59. Botello, J. F., et al. Ribosome Molecular Aging Shapes Translation Dynamics. bioRxiv (2026) doi:10.64898/2026.03.08.710403.

60. Cole, S. E., LaRiviere, F. J., Merrikh, C. N. & Moore, M. J. A convergence of rRNA and mRNA quality control pathways revealed by mechanistic analysis of nonfunctional rRNA decay. Mol Cell 34, 440–450 (2009).

61. Coria, A. R. et al. The integrated stress response regulates 18S nonfunctional rRNA decay in mammals. Mol Cell 85, 787–801.e8 (2025).

62. Huang, Z. et al. RIOK3 mediates the degradation of 40S ribosomes. Mol Cell 85, 802–814.e12 (2025).

63. Ford, P. W. et al. RNF10 and RIOK3 facilitate 40S ribosomal subunit degradation upon 60S biogenesis disruption or amino acid starvation. Cell Rep 44, 115371 (2025).

64. Mills, E. W. & Green, R. Ribosomopathies: There’s strength in numbers. Science 358, (2017).

65. Wilson, A. et al. c-Myc controls the balance between hematopoietic stem cell self-renewal and differentiation. Genes & development 18, (2004).

66. Bahr, C. et al. A Myc enhancer cluster regulates normal and leukaemic haematopoietic stem cell hierarchies. Nature 553, (2018).

67. Gilmour, J. et al. A crucial role for the ubiquitously expressed transcription factor Sp1 at early stages of hematopoietic specification. *Development (Cambridge*, England) 141, (2014).

68. Gilmour, J. et al. Robust hematopoietic specification requires the ubiquitous Sp1 and Sp3 transcription factors. Epigenetics & chromatin 12, (2019).

69. Chan, J. Y. et al. Targeted disruption of the ubiquitous CNC-bZIP transcription factor, Nrf-1, results in anemia and embryonic lethality in mice. EMBO J 17, 1779–1787 (1998).

70. Kim, H. M., Han, J. W. & Chan, J. Y. Nuclear Factor Erythroid-2 Like 1 (NFE2L1): Structure, function and regulation. Gene 584, 17–25 (2016).

71. Pelletier, J., Thomas, G. & Volarević, S. Ribosome biogenesis in cancer: new players and therapeutic avenues. Nature reviews. Cancer 18, (2018).

72. Ferreira, R., et al. The novel RNA polymerase I transcription inhibitor PMR-116 exploits a critical therapeutic vulnerability in a broad-spectrum of high MYC malignancies. bioRxiv (2025) doi:10.1101/2025.04.19.649466.

73. Pastore, S. et al. Mapping human pre-rRNA processing and modification at single nucleotide resolution using long read nanopore sequencing. Nat Commun (2026) doi:10.1038/s41467-026-71164-x.

74. Gonskikh, Y. et al. Noncatalytic regulation of 18 rRNA methyltransferase DIMT1 in acute myeloid leukemia. Genes Dev 37, 321–335 (2023).

75. Raj, A., van den Bogaard, P., Rifkin, S. A., van Oudenaarden, A. & Tyagi, S. Imaging individual mRNA molecules using multiple singly labeled probes. Nature Methods 5, 877–879 (2008).

76. Satija, R., Farrell, J. A., Gennert, D., Schier, A. F. & Regev, A. Spatial reconstruction of single-cell gene expression data. Nature biotechnology 33, (2015).

77. Bolger, A. M., Lohse, M. & Usadel, B. Trimmomatic: a flexible trimmer for Illumina sequence data. Bioinformatics 30, 2114–2120 (2014).

78. Langmead, B. & Salzberg, S. L. Fast gapped-read alignment with Bowtie 2. Nat Methods 9, 357–359 (2012).

79. Krueger, F. & Andrews, S. R. Bismark: a flexible aligner and methylation caller for Bisulfite-Seq applications. *Bioinformatics (Oxford*, England) 27, (2011).

80. Satpathy, A. T. et al. Massively parallel single-cell chromatin landscapes of human immune cell development and intratumoral T cell exhaustion. Nat Biotechnol 37, 925–936 (2019).

81. Granja, J. M. et al. ArchR is a scalable software package for integrative single-cell chromatin accessibility analysis. Nature Genetics 53, 403–411 (2021).

82. Zeng, A. G. X. et al. Single-cell Transcriptional Atlas of Human Hematopoiesis Reveals Genetic and Hierarchy-Based Determinants of Aberrant AML Differentiation. Blood Cancer Discov 6, 307–324 (2025).

83. Ramírez, F. et al. deepTools2: a next generation web server for deep-sequencing data analysis. Nucleic acids research 44, (2016).

84. Wickham, H. ggplot2: Elegant Graphics for Data Analysis. (Springer Science & Business Media, 2009).

85. R Core Team (2023). R: A Language and Environment for Statistical Computing. R Foundation for Statistical Computing, Vienna, Austria. <https://www.R-project.org/>

