## Supplemental Figures for "Dynamics of Ribosomal RNA Transcription and Abundance in Normal and Leukemic Hematopoiesis"

#### Supplemental Figure S1

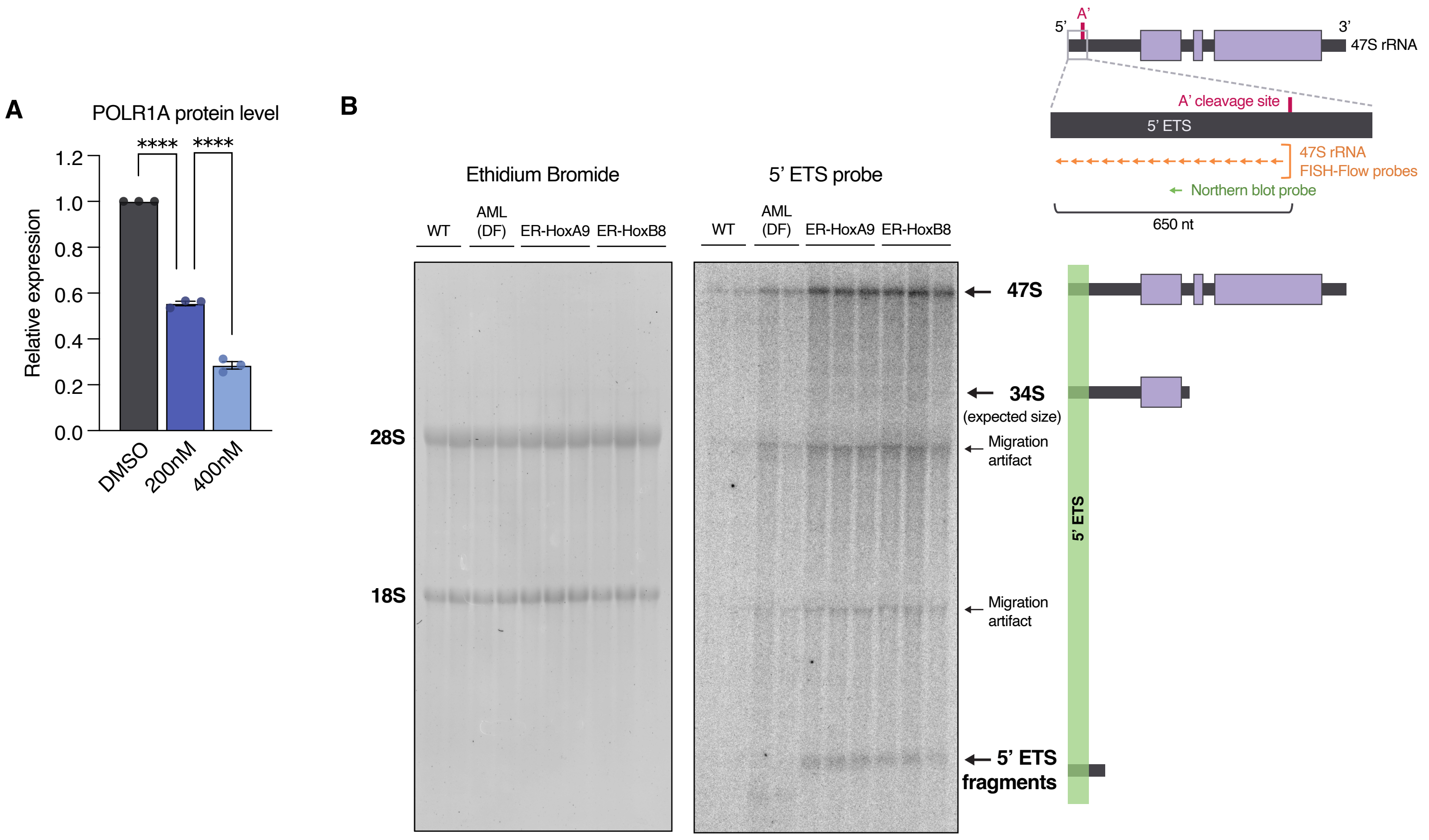


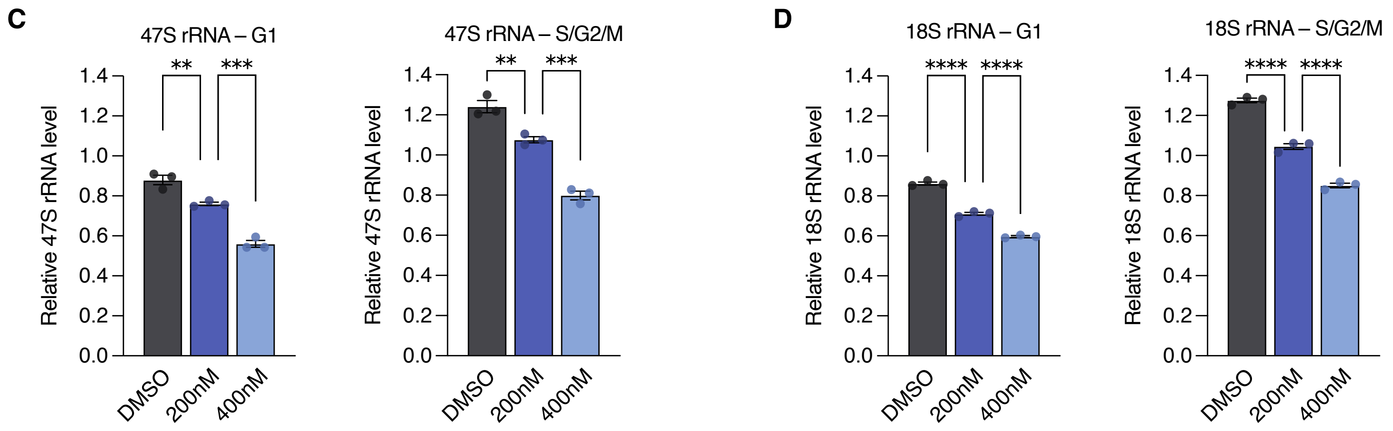


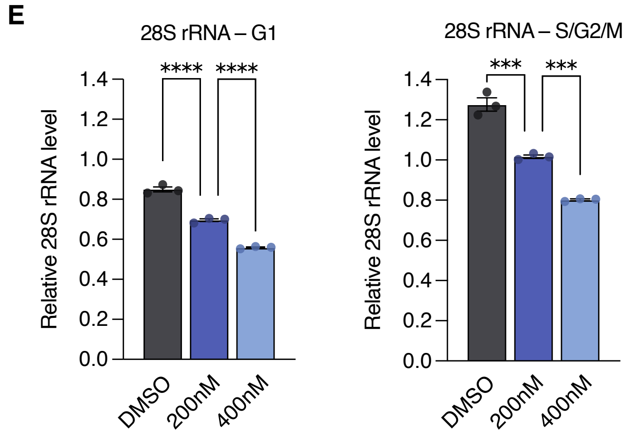

**Supplemental Figure S1. FISH-Flow measures dynamic changes in rRNA abundance in response to Pol I degradation (related to Figure 1)**

**(A)** Relative Pol I abundance measured by Western blot in Pol I degron cells treated with the indicated doses of dTAG for 24 hours, normalized to paired DMSO replicates. n = 3 replicates per condition.

**(B)** Left panel: Northern blot ethidium bromide staining showing loading of total RNA samples. Right panel: RNA molecules detected with a 5' ETS probe overlapping the region containing 47S rRNA FISH-Flow probes in ER-HoxA9 and ER-HoxB8 cell lines and in total wild type and AML (DF) bone marrow cells. The majority of 5' ETS probe signal is detected as full length 47S transcripts, with minimal contribution from small 5' ETS fragments. No aberrant 34S species is detected at the expected size. Migration artifacts are observed surrounding 18S and 28S rRNAs. n = 2-3 replicates per condition.

**(C)** Relative median abundance of nascent 47S rRNA in cells in G1 (left) and S/G2/M (right) cell cycle phases measured by FISH-Flow in Pol I degron cells treated with DMSO or dTAG for 24 hours, normalized to the average of total cells from DMSO replicates. n = 3 replicates per condition.

**(D)** Relative median abundance of mature 18S rRNA in cells in G1 (left) and S/G2/M (right) cell cycle phases measured by FISH-Flow in Pol I degron cells treated with DMSO or dTAG for 24 hours, normalized to the average of total cells from DMSO replicates. n = 3 replicates per condition.

**(E)** Relative median abundance of mature 28S rRNA in cells in G1 (left) and S/G2/M (right) cell cycle phases measured by FISH-Flow in Pol I degron cells treated with DMSO or dTAG for 24 hours, normalized to the average of total cells from DMSO replicates. n = 3 replicates per condition.

All bar graphs show mean ± SEM. ns ($p \geq 0.05$), * ($p < 0.05$), ** ($p < 0.01$), *** ($p < 0.001$), **** ($p < 0.0001$) by one-way ANOVA with Sidak's multiple comparison testing.

#### Supplemental Figure S2

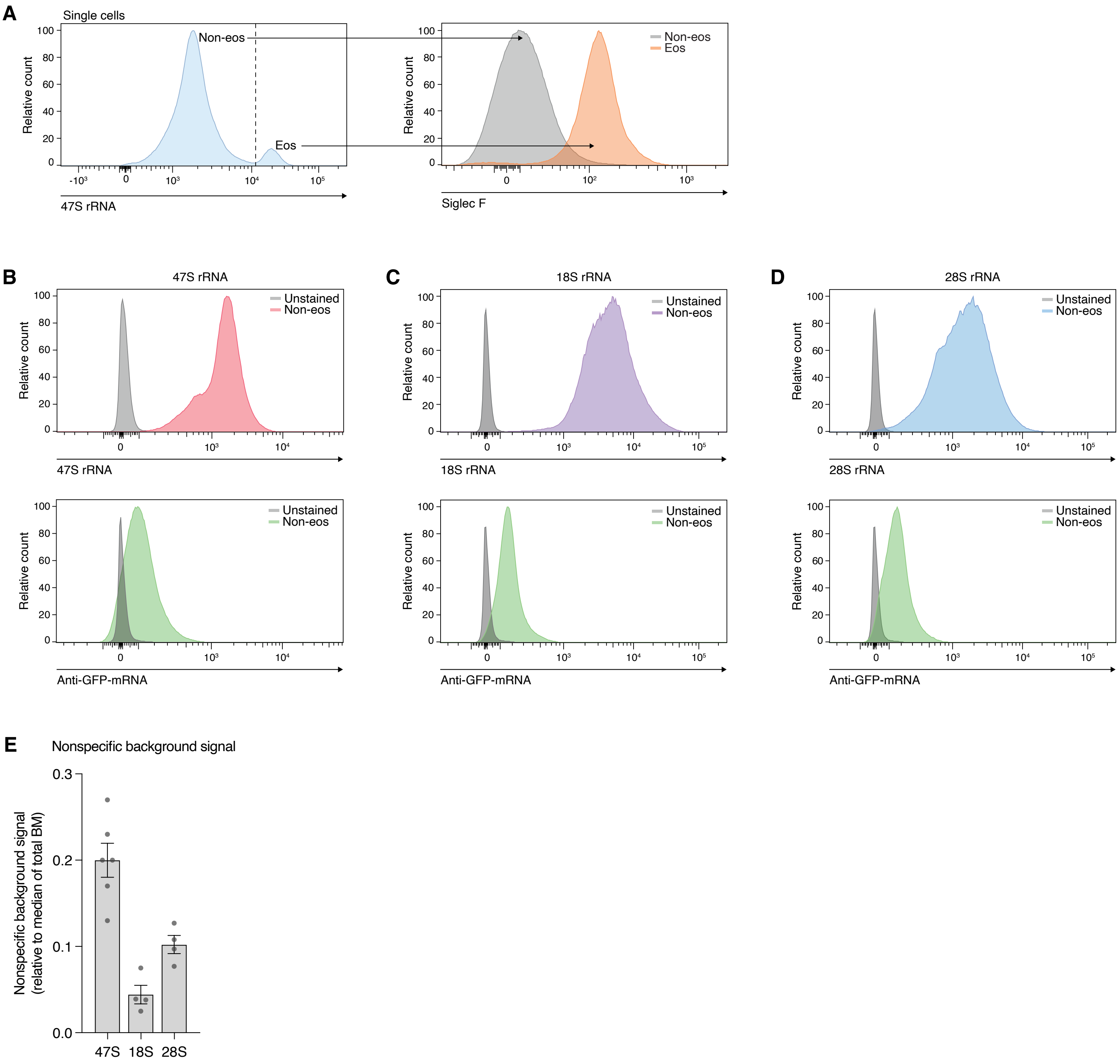

**Supplemental Figure S2. Defining threshold of nonspecific background FISH-Flow signal (related to Figure 2)**

**(A)** (Left) Representative flow cytometry plot of 47S rRNA probe signal in total single cells. Most cells fall within a singular distribution, with the exception of a small population with aberrantly high signal. (Right) The high probe signal population selectively expresses eosinophil (eos) marker Siglec F, and is excluded for all further analyses.

**(B-D)** Representative flow cytometry plot of rRNA FISH-Flow probe signal (top) and negative control anti-GFP-mRNA probe signal (bottom) in all non-eos (cells co-stained with both probe sets) versus unstained cells. Comparisons for 47S (B), 18S (C), and 28S (D) rRNA are shown.

**(E)** Average nonspecific FISH-Flow probe background signal relative to total BM signal for each rRNA probe set. All bars show mean ± SEM.

### Supplemental Figure S3

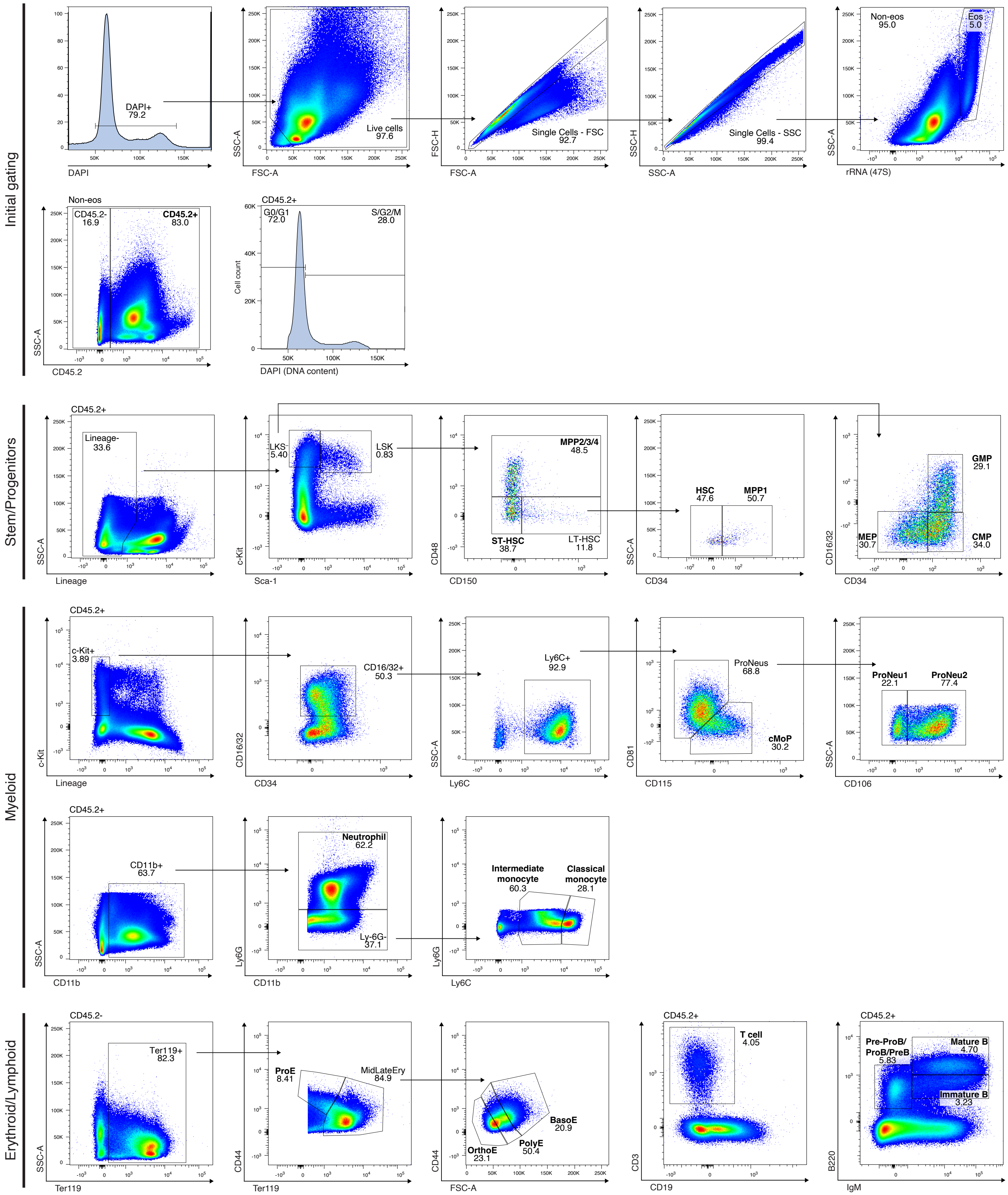

**Supplemental Figure S3. Normal hematopoiesis gating strategies (related to Figure 2)**

Representative flow cytometry plots depicting the gating strategies used for all normal hematopoietic cell types included in this study. Gates listed in the "Initial gating" section were applied to all samples. Final populations used for analysis and plotting are written in bold text. Numbers underneath population names indicate the percentage contribution to the represented plot. Full written gating schemes are listed in Supplemental Table S1.

**Supplemental Figure S4**

**A**

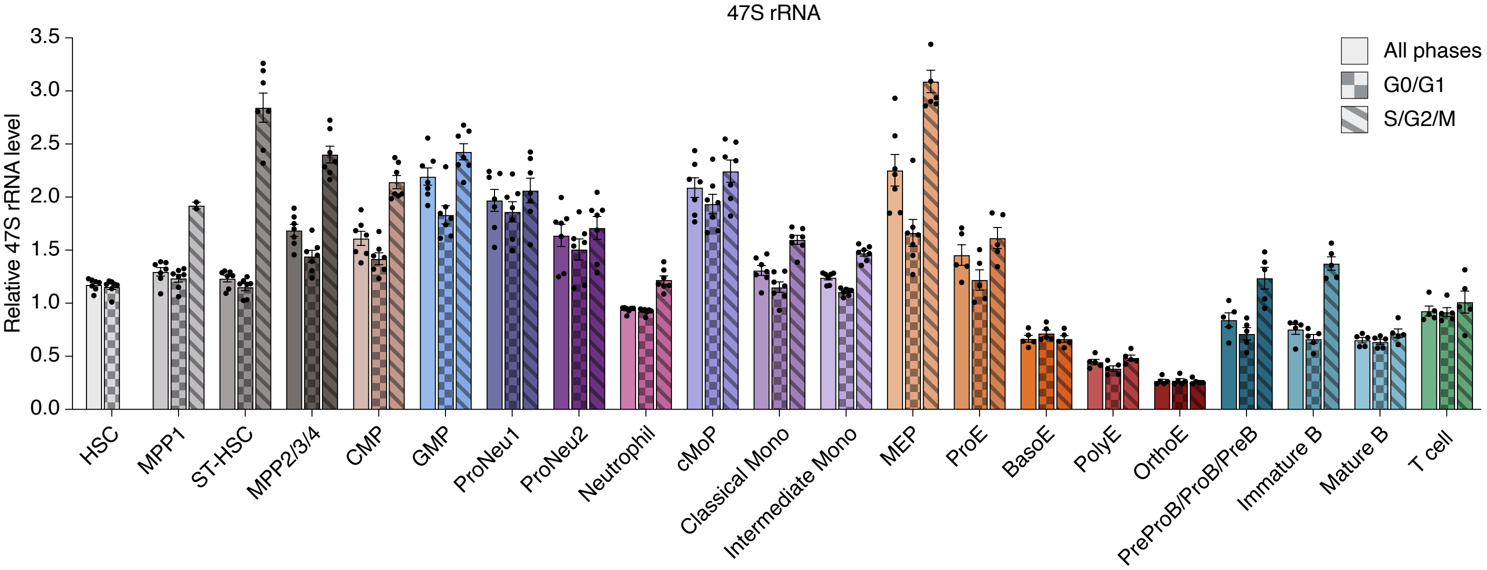


**B**

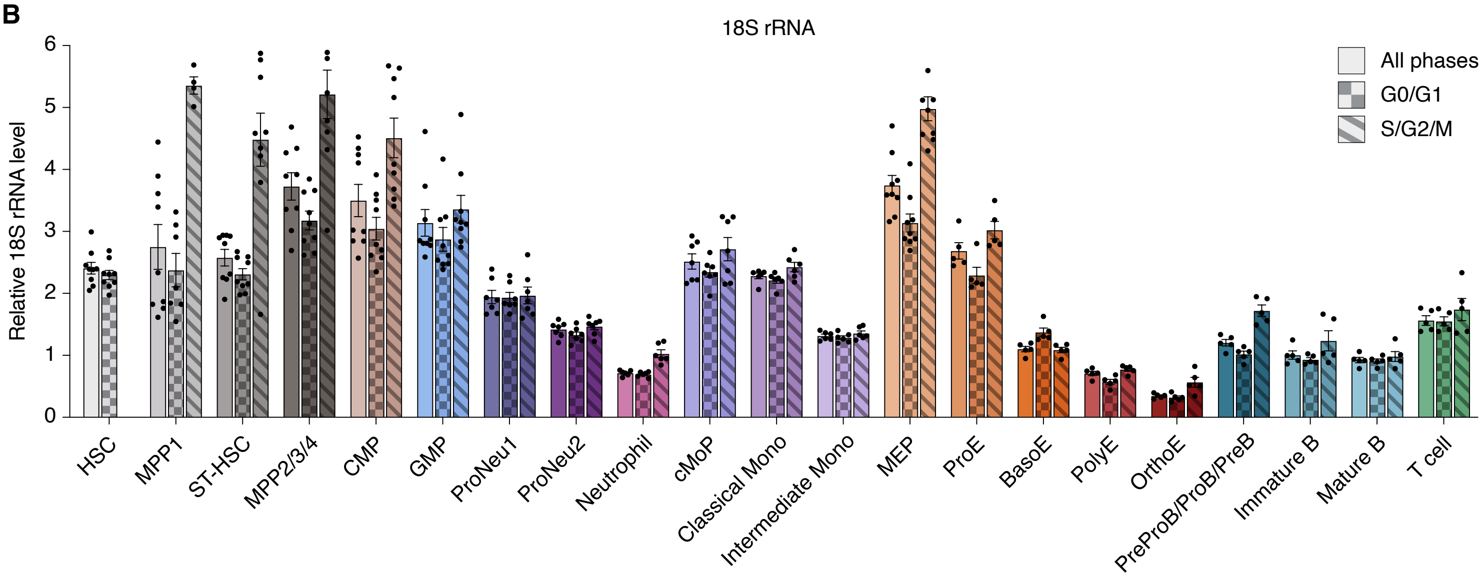


**C**

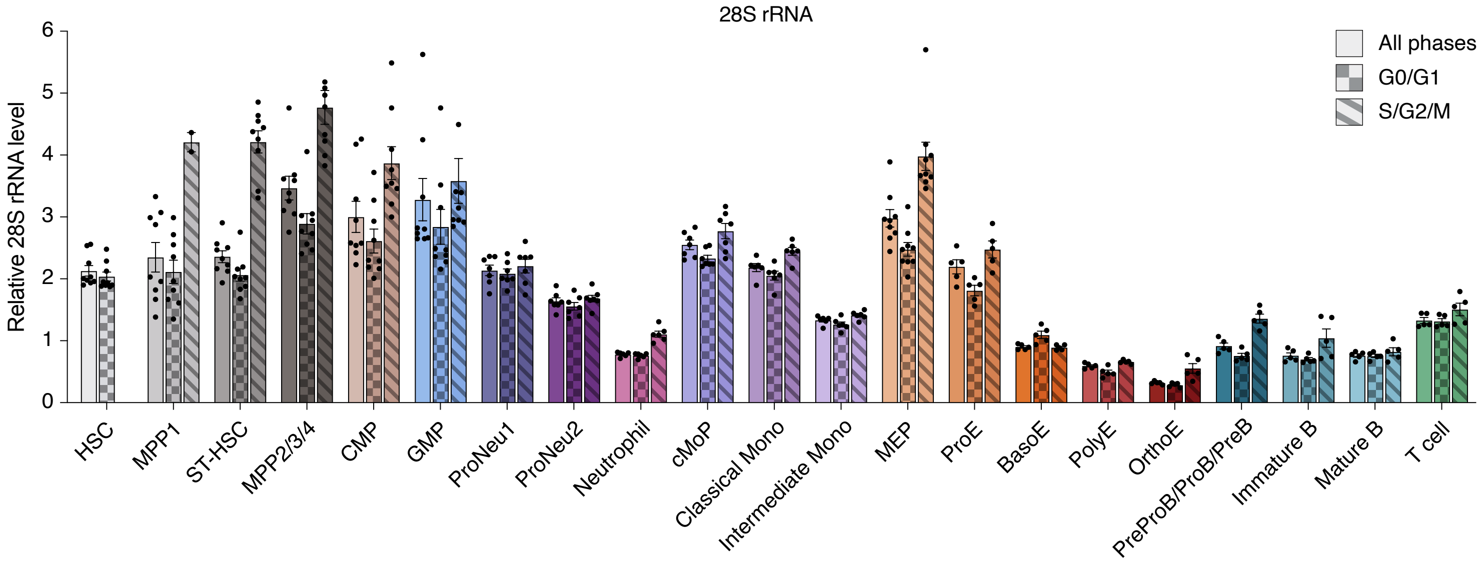

**Supplemental Figure S4. rRNA levels across cell cycle phases in normal hematopoietic cell types (related to Figure 2)**

**(A-C)** Relative abundance of 47S rRNA (A), 18S rRNA (B), and 28S rRNA (C) in cell cycle phase subpopulations of each normal hematopoietic cell type. The total cell population is represented by a solid bar, the cell population in G0/G1 phases is represented by a checkered bar, and the cell population in S/G2/M phases is represented by a striped bar. All values are normalized to the median level of each respective rRNA in the total BM $CD45^+$ population (total cells). A minimal number of HSCs are in S/G2/M, and are not shown.

In some cases, replicates outside of the axis limits are not shown ( ≤ 2 replicates per cell type). All bar graphs and scatter plots show mean ± SEM.

### Supplemental Figure S5

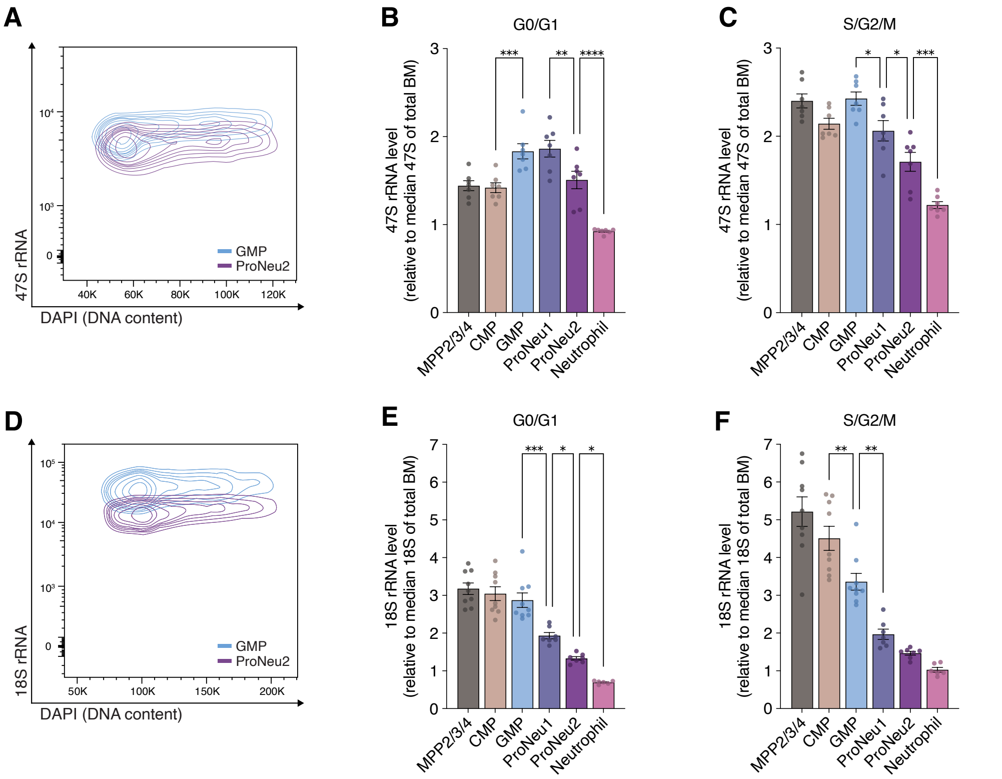


G

| Cell type | % S/G2/M |
|---|---|
| HSC | 11.7 |
| MPP1 | 18.9 |
| ST-HSC | 20.2 |
| MPP2/3/4 | 34.3 |
| CMP | 33.3 |
| GMP | 63.6 |
| ProNeu1 | 63.3 |
| ProNeu2 | 70.2 |
| Neutrophil | 12.2 |
| cMoP | 57.8 |
| Classical Monocyte | 44.8 |
| Intermediate Monocyte | 45.0 |
| MEP | 43.4 |
| ProE | 63.2 |
| BasoE | 95.5 |
| PolyE | 61.5 |
| OrthoE | 25.3 |
| PreProB/ProB/PreB | 34.6 |
| Immature B | 32.2 |
| Mature B | 39.2 |
| T cell | 14.2 |

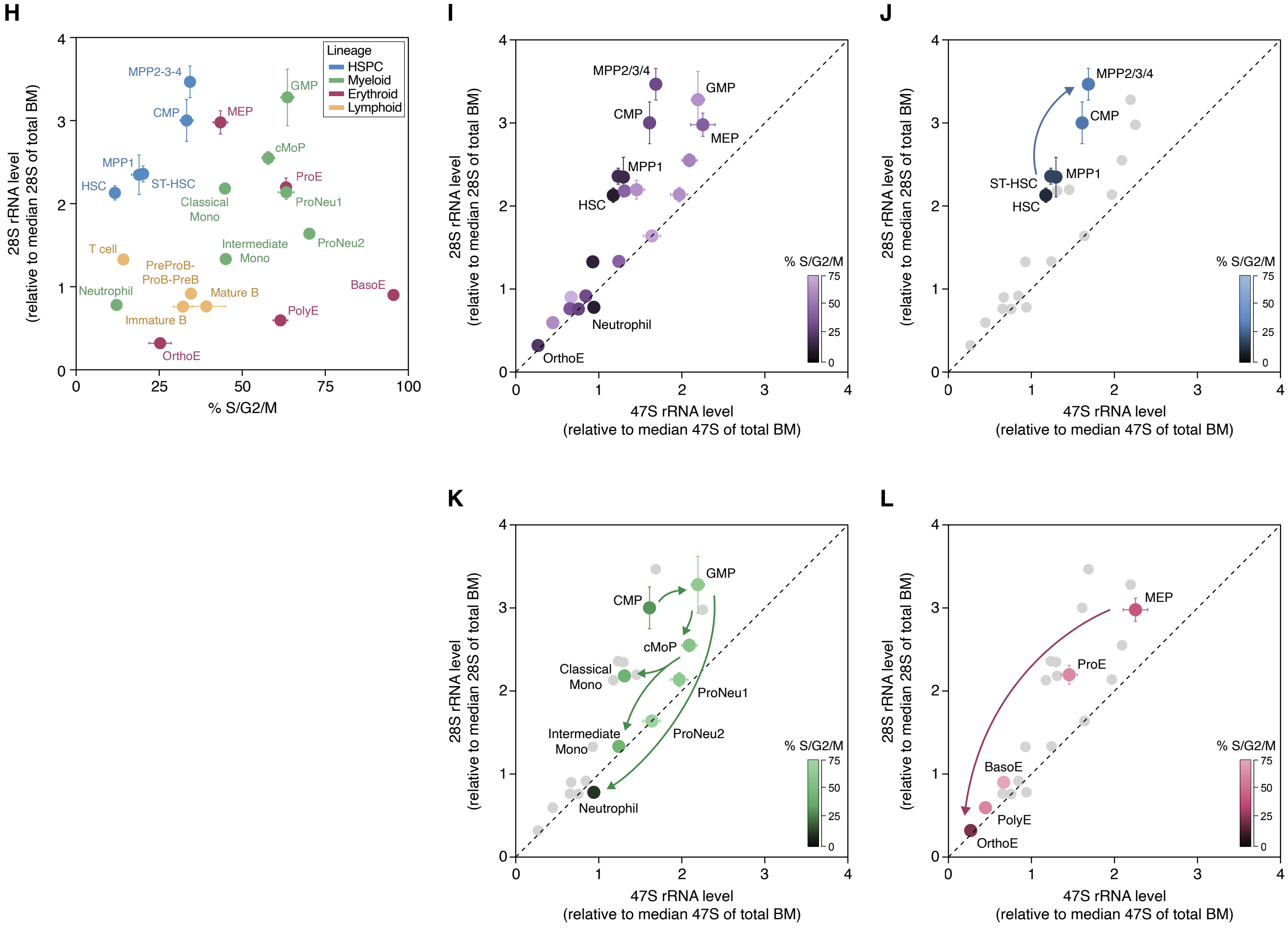

**Supplemental Figure S5. Trends in nascent and mature rRNAs and cell cycling in normal hematopoiesis (related to Figures 2 and 3)**

**(A)** Representative flow cytometry plot comparing 47S rRNA abundance across cell cycle phases (DAPI) in GMPs (top, blue) and ProNeu2 (bottom, purple) populations.

**(B-C)** Relative abundance of 47S rRNA (normalized to the median 47S level of total BM) across the myeloid lineage in cells in G0/G1 (B) and S/G2/M (C) phases of the cell cycle.

**(D-F)** Same as in A-C, showing 18S rRNA.

**(G)** Table of the average percentage of cells in S/G2/M cell cycle phases for all normal hematopoiesis cell types.

**(H)** Comparison of cell cycle phase distribution (% S/G2/M) and 28S rRNA level across normal hematopoietic lineages.

**(I-L)** Comparison of relative rRNA transcription level (47S rRNA) and ribosome subunit abundance (28S rRNA) across all normal hematopoietic cell types (I), HSPCs (J), myeloid lineage (K), and erythroid lineage (L). The color of each cell type represents the percentage of cells in S/G2/M cell cycle phases. Color scales are capped at 75%.

In some cases, replicates outside of the axis limits are not shown ( ≤ 2 replicates per cell type). All bar graphs and scatter plots show mean ± SEM. ns ($p \geq 0.05$), * ($p < 0.05$), ** ($p < 0.01$), *** ($p < 0.001$), **** ($p < 0.0001$) by one-way ANOVA with Sidak's multiple comparison testing.

#### Supplemental Figure S6

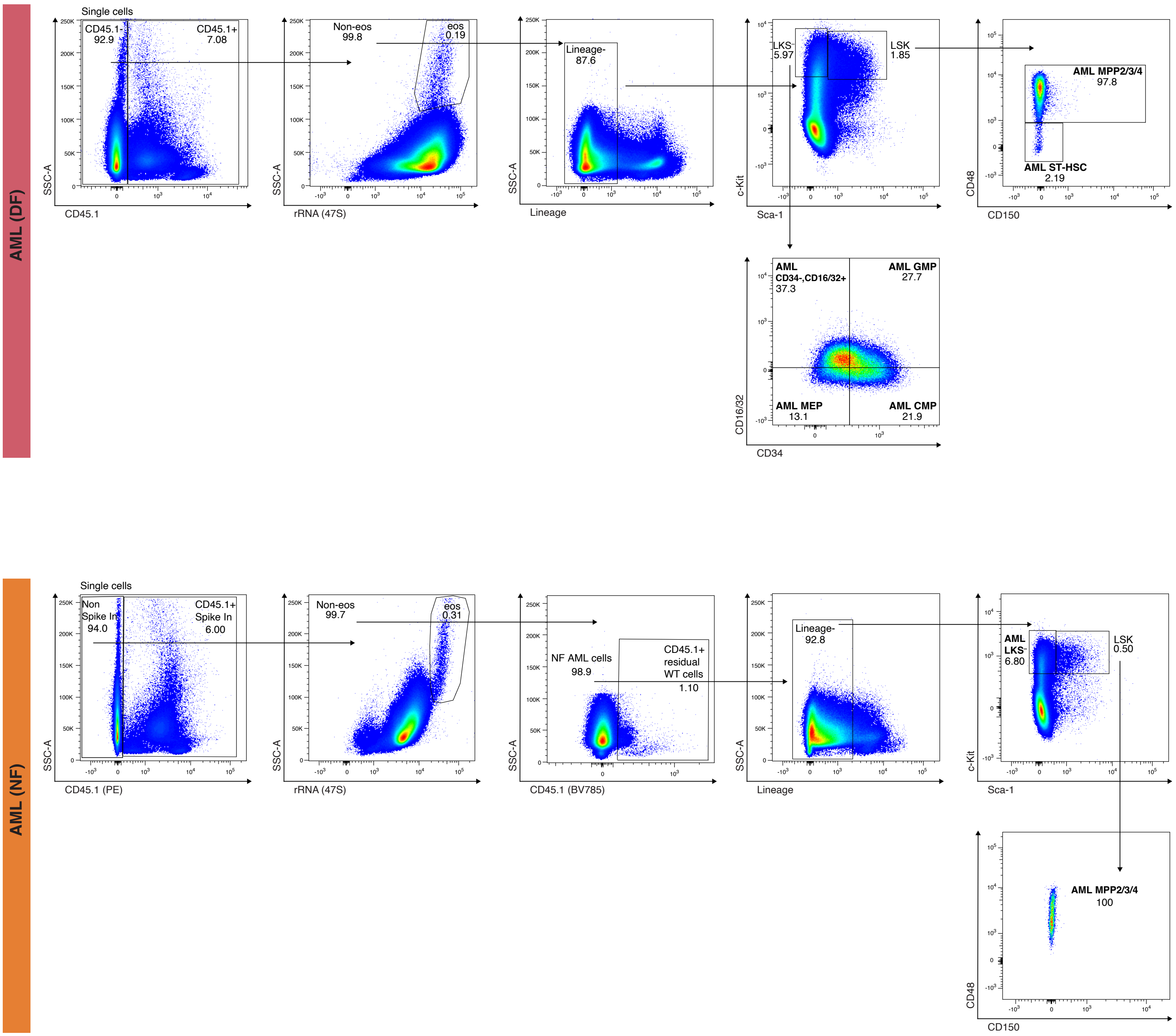

**Supplemental Figure S6. AML gating strategies (related to Figure 4)**

Representative flow cytometry plots depicting the gating strategies used for AML (DF) and AML (NF) rRNA FISH-Flow analyses. Gates were guided by normal BM profiles and unstained fluorescence controls. Final populations used for analysis and plotting are written in bold text. Numbers underneath population names indicate the percentage contribution to the represented plot. Full written gating schemes are listed in Supplemental Table S1. Of note, these AML models do not have any cells falling within LT-HSC/HSC/MPP1 gates, and the AML (NF) model does not have any cells found within the ST-HSC gate. While the AML (NF) model has abundant cells within the LKS$^{-}$ gate, they did not form distinct CMP, GMP, or MEP populations.

#### Supplemental Figure S7

**A**

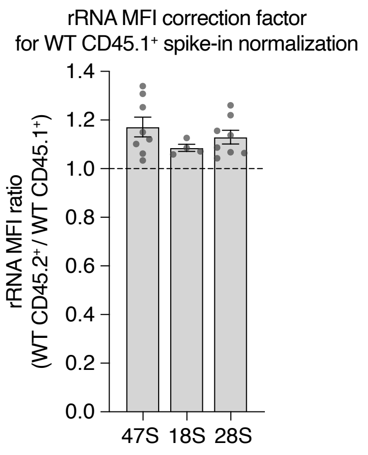


**B**

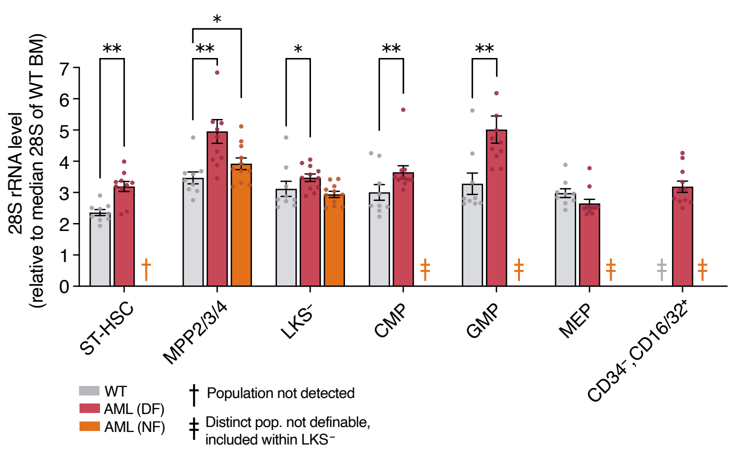


**C**

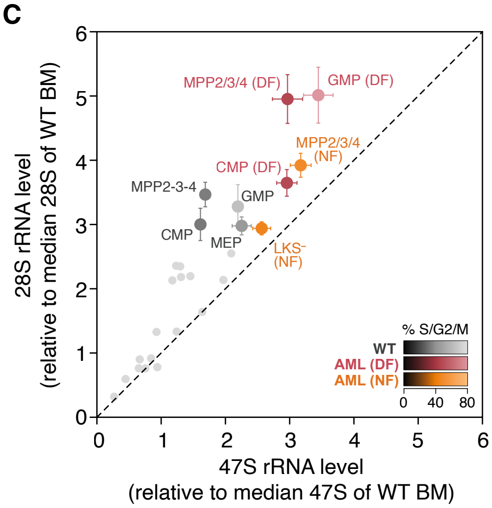


**D**

### % S/G2/M

| | WT | AML (DF) | AML (NF) |
|---|---|---|---|
| ST-HSC | 20.2 | 46.3 | |
| MPP2/3/4 | 34.3 | 47.1 | 49.1 |
| LKS⁻ | 46.0 | 48.9 | 45.1 |
| CMP | 33.3 | 48.5 | |
| GMP | 63.6 | 80.0 | |
| MEP | 43.4 | 32.4 | |
| CD34⁻,CD16/32⁺ | | 52.9 | |

**E**

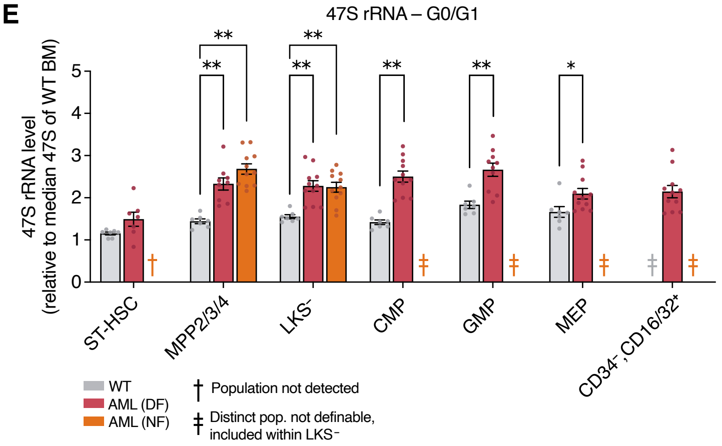


**F**

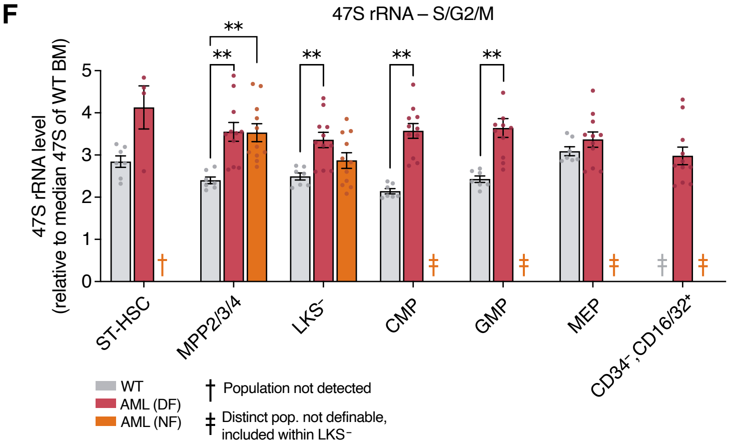


**G**

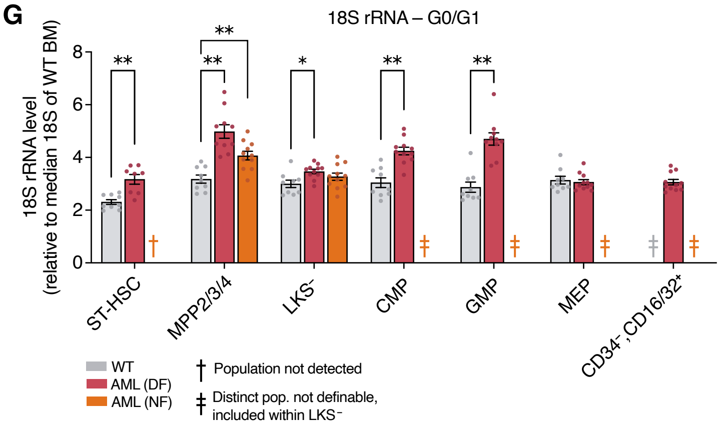


**H**

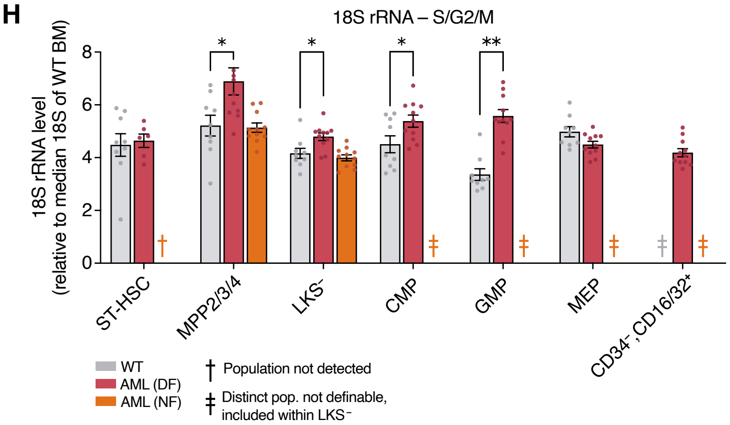

**Supplemental Figure S7. rRNA dynamics in AML progenitor-like cells (related to Figure 4)**

**(A)** Ratio of total BM $CD45^+$ rRNA MFI in wild-type CD45.2 mice (used for normal BM FISH-Flow) versus wild-type CD45.1 mice (used as spike-in for AML FISH-Flow). Ratios were applied as correction factors to the spike-in-normalized AML rRNA MFIs by dividing each normalized MFI by the corresponding correction factor.

**(B)** Relative levels of 28S rRNA in AML stem and progenitor-like cells compared to matched normal counterparts. All levels normalized to the median rRNA levels of total WT BM. **†** indicates populations that were not detected, and **‡** indicates populations that were not separately definable (did not form distinct populations on flow cytometry) and are instead included within the $LKS^-$ population.

**(C)** Comparison of relative nascent rRNA (47S rRNA) and ribosome subunit abundance (28S rRNA) across AML populations and normal counterparts (normalized to the median rRNA levels of total WT BM). The color of each cell type represents the percentage of cells in S/G2/M cell cycle phases.

**(D)** Table of the average percentage of cells in S/G2/M phases in all leukemic populations included in this study and their normal counterpart populations.

**(E-F)** Relative levels of 47S rRNA in AML stem and progenitor-like cells in G0/G1 (E) and S/G2/M (F) compared to matched normal counterparts. All levels normalized to the median 47S rRNA level of total WT BM (total cells).

**(G-H)** Same as E-F, showing 18S rRNA.

In some cases, replicates outside of the axis limits are not shown ( ≤ 2 replicates per cell type). All bar graphs and scatter plots show mean ± SEM. Comparisons by two-sided Mann-Whitney tests with FDR correction (Benjamini–Hochberg). ns ($p \geq 0.05$), * ($p < 0.05$), ** ($p < 0.01$), *** ($p < 0.001$), **** ($p < 0.0001$).

#### Supplemental Figure S8

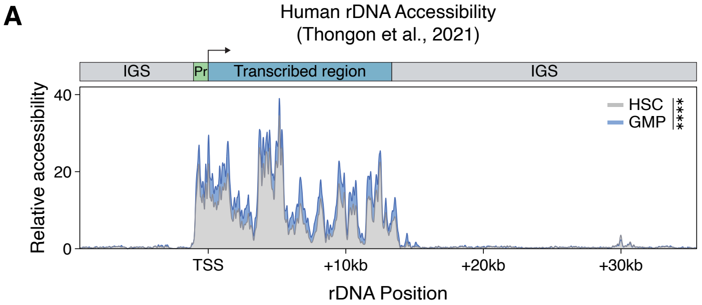


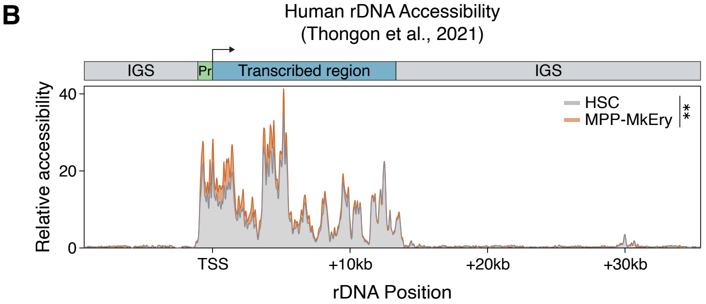


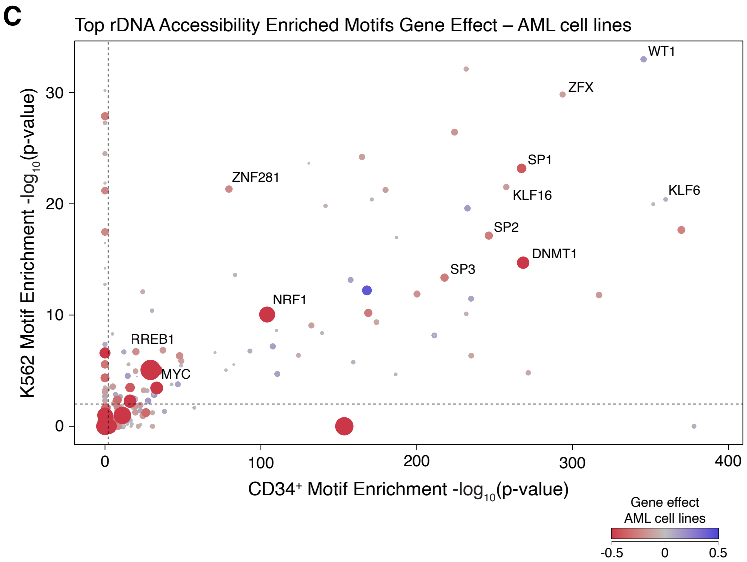


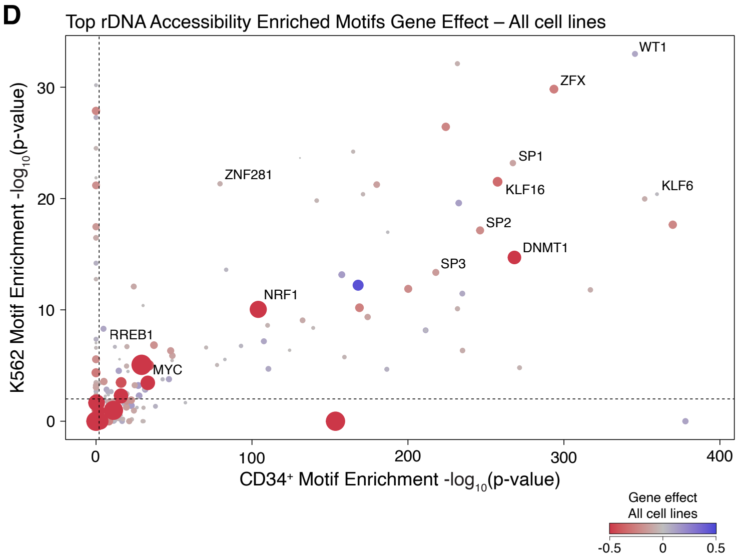

**Supplemental Figure S8. rDNA accessibility trends across hematopoietic cell types (related to Figure 6)**

**(A-B)** Representative tracks depicting accessibility along the rDNA transcribed region and intergenic spacer (IGS) in human myeloid (A) and erythroid (B) progenitors. The top schematic indicates the location of the rDNA promoter, transcribed region, and IGS. Signal at each position represents composite accessibility across all rDNA repeats from all cells in the assayed population. Comparisons by two-sided Mann-Whitney tests with FDR correction (Benjamini–Hochberg).

**(C-D)** CRISPR dependency gene effect scores in AML cell lines (C) and all cancer cell lines (D) for TFs enriched in Figure 6H-I. More negative scores indicate more severe dependency. Point size and color denote the gene effect score (scores capped at -0.5 and 0.5). Axes show $-\log_{10}$(FDR) from two-sided Mann-Whitney tests with FDR correction (Benjamini–Hochberg). Dashed lines indicate a TF motif enrichment significance threshold of $p < 0.01$. Scores represented in these plots were used to calculate the (AML minus all cancer) gene effect difference plotted in Figure 6J.

#### Supplemental Figure S9

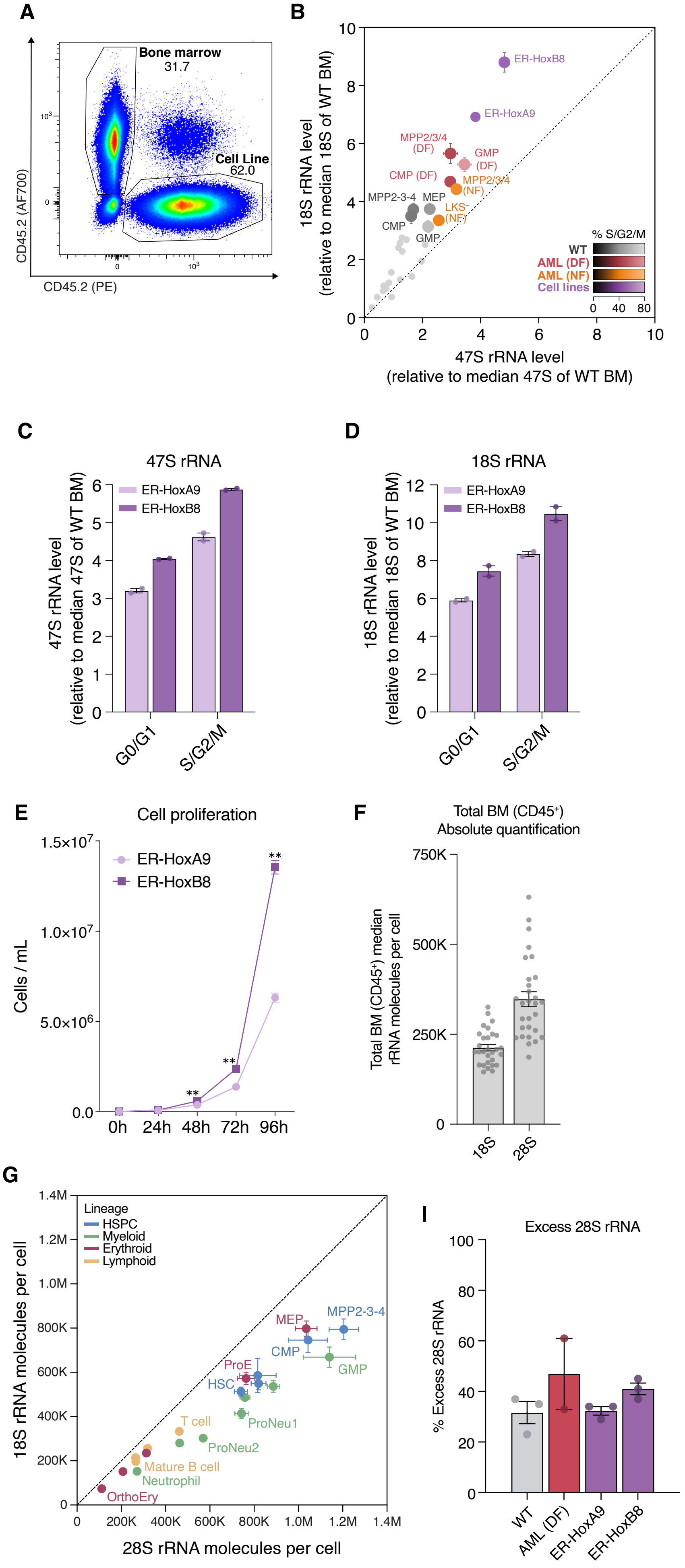


**H**

| | Cell type | 18S rRNA molecules per cell | 28S rRNA molecules per cell | Excess 28S rRNA molecules (% excess) |
|---|---|---|---|---|
| **WT** | HSC | 513K ± 20.2K | 740K ± 29.8K | 227K (45%) |
| | MPP1 | 586K ± 77.0K | 816K ± 82.9K | 230K (45%) |
| | ST-HSC | 549K ± 28.5K | 820K ± 33.5K | 271K (52%) |
| | MPP2/3/4 | 794K ± 46.9K | 1.20M ± 66.3K | 411K (53%) |
| | CMP | 745K ± 55.4K | 1.04M ± 87.3K | 298K (40%) |
| | GMP | 668K ± 45.8K | 1.14M ± 118.4K | 472K (68%) |
| | ProNeu1 | 413K ± 22.7K | 743K ± 30.0K | 329K (81%) |
| | ProNeu2 | 302K ± 10.2K | 569K ± 19.0K | 267K (89%) |
| | Neutrophil | 152K ± 3.83K | 271K ± 5.40K | 119K (78%) |
| | cMoP | 536K ± 26.2K | 886K ± 27.0K | 350K (67%) |
| | Classical Monocyte | 486K ± 9.60K | 758K ± 22.3K | 272K (56%) |
| | Intermediate Monocyte | 280K ± 5.96K | 463K ± 9.81K | 183K (66%) |
| | MEP | 797K ± 34.6K | 1.04M ± 48.5K | 238K (30%) |
| | ProE | 572K ± 28.0K | 763K ± 39.3K | 191K (34%) |
| | BasoE | 235K ± 9.16K | 313K ± 6.60K | 78.5K (34%) |
| | PolyE | 151K ± 7.51K | 207K ± 6.58 | 55.9K (38%) |
| | OrthoE | 73.5K ± 3.6K | 111K ± 3.9K | 37.7K (52%) |
| | PreProB/ProB/PreB | 256K ± 11.3K | 319K ± 17.0K | 62.1K (24%) |
| | Immature B | 214K ± 14.1K | 264K ± 15.3K | 50.1K (24%) |
| | Mature B | 197K ± 9.33K | 265K ± 10.0K | 68.4K (35%) |
| | T cell | 333K ± 16.5K | 461K ± 16.5K | 129K (39%) |
| **AML (DF)** | ST-HSC | 783K ± 45.1K | 1.11M ± 55.9K | 327K (43%) |
| | MPP2/3/4 | 1.20M ± 72.5K | 1.72M ± 131.9K | 516K (42%) |
| | CMP | 998K ± 39.8K | 1.27M ± 71.9K | 269K (27%) |
| | GMP | 1.12M ± 51.7K | 1.74M ± 151.1K | 618K (54%) |
| | MEP | 722K ± 27.9K | 920K ± 46.4K | 198K (27%) |
| | CD34$^-$,CD16/32$^+$ | 752K ± 33.4K | 1.11M ± 62.2K | 355K (47%) |
| **AML (NF)** | MPP2/3/4 | 943K ± 34.1K | 1.36M ± 64.9K | 420K (44%) |
| | LKS$^-$ | 714K ± 21.2K | 1.02M ± 35.2K | 307K (43%) |
| **Cell lines** | ER-HoxA9 | 1.47M ± 22.2K | 1.94M ± 98.5K | 469K (32%) |
| | ER-HoxB8 | 1.84M ± 72.5K | 2.59M ± 2.73K | 717K (38%) |

**Supplemental Figure S9. Myeloid cell lines enable absolute quantification of 18S and 28S rRNA molecules per cell (related to Figure 7)**

**(A)** Gating strategy used to identify cell line and WT bone marrow spike-in populations.

**(B)** Comparison of relative nascent rRNA (47S rRNA) and ribosome subunit abundance (18S rRNA) across all contexts (normalized to the median rRNA levels of total WT BM). The color of each cell type represents the percentage of cells in S/G2/M cell cycle phases.

**(C-D)** Relative levels of 47S rRNA (C) and 18S rRNA (D) in ER-HoxA9 and ER-HoxB8 cell lines in G0/G1 and S/G2/M cell cycle phases. All levels normalized to the median level of each respective rRNA in total WT BM (total cells).

**(E)** Proliferation curve of ER-HoxA9 and ER-HoxB8 cells. Comparisons at each timepoint by unpaired, two-sided Mann-Whitney test.

**(F)** Average absolute 18S and 28S rRNA molecules per cell calculated for the total mouse BM $CD45^+$ population using multiple biological and technical replicates of total RNA quantification for ER-HoxA9 and ER-HoxB8 cell lines, combined with relative MFI measurement of each cell line compared to spike-in WT bone marrow. Average values were used to calculate absolute mature rRNA molecules per cell for all populations included in this study based on relative rRNA FISH-Flow measurements.

**(G)** Comparison of the average number of 28S and 18S rRNA molecules per cell across all normal hematopoietic cell types. Values represent the average number of rRNA molecules per cell across all replicates per cell type. Cell types are colored by lineage.

**(H)** Table summarizing the average mature rRNA molecules per cell (range indicates ± SEM) and excess 28S rRNA molecules (number of molecules and percent excess) across all cell types included in this study.

**(I)** Excess 28S rRNA molecules per cell compared to 18S from total RNA samples derived from wild type and AML (DF) total bone marrow and from ER-HoxA9 and ER-HoxB8 cell lines. Each point represents the comparison of 28S and 18S rRNA abundance in paired samples from the same biological replicate. n = 2-3 replicates.

All bar graphs and scatter plots show mean ± SEM.

#### Supplemental Figure S10

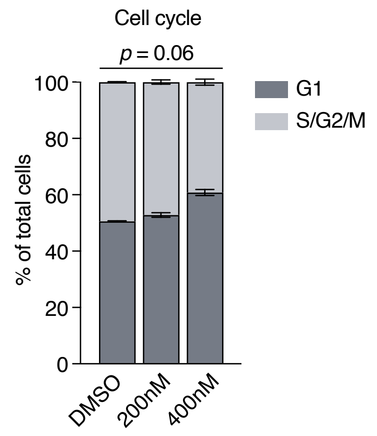


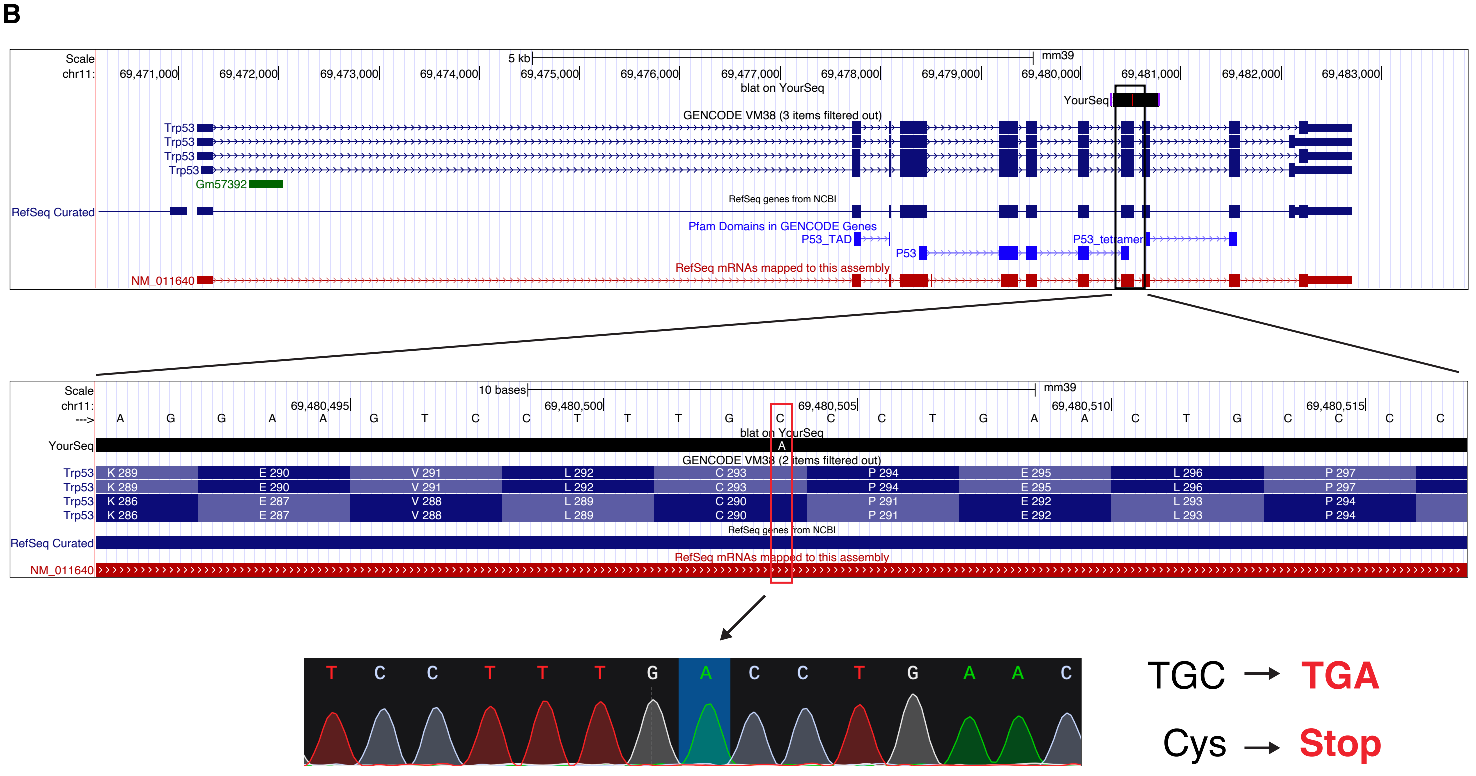


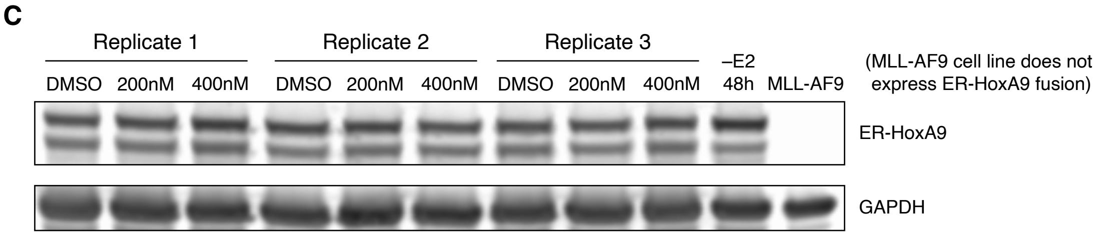

**Supplemental Figure S10. Downstream effects of Pol I degradation occur without functional p53 or changes in ER-HoxA9 fusion expression (related to Figure 8)**

**(A)** Proportion of Pol I degron cells in G1 or S/G2/M phases of the cell cycle after culturing with DMSO or the indicated doses of dTAG for 48 hours. n = 3 replicates, comparison by one-way ANOVA.

**(B)** Sanger sequencing chromatogram and UCSC Genome Browser visualization of the Cys → Stop mutation (C-to-A mutation) in *Trp53* in ER-HoxA9 Pol I degron cells. The ploidy of the *Trp53* allele in this cell line is unknown.

**(C)** Western blot showing unchanged expression of the ER-HoxA9 transgene fusion protein in Pol I degron cells treated with DMSO or the indicated doses of dTAG for 72 hours. The MLL-AF9 cell line does not contain the ER-HoxA9 transgene and therefore does not express the fusion protein. GAPDH is shown as a loading control. Samples lysed and loaded according to equal cell count.
