## Supplemental Table 1 for "Dynamics of Ribosomal RNA Transcription and Abundance in Normal and Leukemic Hematopoiesis"

| | Cell type | Gating strategy |
|---|---|---|
| WT | HSC | CD45+, *Lineage-, Sca1+, cKit+, CD150+, CD48-, CD34- |
| | MPP1 | CD45+, *Lineage-, Sca1+, cKit+, CD150+, CD48-, CD34+ |
| | ST-HSC | CD45+, *Lineage-, Sca1+, cKit+, CD150-, CD48- |
| | MPP2/3/4 | CD45+, *Lineage-, Sca1+, cKit+, CD48+ |
| | CMP | CD45+, *Lineage-, Sca1-, cKit+, CD34+, CD16/32- |
| | GMP | CD45+, *Lineage-, Sca1-, cKit+, CD34+, CD16/32+ |
| | ProNeu1 | CD45+, **Lineage-, cKit+, CD34+, CD16/32+, Ly6C+, CD115-, CD81+, CD106- |
| | ProNeu2 | CD45+, **Lineage-, cKit+, CD34+, CD16/32+, Ly6C+, CD115-, CD81+, CD106+ |
| | Neutrophil | CD45+, CD11b+, Ly6G+ |
| | cMoP | CD45+, **Lineage-, cKit+, CD34+, CD16/32+, Ly6C+, CD115+, CD81- |
| | Classical Monocyte | CD45+, CD11b+, Ly6G-, Ly6Chigh |
| | Intermediate Monocyte | CD45+, CD11b+, Ly6G-, Ly6Cmid |
| | MEP | CD45+, Lineage-, Sca1-, cKit+, CD34-, CD16/32- |
| | ProE | CD45-, Ter119+, Ter119low, CD44+ |
| | BasoE | CD45-, Ter119+, Ter119high, CD44+, FSC-Ahigh, CD44high |
| | PolyE | CD45-, Ter119+, Ter119high, CD44+, FSC-Amid, CD44mid |
| | OrthoE | CD45-, Ter119+, Ter119high, CD44+, FSC-Alow, CD44low |
| | PreProB/ProB/PreB | CD45+, IgM-, B220mid |
| | Immature B | CD45+, IgM+, B220mid |
| | Mature B | CD45+, IgM+, B220high |
| | T cell | CD45+, CD19-, CD3+ |
| AML (DF) | AML ST-HSC | *Lineage-, Sca1+, cKit+, CD150-, CD48- |
| | AML MPP2/3/4 | *Lineage-, Sca1+, cKit+, CD48+ |
| | AML CMP | *Lineage-, Sca1-, cKit+, CD34+, CD16/32- |
| | AML GMP | *Lineage-, Sca1-, cKit+, CD34+, CD16/32+ |
| | AML MEP | *Lineage-, Sca1-, cKit+, CD34-, CD16/32- |
| | AML CD34-,CD16/32+ | *Lineage-, Sca1-, cKit+, CD34-, CD16/32+ |
| AML (NF) | AML LKS- | *Lineage-, Sca1-, cKit+ |
| | AML MPP2/3/4 | *Lineage-, Sca1+, cKit+, CD48+ |

All cells are DAPI+, live cells, single cells

* Lineage markers: Ter119, B220, CD19, CD3, CD49b, Gr-1, CD11c

** Lineage markers: Sca1, NK1.1, CD90.2, B220, Ly6G
